# The Natural Products Discovery Center: Release of the First 8490 Sequenced Strains for Exploring Actinobacteria Biosynthetic Diversity

**DOI:** 10.1101/2023.12.14.571759

**Authors:** Edward Kalkreuter, Satria A. Kautsar, Dong Yang, Chantal D. Bader, Christiana N. Teijaro, Lucas L. Fluegel, Christina M. Davis, Johnathon R. Simpson, Lukas Lauterbach, Andrew D. Steele, Chun Gui, Song Meng, Gengnan Li, Konrad Viehrig, Fei Ye, Ping Su, Alexander F. Kiefer, Angela Nichols, Alexis J. Cepeda, Wei Yan, Boyi Fan, Yanlong Jiang, Ajeeth Adhikari, Cheng-Jian Zheng, Layla Schuster, Tyler M. Cowan, Michael J. Smanski, Marc G. Chevrette, Luiz P. S. de Carvalho, Ben Shen

## Abstract

Actinobacteria, the bacterial phylum most renowned for natural product discovery, has been established as a valuable source for drug discovery and biotechnology but is underrepresented within accessible genome and strain collections. Herein, we introduce the Natural Products Discovery Center (NPDC), featuring 122,449 strains assembled over eight decades, the genomes of the first 8490 NPDC strains (7142 Actinobacteria), and the online NPDC Portal making both strains and genomes publicly available. A comparative survey of RefSeq and NPDC Actinobacteria highlights the taxonomic and biosynthetic diversity within the NPDC collection, including three new genera, hundreds of new species, and ∼7000 new gene cluster families. Selected examples demonstrate how the NPDC Portal’s strain metadata, genomes, and biosynthetic gene clusters can be leveraged using genome mining approaches. Our findings underscore the ongoing significance of Actinobacteria in natural product discovery, and the NPDC serves as an unparalleled resource for both Actinobacteria strains and genomes.

## Introduction

Natural products (NPs) have played a pivotal role in revolutionizing modern and traditional medicine, serving as the foundation for nearly half of all approved drugs available to combat infectious diseases, cancer, endocrine disorders, and various other ailments. ^1^ Throughout history, humanity has naively harnessed diverse sources of medicinal NPs, but it was not until the mid-20th century that the pursuit of these valuable compounds became a targeted effort, spurred by the groundbreaking discovery of streptomycin from the soil-dwelling Actinobacterium *Streptomyces griseus* in 1942. ^2, 3^ During this era of rapid discovery, many pharmaceutical companies leveraged their resources to search the globe for the next blockbuster drugs and their associated microbial producers. ^4^ These efforts resulted in the accumulation of vast microbial strain collections, which served as the global basis for drug discovery for decades. Despite the immense success of this approach, a confluence of factors, including high rediscovery rates, advancements in combinatorial chemistry and high-throughput screening, and economic considerations, prompted a shift away from industrial NP exploration and microbial strain collections. ^3, 5^ However, with the advent of the genomics era, it became increasingly evident that bacteria, especially prolific taxa such as Actinobacteria, possessed far more biosynthetic diversity than had been predicted prior to genome sequencing. ^6^

Actinobacteria, recently renamed as Actinomycetota, ^7^ are found ubiquitously, spanning environments from soil and plants (e.g., *Streptomyces* and *Micromonospora*) to humans (e.g., *Corynebacterium* and *Mycobacterium*). ^8, 9, 10^ The expansive secondary metabolomes of most non-pathogenic Actinobacteria enable them to rapidly respond to environmental changes and occupy unique ecologies, in part due to highly diverse chemistry and fine-tuned regulation. ^11^ Consequently, Actinobacteria have emerged as the most extensively studied bacterial phylum for NP discovery, accounting for more than 60% of all characterized bacterial NPs (Fig. 1A), with its largest genus, *Streptomyces*, particularly standing out. ^12^ Genomic analyses have consistently indicated rich biosynthetic gene cluster (BGC) diversity in many Actinobacteria genera that matches their central role in NP exploration. ^6^ However, these bacteria remain disproportionately underrepresented in accessible genome databases such as NCBI’s RefSeq relative to their prevalence in NP databases (Fig. 1B). ^13, 14^ A recent study of 824 diverse Actinobacteria genomes has predicted that as little as 30-50% of the projected Actinobacteria phylogenetic diversity is represented in public genomic datasets, indicating that much work is remaining for the community. ^15^ Furthermore, most public Actinobacteria genomes belong to a few health- or disease-associated species such as the causative agent of tuberculosis, *Mycobacterium tuberculosis* (Fig. 1B), resulting in the available Actinobacteria genomes harboring only a small fraction of the potential BGC and NP diversity of this phylum. ^14^

**Figure 1.**
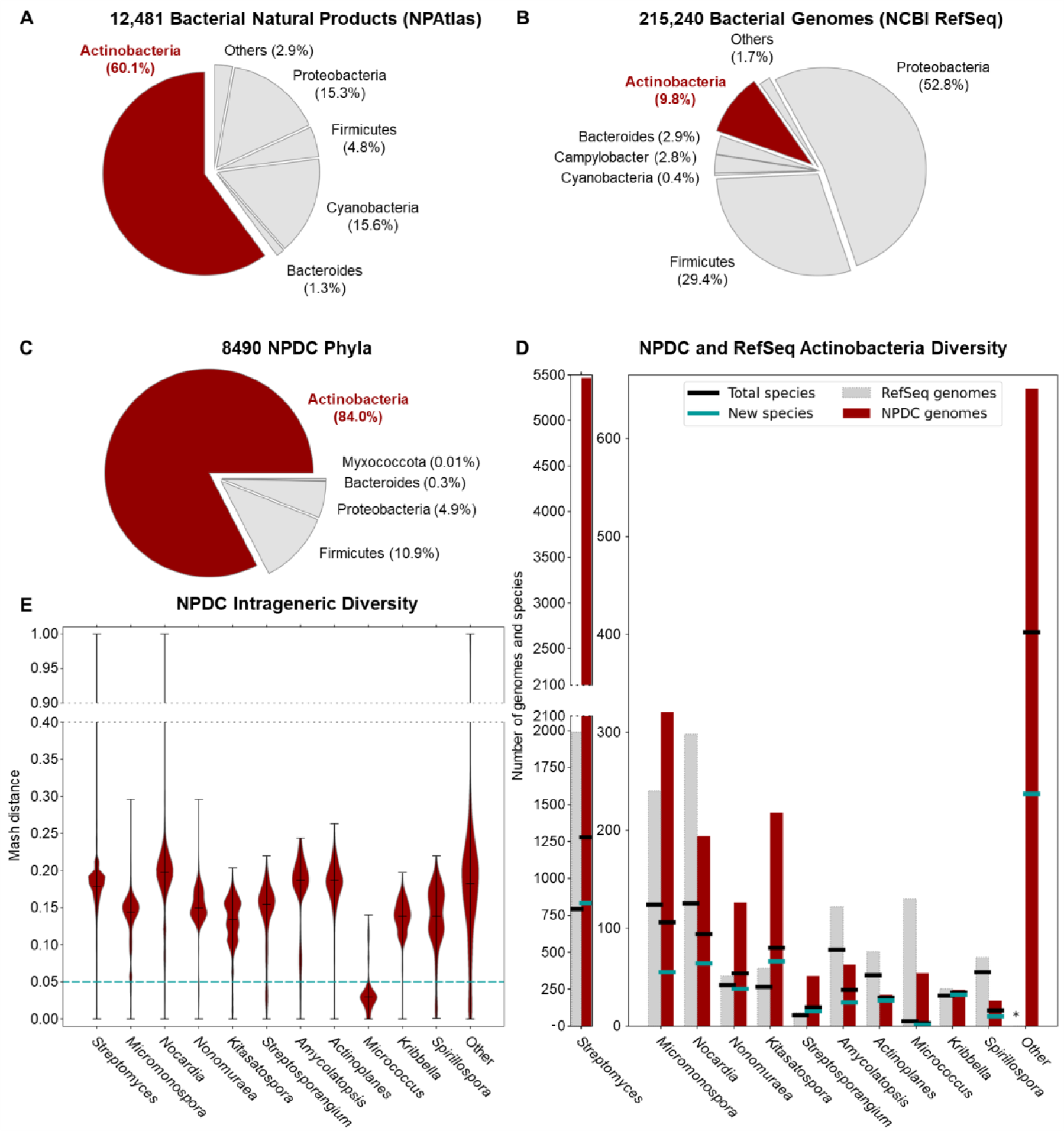
Natural product-producing Actinobacteria are well-represented among NPDC strains. (A) NPs derived from Actinobacteria (red) are overrepresented in comparison to the NPs isolated from other bacterial sources according to NPAtlas. ^13^ Actinobacteria NPs are primarily isolated from *Streptomyces* (73%), *Micromonospora* (4%), and *Actinomadura* (2%). (B) Actinobacteria genomes (red) are underrepresented in comparison to other bacterial genomes available in NCBI RefSeq. ^16^ Actinobacteria RefSeq genomes come primarily from pathogens or other health-associated bacteria such as *Mycobacterium* (45%), *Bifidobacterium* (7%), or *Corynebacterium* (6%). *Streptomyces* make up only a small fraction of Actinobacteria RefSeq genomes (9%). (C) Breakdown of NPDC genomes by phylum, confirming that most NPDC strains are Actinobacteria. (D) Actinobacteria genomes from representative genera in the NPDC (red bars) are compared to genomes from RefSeq (grey bars). Total numbers of species in a genus, based on a 0.05 Mash distance cut-off, are indicated by black lines. Numbers of new species in the NPDC, defined as lacking a closely related representative in the Genome Taxonomy Database (GTDB), are indicated by teal lines. The genomes from non-highlighted genera are combined into a single ‘Other’ category here, and the breakdowns by individual genera are depicted in Supplementary Fig. 8. (E) Intrageneric diversity in representative, well-populated NPDC genera are displayed by Mash distance distribution. Mash distances smaller than 0.05 are treated as the same species.

The available Actinobacteria genomes highlight their immense potential for NP discovery, but translation of their genomic potential into isolated NPs remains a resource- and time-intensive endeavor. ^6^ However, the scale and speed of genome sequencing can help streamline this process by enabling the rapid categorization of BGCs. By identifying groups of related BGCs, known as gene cluster families (GCFs), redundant efforts can be avoided and the most promising strains among thousands can be strategically prioritized for fermentation. ^17, 18^ To democratize the NP discovery process, these strains and their genomes must be made readily available, which serves as the inspiration behind the Natural Products Discovery Center (NPDC) at the Herbert Wertheim UF Scripps Institute for Biomedical Innovation & Technology (UF Scripps). Comprised primarily of strains collected by industry between the 1940s and the 2010s, the NPDC collection encompasses a total of 122,449 strains, with more than half morphologically assigned as Actinobacteria. ^5^ Notably, at the time of the NPDC launch, these Actinobacteria outnumber the collective total Actinobacteria deposited in the world’s largest public strain collections, including ATCC (3564), NBRC (3266), NRRL (1863), and DSMZ (7223), by several fold (Supplementary Fig. 1). ^19, 20, 21, 22^

To develop this unique collection into a community resource for NP discovery and other user-driven needs such as new biocatalytic, biotechnological, or microbiological applications, the NPDC strains are being continuously grown and their genomes sequenced. Herein, we present a comprehensive analysis of the taxonomic diversity and biosynthetic potential of the initial 8490 sequenced strains. Moreover, we introduce an online portal that facilitates the exploration of their genomes, particularly their biosynthetic potential for NP production, and access to the physical strains themselves. We further utilize the NPDC Portal tools to identify the bonnevillamides as narrow-spectrum antibiotics and validate the esperamicin BGC in multiple alternative producers.

## Results

### The NPDC collection greatly expands the known taxonomic diversity of Actinobacteria

Genome sequencing facilitates accurate taxonomic classification of bacterial strains and the unveiling of evolutionary relationships between microorganisms. Therefore, a subset of the NPDC collection, 8490 bacteria making up 6.9% of the whole collection and expected to be representative of the ecologic, chronologic, and geographic diversity of the NPDC collection, was subjected to short-read genome sequencing (Supplementary Fig. 2-4 and Supplementary Table 1). ^5^ All of the strains referenced in the manuscript are part of the UF Scripps NPDC bacterial strain collection, and all of these bacterial strains fall within the scope of the Nagoya Protocol, which entered into force on October 12, 2014. ^23^ The median genome in the dataset has a size of 8.1 Mb with 76 contigs and an N50 of 245 kb, making their quality compare favorably to most RefSeq genomes (Supplementary Fig. 5-7). ^16^ The genome-based taxonomy of the 8940 bacteria was determined by GTDB-Tk-v2, revealing 7142 (84.1%) as Actinobacteria and the remainder as primarily Firmicutes and Proteobacteria (Fig. 1C and Supplementary Table 2). ^24^ Although unclear if the non-Actinobacteria strains were the original isolation targets or potential contaminations introduced during various stages of sample collection or handling, this ratio can be extrapolated to indicate the presence of more than 102,000 Actinobacteria within the NPDC collection. Notably, of the original 7055 Actinobacteria phenotypic assignments, 6247 (88.5%) were correctly assigned as Actinobacteria; however, this historical accuracy varies greatly when examined at the genus level (Extended Data Fig. 1 and Supplementary Table 3). While historical classifications for well-studied genera like *Streptomyces* (91.9%) or *Micromonospora* (78.0%) were comparatively accurate, genomic assignments for rarer genera such as *Nocardia* (18.6%), *Streptosporangium* (20.0%), *Actinoplanes* (29.1%), and *Actinomadura* (0.0%) display notable inconsistencies with their historical designations (Extended Data Fig. 1 and Supplementary Table 3).

Upon the public launch of the NPDC in late 2022, the 7142 NPDC Actinobacteria genomes expanded the total RefSeq Actinobacteria genomes by more than 33% and demonstrated the expansive taxonomic diversity encompassed within. These genomes are distributed among 95 Actinobacteria genera (in addition to 1348 NPDC genomes from 104 non-Actinobacteria genera), six of which comprise more than 100 NPDC genomes each (Fig. 1D and Supplementary Fig. 8). The polyphyletic genus *Streptomyces*, considered the largest bacterial genus by species count, has the largest number of NPDC genomes (5386), greatly surpassing the number of RefSeq *Streptomyces* (1990) and underscoring the NPDC’s focus on NP-producing genera (Fig. 1D). ^25^ Of these 95 genera, only 19 have 20 or more characterized NPs in the NPAtlas database. ^13^ Additionally, ten smaller and rarer genera including *Micromonospora*, *Kitasatospora*, *Nonomuraea*, and *Streptosporangium* minimally doubled their respective numbers of RefSeq genomes (Fig. 1D and Supplementary Fig. 8). All other genera highlighted in Fig. 1C-E were expanded by at least 40%. The release of these genomes has also bolstered species coverage across Actinobacteria, with the number of newly sequenced species at least doubling the number of RefSeq species in ten genera, including *Streptomyces*, *Kitasatospora*, *Streptosporangium*, and *Kribbella* (Fig. 1D and Supplementary Fig. 8). In total, more than 65% of all 2152 Actinobacteria species present in the NPDC collection were not previously present in the RefSeq dataset, though RefSeq species are more represented (6.2 strains/species) than new species (1.8 strains/species) among NDPC strains (Supplementary Fig. 9).

The NPDC taxonomic novelty is primarily found in groups of related clades spread throughout the Actinobacteria families best known for NP production (Fig. 2). The observed strain distribution suggests that the NPDC not only contains new species and subspecies but also addresses these preexisting phylogenetic gaps with multiple representatives of related strains. This distribution pattern is particularly evident in *Streptomyces*, wherein most clades are enriched by new NPDC species except for a few clades already covered deeply by RefSeq genomes (Fig. 2). However, some genera, especially those related to health and pathogenicity such as *Mycobacterium*, *Bifidobacterium*, and *Corynebacterium*, are notably underrepresented in the NPDC genomes.

**Figure 2.**
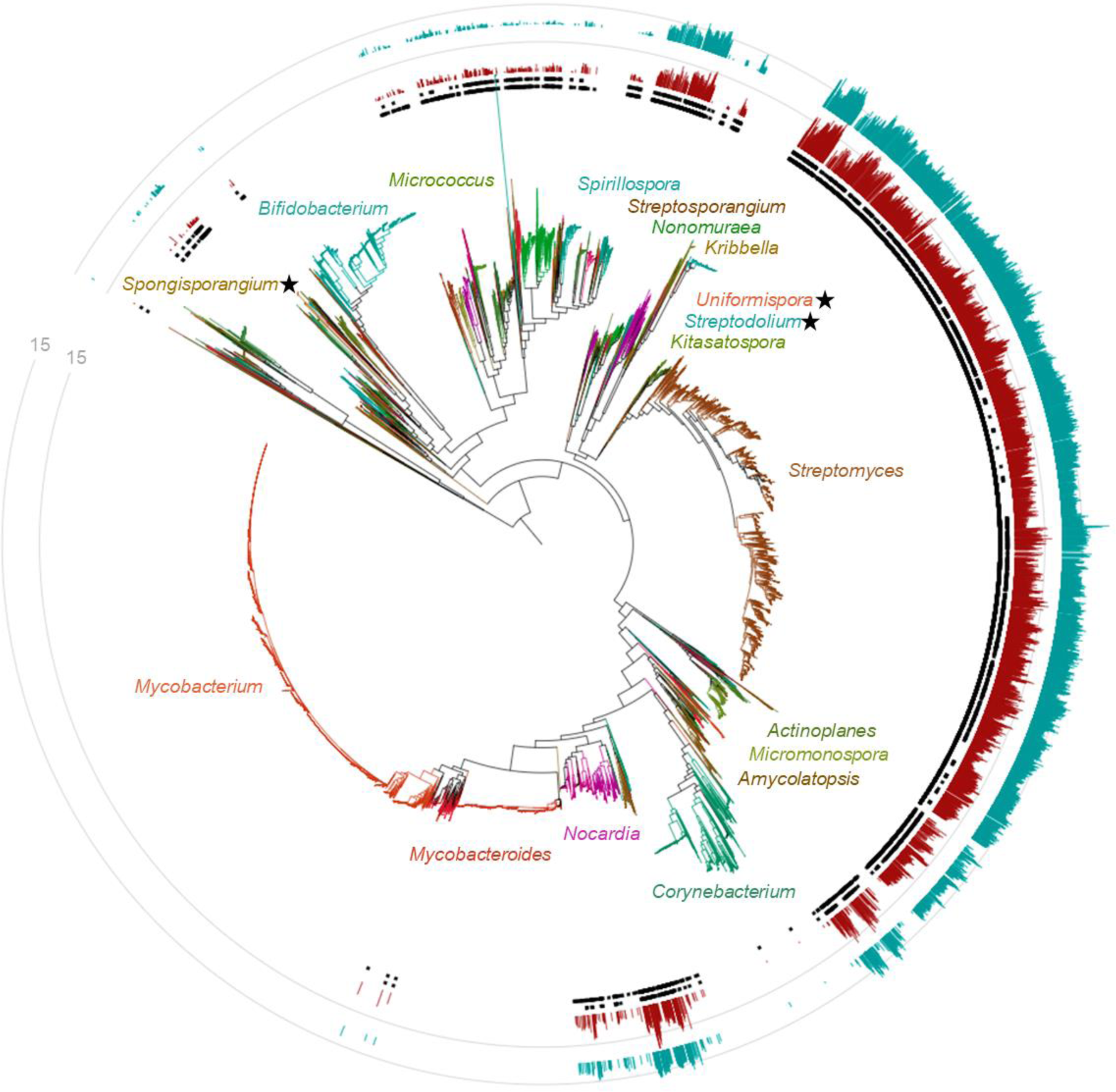
Taxonomic distribution of NPDC Actinobacteria strains and GCFs. The genome-based maximum likelihood phylogenetic tree of RefSeq and NPDC Actinobacteria is color-coded by genus, with representative genera labeled. The inner circle of black squares represents NPDC strains. The outer circle of black squares represents new species. The inner red bar chart indicates the number of new GCFs. The outer teal bar chart indicates the number of GCFs that overlap with RefSeq GCFs. New genera identified in this study are indicated with stars.

For most Actinobacteria genera, the NPDC genomes display high intrageneric diversity, as evidenced by their first quartile Mash distance exceeding 0.05, indicative of distinct species (Supplementary Fig. 10). ^26^ Mash distance correlates strongly with average nucleotide identity (ANI), especially in the ANI range of 90 to 100% (0.1 to 0 Mash distances, respectively). ^26^ Therefore, the high Mash distances observed for most NPDC Actinobacteria mirror their species-level diversity, especially in genera such as *Nocardia* and *Amycolatopsis* (Fig. 1E). Notably, a few rare genera such as *Micrococcus* are exceptions among Actinobacteria, with most Mash distances below this cutoff, consistent with previous findings of *M. luteus* as the single dominant species within this genus. ^27, 28^ In contrast, Mash distance analysis of *Mycobacterium* RefSeq genomes shows a clear multimodal distribution with the majority of genomes clustering around a Mash distance of 0 (Supplementary Fig. 11), reflecting the overrepresentation of *M. tuberculosis* and other pathogens.

Within the NPDC Actinobacteria, three putative new genera were identified based on GTDB-Tk-v2 and RED values. ^24, 29^ NPDC049639, herein designated as *Spongisporangium articulatum*, is the sole member of a new genus within the *Kineosporiaceae* family (RED value 0.829) and is most closely related to *Kineosporia* (Fig. 3A). However, while *Kineosporia* strains have an average genome size of 8.11 Mb, NPDC049639 has a significantly smaller 4.86 Mb genome encoding 11 BGCs. Further differentiating itself from other members of *Kineosporiaceae*, NPDC049639 forms large spherical structures during growth and shows no evidence of motile spores (Fig. 3B and Supplementary Fig. 12). The two other genera, herein named *Streptodolium* and *Uniformispora*, belong to the *Streptomycetaceae* family and are most closely related to *Yinghuangia* and *Embleya*, two genera reclassified from *Streptomyces* in 2018 (Fig. 3C). ^30^ The two strains of *Streptodolium elevatio*, NPDC002781 and NPDC048946 (RED values 0.843 and 0.844, respectively), share 98.2% genome similarity by ANI, suggesting two strains within a shared species (Supplementary Table 4). ^31^ NPDC002781 and NPDC048946 have 25 and 29 BGCs in their large 9.46 and 9.59 Mb genomes, respectively. By scanning electron microscopy (SEM), *S. elevatio* NPDC002781 displays long protruding aerial hyphae and barrel-shaped spores (Fig. 3D and Supplementary Fig. 13). In contrast, the two isolates of *Uniformispora flossi*, NPDC059210 and NPDC059280 (RED value 0.843), are likely the same strain (99.98% genome identity by ANI; Supplementary Table 4). ^31^ *U. flossi* NPDC59280 harbors 24 BGCs within its 8.80 Mb genome, and as indicated by its genus name, its aerial hyphae are long and uniformly segmented, except for nearly spherical terminal spores (Fig. 3E and Supplementary Fig. 14).

**Figure 3.**
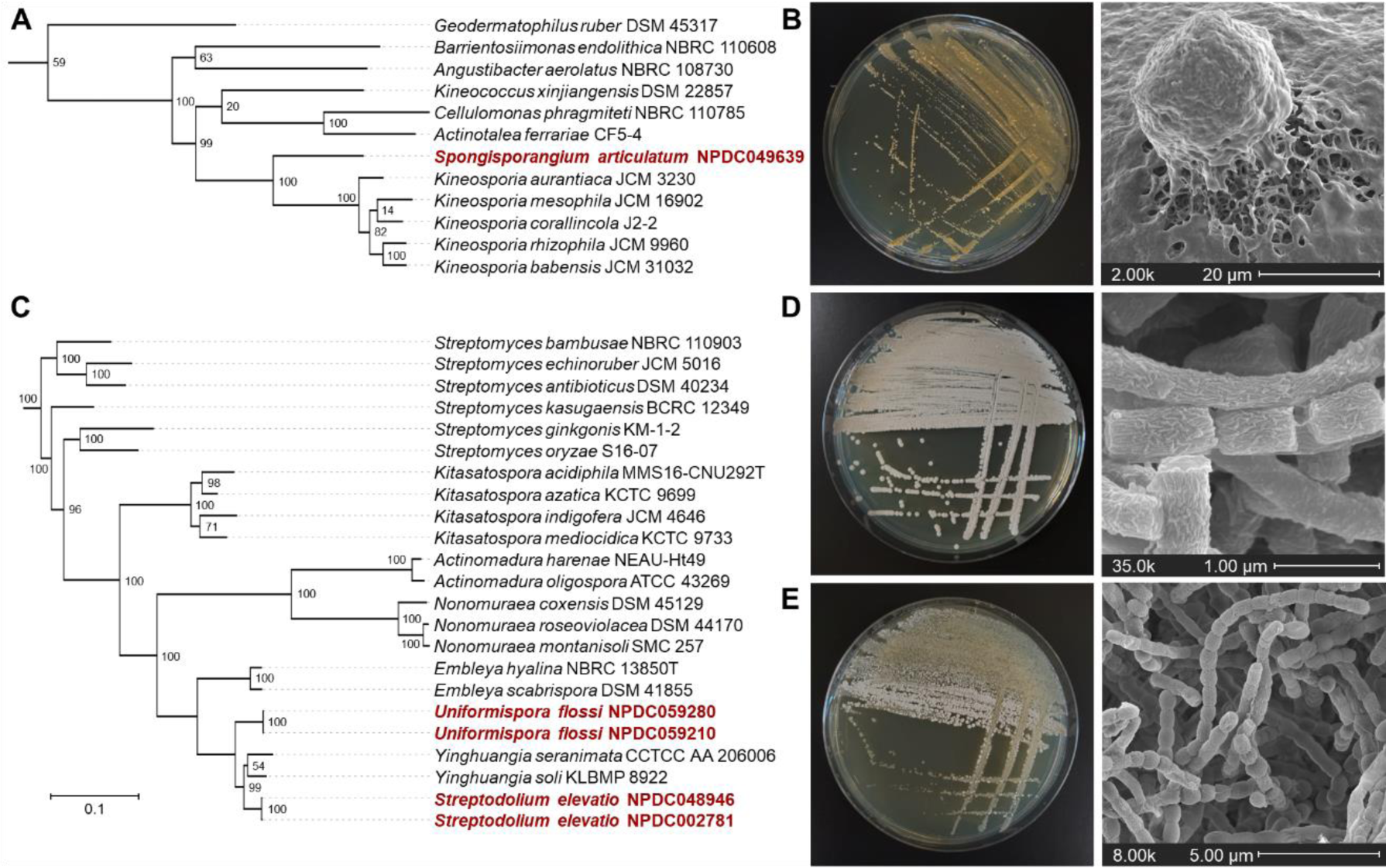
Proposed new genera identified in the NPDC. (A) Genome-based maximum likelihood phylogenetic tree showing the position of *Spongisporangium articulatum* NPDC049639 among representative type strains. (B) Representative macroscopic and SEM images of *S. articulatum* NPDC049639 highlighting the large spherical bodies on the spongelike culture surface inspiring the genus name. (C) Genome-based maximum likelihood phylogenetic tree showing the positions of *Streptodolium elevatio* NPDC002781 and NPDC048946 and *Uniformispora flossi* NPDC059210 and NPDC059280 among representative type strains. (D) Representative macroscopic and SEM images of *S. elevatio* NPDC002781 highlighting the fibrous sheath of the aerial hyphae. (E) Representative macroscopic and SEM images of *U. flossi* NPDC059280 highlighting the long chains of uniformly segmented aerial hyphae interconnected with thin filamentous strands. Additional SEM images are found in Supplementary Fig. 12-14.

New microbiology and taxonomic diversity can also be correlated with the biosynthetic diversity and potential of the strains of the NPDC. Actinobacteria typically allocate substantial portions of their genomes (up to 20%) to BGCs encoding NPs (Extended Data Fig. 2). However, not all genera are equal in their biosynthetic capacities, as might be expected for genera that reside in varying numbers of diverse or isolated ecologies (Supplementary Fig. 15). An earlier study has shown that host-associated Actinobacteria, though still prolific relative to other bacteria, utilize ∼20% less genomic capital for NP biosynthesis than their environmental counterparts. ^15, 32^ NPDC *Streptomyces* harbor more than 30 BGCs per genome, accounting for 11.7% of their total genomes on average, which is superior to most bacterial genera but falls below 23 other NPDC Actinobacteria genera (Fig. 2 and Extended Data Fig. 2). This finding is consistent with what has been observed in non-NPDC *Streptomyces* genomes. ^33^ Many understudied genera such as *Nocardia*, *Actinosynnema*, and the previously highlighted *Embleya*^34^ display exceptional biosynthetic potential, with *Embleya* averaging more than 50 BGCs across its ten NPDC genomes. The previous record for highest BGC dedication among Actinobacteria was held by *Kitasatospora kifunensis* DSM 41654 with 26.50% of its genome assigned to BGCs. ^15^ Within the NPDC, three strains have surpassed this threshold: *Kitasatospora* sp. NPDC008050 (30.32%), *Streptomyces olivoreticuli* NPDC020840 (28.17%), and *Streptomyces klenkii* NPDC049569 (26.51%). In contrast, other genera such as *Rothia*, *Micrococcus*, and *Agromyces* all harbor fewer than five BGCs per NPDC genome. Of the new genera, none of the 11 *Spongisporangium* BGCs share notable similarity to known BGCs available in the MIBiG database, with the most similar BGCs originating from strains in other *Kineosporiaceae* genera or less related genera such as *Mycobacterium*. *Streptodolium* and *Uniformispora*, in contrast, utilize more than 16% of their genomes for secondary metabolism (Extended Data Fig. 2), sharing more in common with the related *Embleya* than the type genus of the family, *Streptomyces*. Compared to *Streptodolium*, *Uniformispora* shares especially little biosynthetic similarity to other known strains. Overall, the NPDC Actinobacteria are heavily biased towards strains that dedicate significant genomic resources towards NP biosynthesis, especially in comparison to efforts that are purely taxonomically driven (Extended Data Fig. 3). ^15^

### NPDC Actinobacteria significantly expand the accessible biosynthetic potential

By collecting and sequencing strains with untapped and exceptional biosynthetic potential, more than 200,000 BGCs have already been identified in NPDC Actinobacteria genomes in this study. Extrapolating this number to encompass the entire NPDC collection based on current sequencing trends suggests the presence of nearly 3.25 million BGCs in total. Grouping of BGCs into GCFs allows for mapping of gene clusters with similar architectures often encoding related molecules and facilitates dereplication of BGCs with known NPs. ^35^ Additionally, while most NPDC BGCs are predicted by antiSMASH to be intact (Extended Data Fig. 4A), GCFs have the advantage of grouping the remaining fragmented BGCs together, enabling identification of otherwise missed producers of a desired NP type (Extended Data Fig. 4B).

Comparison between RefSeq and NPDC Actinobacteria GCFs shows that the rate of GCF discovery directly correlates to the increase in Actinobacteria genomes. A total of 20,112 GCFs were extracted from the 21,186 RefSeq Actinobacteria, and the 7142 NPDC Actinobacteria harbor 16,902 GCFs. With more than 40% of the identified GCFs being unique to the NPDC, release of these genomes results in ∼33% increases of both publicly available genomes and GCFs (Fig. 4A). More than 85% of the NPDC GCFs are found exclusively in only a single genus, including most of the GCFs found in RefSeq and NPDC. Further, 36% of NPDC GCFs are unique to individual strains (Extended Data Fig. 4A). In contrast, the most conserved GCFs in most Actinobacteria genera are found in over 90% of NPDC genomes (Extended Data Fig. 4B).

**Figure 4.**
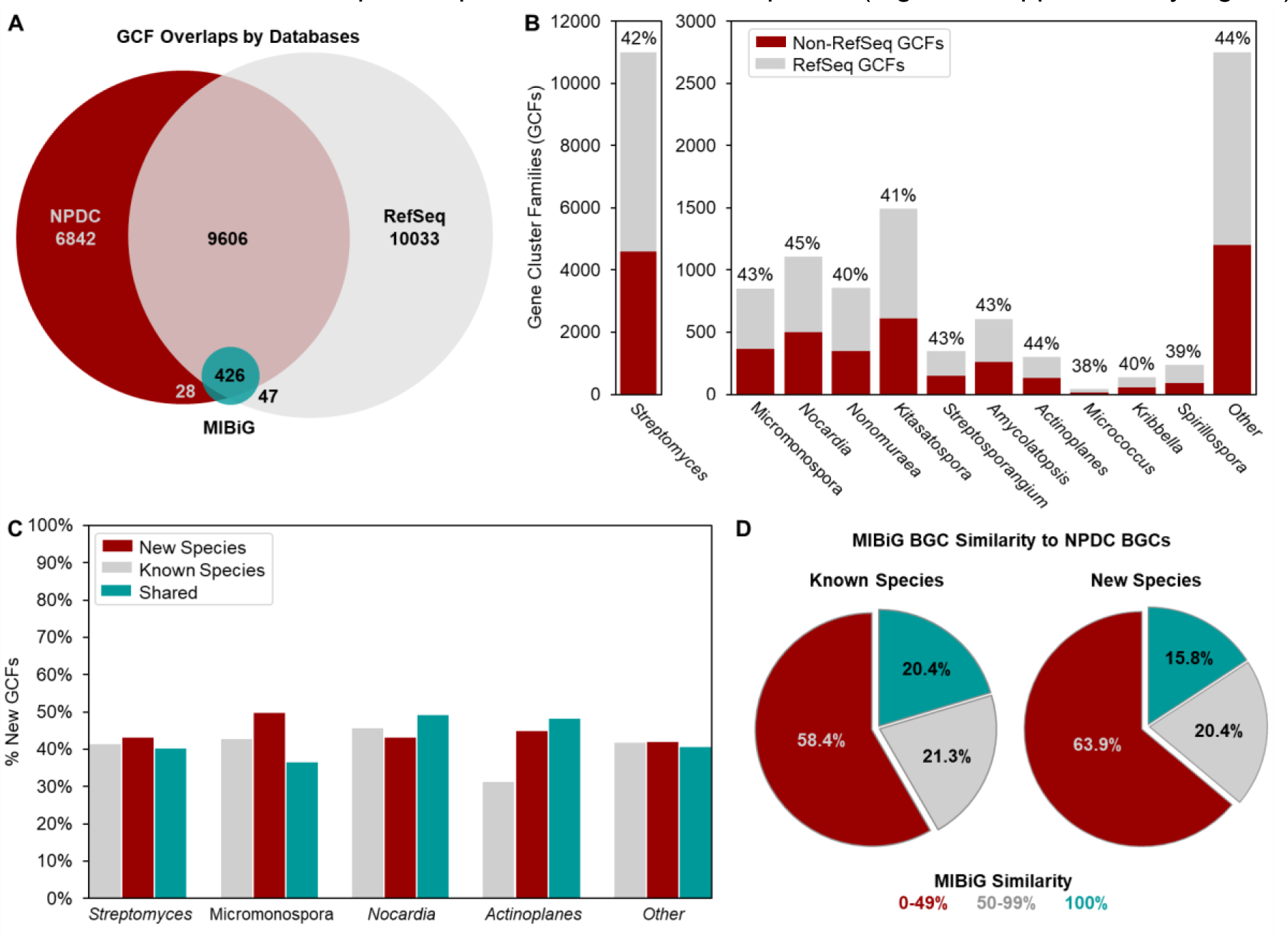
The NPDC genomes expand the accessible biosynthetic potential across different genera. (A) Distribution of GCFs in RefSeq (grey) and NPDC (red) genomes, with Actinobacteria MIBiG GCFs (teal) included separately. (B) Distribution of non-RefSeq GCFs (red) and GCFs observed in both NPDC and RefSeq genomes (grey) across representative genera as visualized in a stacked bar chart. Between 40-50% of total GCFs by NPDC genus are not observed in RefSeq (indicated by percentage). Also see Supplementary Fig. 17. (C) Percentages of non-RefSeq GCFs in known species (grey), proposed new species (red), and shared between them (teal) across representative genera. Also see Supplementary Fig. 18. (D) Distribution of NPDC BGCs by antiSMASH-assigned similarity to characterized MIBiG BGCs. A higher percentage of BGCs with low similarity to MIBiG BGCs is observed for new species relative to known ones.

Out of the 16,902 GCFs in NPDC genomes, most are expected to represent unexplored chemical space (Supplementary Fig. 16). Only 454 GCFs contain characterized BGCs from the MIBiG BGC database— including 28 GCFs with MIBiG entries lacking available genomes for the respective strains (Fig. 4A and Supplementary Table 5). ^36^ Nearly 12,000 total GCFs and 5000 non-RefSeq GCFs originate from NPDC *Streptomyces* strains, in line with the overall breakdown of NPDC GCFs (Fig. 4B). As with taxonomic diversity, not all genera show similarly high percentages of non-RefSeq GCFs. All genera will eventually reach a plateau at which they will experience diminishing rates of GCF novelty by genome sequencing, but the rate at which they will reach that plateau varies significantly. For example, the phylogenetically uniform and BGC-poor *Micrococcus* harbors almost no non-RefSeq GCFs despite dozens of NPDC genomes (Fig. 4B). Many prominent genera, such as *Streptomyces*, *Nocardia*, and *Actinoplanes*, well-regarded for their biosynthetic potential and with high taxonomic diversity (Fig. 1A-B), exhibit between 40-50% non-RefSeq GCFs (Fig. 4B), higher than most non-Actinobacteria genera. ^6^ In many of the rarer, underexplored Actinobacteria genera such as *Umezawaea* (58.1%), *Microbispora* (60.2%), and *Actinosynnema* (59.2%), this trend is even more pronounced, emphasizing them as valuable targets for future genome sequencing with their plateaus approaching significantly slower than in the well-studied genera (Supplementary Fig. 17).

While the level of GCF novelty varies by genus, the overall trend only slightly favors new species (42.0% non-RefSeq GCFs) over known species (41.1% non-RefSeq GCFs) among NPDC genomes (Fig. 4C). This small difference is unsurprising as more than 80% of NPDC GCFs are found in ten or fewer genomes (Extended Data Fig. 5), and most GCFs are strain- or species-specific, even when closely related taxonomically. ^6, 37^ This trend is also reflected by more than half of NPDC BGCs, from both GTDB known and new species, displaying little similarity to characterized BGCs in the MIBiG database (Fig. 4D). ^36^ Many genera or species, especially rare ones, have adapted to specific environments through the acquisition and evolution of tailored BGCs. In particular, Actinobacteria from the soil tend to have evolved more environment-specific GCFs, with two-thirds of GCFs identified in the genomes of NPDC Actinobacteria isolated from soil not being found in other environments (Supplementary Fig. 15B). This trend is more pronounced in certain rare genera, such as *Actinoplanes* (44.9% versus 31.3%), in which new species harbor far more non-RefSeq GCFs per strain than known species (Fig. 4C, Supplementary Fig. 18).

### The NPDC Portal enables virtual and physical access to the NPDC strain collection

The combined phylogenetic and biosynthetic diversity of the NPDC strains underlines its value to the research community, but its full potential can only be realized by making both the genomes and physical strains accessible. We have created an online portal (npdc.rc.ufl.edu/home) as a central hub for making this growing and incomparable resource accessible to enable users worldwide to take advantage of its unique resources. The NPDC Portal is centered around three primary functions: (i) strain and genome databases, (ii) integrated bioinformatics analyses, and (iii) strain requests for physical experimentation (Supplementary Fig. 19, Extended Data Fig. 6).

Many of the NPDC strains have metadata associated with them, so in addition to the genomes, the NPDC Portal contains any associated collection information (ecology, date, and location), media the strain has been grown in, genome-based taxonomy including related strains, and strain pictures (Supplementary Fig. 20). ^5^ Additionally, antiSMASH-generated BGCs are available for download or online browsing, and BGC annotations are provided for all genomes (Supplementary Fig. 21). NPDC strain, genome, and BGC metadata are fully searchable and filterable, and varying queries can be combined to enable combinatorial searches. For example, the query “NRPS|indole[BGC class] and Streptomyces [Taxonomy]” for the BGC database would provide a list of all *Streptomyces* BGCs with both NRPS and indole BGC classes annotated, while the query “>39[Num. of BGCs]” for the Genomes database would yield a list of all genomes containing 40 or more BGCs. At the sequence level, the built-in customized DIAMOND-BLASTP tool can be used to query protein databases encompassing all NPDC genomes or BGCs. ^38^ The NPDC BGC database, containing more than 224,000 BGCs, can be queried directly, with the added option to use up to five sequences simultaneously to explore co-occurrence within individual BGCs (Supplementary Fig. 22). The BLASTP results can be downloaded in various formats, including a standard BLAST tabular result, a multiFASTA file containing all protein hits, and a multiFASTA file containing all protein sequences from BGCs encoding hits. The multiFASTA files directly provide the sequences needed to construct sequence similarity networks (SSNs) or genome neighborhood networks (GNNs) using established tools such as EFI, which can be powerful for biosynthetic studies or prioritization of BGCs for discovery. ^39^ The BGC BLASTP results are also tied to NPDC GCF numbers and to antiSMASH-generated MIBiG BGC assignments, so related hits can be quickly extrapolated from a single BLASTP search.

### Applications of the NPDC Portal to explore the biosynthetic potential of the collection

To highlight the potential of the NPDC Portal, we have chosen four examples to emphasize possibilities for mining the BGC database with BLASTP, identifying associated multi-gene sets using BLASTP, and generating outputs to build SSNs and GNNs (Fig. 5-6, Extended Data Fig. 7). Our chosen examples cover resistance gene mining for the discovery of antibiotics, scaffold-specific mining for NPs to diversify the characterized chemical space, prioritization of enzymes for exploitation as biocatalysts, and mapping of biosynthetic proteins by GCFs.

**Figure 5.**
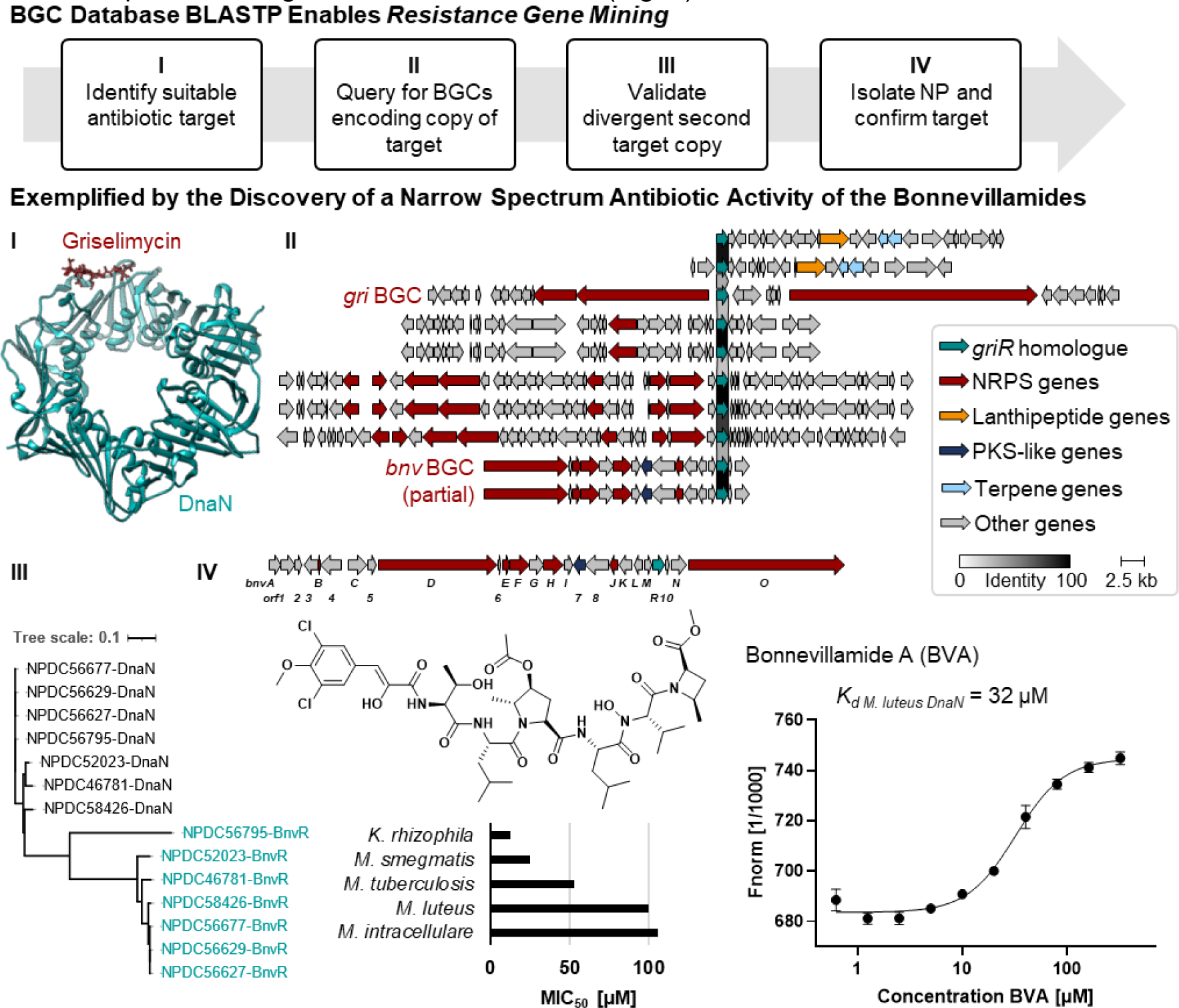
Resistance-guided genome mining for NPs acting on predicted potential targets. (I) Inspired by griselimycin, the only known DnaN-targeting NP to date and shown bound to DnaN (PDB: 6PTV), NPDC BGCs encoding DnaN homologues were identified, and (II) BGCs from four GCFs were aligned with the *gri* BGC using Clinker. ^40, 43^ (III) The DIAMOND-BLASTP tool allowed querying of the BGC database allowing exclusion of most primary metabolism hits. (IV) Bonnevillamide A, a NP associated with one GCF, ^41, 42^ was isolated and shown to bind the *Micrococcus luteus* DnaN and exhibited narrow-spectrum antibiotic activity.

**Figure 6.**
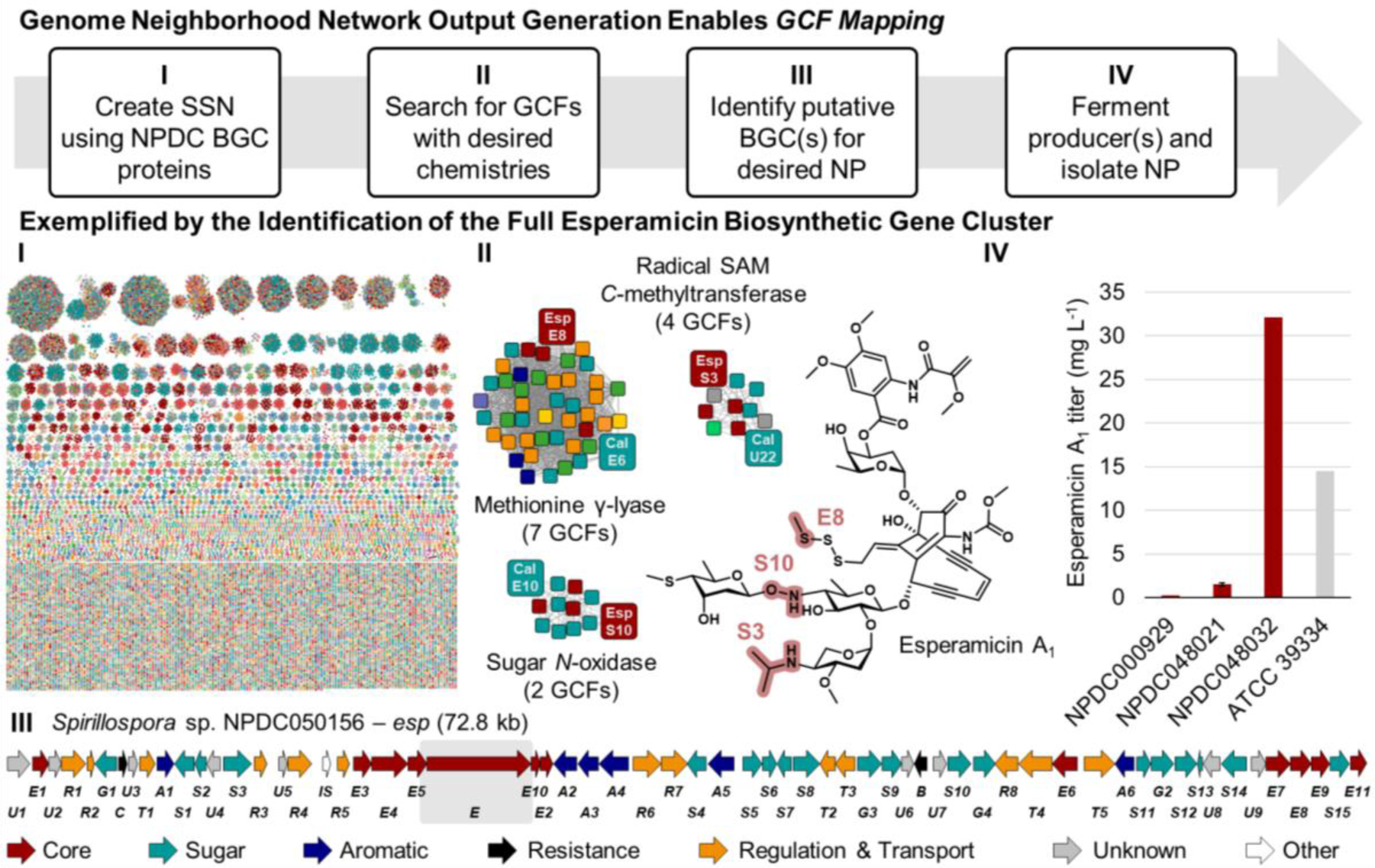
Identification of esperamicin BGC and alternative producers. (I) Using combined BLASTP queries for PksE and E4, all 31,029 protein sequences encoded by 659 NPDC enediyne BGCs were downloaded and grouped using the EFI SSN tool (e-value e^-100^). ^39^ Nodes were color-coded based on GCFs. (II) Zoomed-in protein clusters from the SSN showcase the diversity of GCFs encoding each protein responsible for the highlighted moieties in the *esp* BGC (red nodes) or *cal* BGC (teal nodes; see Supplementary Fig. 32). (III) The *esp* BGC is color-coded based on predicted gene product functions, and a grey box is around the previously sequenced region from the original ATCC producer. ^48^ (IV) Titers of ESP A_1_ were calculated for NPDC strains and compared to those from the original ATCC producer under the same conditions.

The NPDC BGC BLASTP database enables a user to search directly for protein hits within BGCs, which is especially advantageous to distinguish between primary metabolism proteins and those that have evolved into NP-specific roles. In the specific case of antibiotic resistance genes, this differentiation is particularly helpful as a second copy encoding a resistant variant is often localized within the BGC encoding the antibiotic NP to confer resistance to the producing organism. ^17^ Resistance gene-guided genome mining can therefore be used to prioritize BGCs for NPs with specific targets such as DnaN, the beta subunit of the DNA polymerase III. DnaN has been previously validated as an attractive novel antibacterial target; however, griselimycin, with its resistance protein GriR, remains the only DnaN-targeting NP to date (Fig. 5). ^40^ To identify other BGCs associated with NPs potentially targeting DnaN, we queried the NPDC BGC database for GriR homologues. After inspection for conservation across GCFs and for dissimilarity to primary metabolism DnaN copies, at least 14 BGCs across four GCFs from *Streptomyces* and *Kitasatospora*, each distinct from griselimycin, have been prioritized for ongoing follow-up studies (Fig. 5 and Supplementary Fig. 23).

One of the GCFs containing an unusual second copy of *dnaN* included the known *bnv* BGC for the bonnevillamides, a family of highly decorated linear non-ribosomal peptides with no observed antibiotic activity against *Staphylococcus aureus*, *Escherichia coli*, *Pseudomonas aeruginosa*, *Klebsiella pneumoniae*, or *Acinetobacter baumannii* (Fig. 5). ^41, 42^ Of the seven bonnevillamide BGCs found in NPDC genomes, all contained *dnaN* copies that were evolutionarily distinct from their conserved copy of *dnaN* that more closely matched those of other *Streptomyces* strains (Fig. 5). To validate if this pipeline could yield useful results, 3.3 mg bonnevillamide A (BVA) was isolated from *Streptomyces* sp. NPDC056627, as confirmed by 1D and 2D NMR, UV/Vis, and high-resolution mass spectrometry (HRMS) (Fig. 5, Supplementary Fig. 24-28, and Supplementary Table 6). As griselimycin is most potent against Actinobacteria including various *Mycobacterium* pathogens, BVA was tested against a diverse panel of pathogenic and non-pathogenic Actinobacteria. ^40^ BVA exhibited micromolar minimum inhibitory concentrations (MICs) against *Kocuria rhizophila* (12.5 μM), *Mycobacterium smegmatis* (25 μM), *Mycobacterium tuberculosis* (53 μM), *Micrococcus luteus* (100 μM), and *Mycobacterium intracellulare* (106 μM), though not against *Streptomyces albidoflavus* or *Mycobacterium abscessus*. To determine if BVA could bind DnaN, *dnaN* from *M. luteus* NPDC049463 was heterologously expressed in *E. coli* and its product purified for microscale thermophoresis (MST) experiments (Supplementary Fig. 29). MST showed a *K_d_* of 32.2 μM, validating that BVA binds to DnaN (Fig. 5).

In a related approach, the NPDC BGC database and its associated BLASTP tool facilitate the discovery of BGCs responsible for complex biosynthetic transformations or installation of multiple pharmacophores involving up to five proteins. Often, structural complexity is introduced by multi-enzyme machineries, and searching for individual instances of proteins is insufficient for identifying BGCs in which they co-occur to encode the biosynthesis of the target structural moiety. To demonstrate the efficacy of this approach, the five proteins responsible for introduction of an uncommon *N*-*N* bond in the antifungal fosfazinomycin via a glutamylhydrazine intermediate (FzmN-R) were simultaneously used to query NPDC BGCs (Extended Data Fig. 7A and Supplementary Fig. 30). ^44^ By downloading the hit BGCs and clustering them using the external tool BiG-SCAPE, four NP families could be grouped together, including fosfazinomycin-like, kinamycin-like, and lomaiviticin-like NPs (Extended Data Fig. 7A). ^45^ While some encoding strains represent alternative producers, many others represent novel variations, as well as a predicted novel *N*-*N* bond-containing NP scaffold without notable similarity to BGCs in the MIBiG database (Supplementary Fig. 30).

Beyond the discovery of NPs, the NPDC also facilitates the discovery of enzymes for use as biocatalysts for chemical synthesis. As above, searching the BGC database directly focuses the search on non-primary metabolism hits, and the accompanying downloadable BGC metadata file provides the BGC class and GCF for each hit. By using external tools such as the Clustal Omega sequence aligner in combination with NPDC metadata, the enzyme hits can be mapped to putative functions or substrates while exploring sequence diversity. ^46^ In an example, the NPDC was mined for BGC-associated α-ketoglutarate-dependent dioxygenases related to the asparaginyl oxygenase AsnO, resulting in 1223 hits from 163 GCFs spread across 318 species and 18 genera (Extended Data Fig. 7B). ^47^ In screening potential biocatalysts, a number of factors are critical, including solubility, expression levels, stability, kinetic parameters, lifetime, yield, solvent tolerance, and regio- and stereoselectivity. Given the importance of sequence diversity in screening for so many different parameters, NPDC users can use the “Download protein hits multiFASTA” option after BLASTP to obtain all hit sequences and metadata and select representative proteins to screen for their suitability for biocatalytic applications. The availability of GCF and BGC class information can also provide additional validation for researchers searching for enzymes expected to perform identical chemistry or to search for related chemistry much more rapidly than by looking at sequence alignments in the absence of additional information.

Analogous to identifying related enzymes, the NPDC BLASTP tool can deliver metadata and sequences for all proteins from a set of related BGCs. In combination with the web-based EFI SSN tool, large-scale comparison can be performed for thousands of individual proteins encoded by GCFs related by core genes. ^39^ In doing so, trends that are not readily apparent from smaller BGC alignments may be identified. One NP family where this analysis can provide valuable biosynthetic insights is the enediyne family of NPs. The enediynes are structurally related by their DNA-damaging 1,5-diyne-3-ene core moiety (Fig. 6), which is biosynthetically derived from the enediyne polyketide synthase (PKSE), its thioesterase, and three unique enediyne biosynthesis proteins (E3, E4, and E5). ^48, 49, 50^ However, unlike with the *N*-*N* bond-encoding BGCs, the 659 NPDC enediyne BGCs display substantial biosynthetic and taxonomic variety outside of these five gene products even for structurally-related compounds, leading to inconsistent subclassifications via whole-BGC clustering methods such as BiG-SCAPE. ^6, 45^ With the NPDC tools, all enediyne BGCs can be identified using the multi-protein BLASTP. Follow-up querying for specific proteins responsible for conserved moieties can then be used to classify different enediyne BGC subclasses, e.g. the methionine γ-lyase responsible for the installation of the methyltrisulfide in calicheamicin (Supplementary Fig. 31-32). Because the NPDC BLASTP tool maps each protein hit to a GCF, this process enables rapid identification of related but distinct GCFs containing key moieties.

For proof of principle, the NPDC tools were utilized to identify the previously unknown BGC associated with esperamicin A_1_ (ESP A_1_) (Fig. 6). ESP A_1_ was first discovered in 1985 from *Actinomadura verrucosospora* ATCC 39334, and it has been extensively studied as an antitumor agent. ^51, 52, 53^ However, the ESP A_1_ producer has never been sequenced aside from the single PKS gene, and no alternative producers of this valuable NP have ever been identified. ^48^ Of the nearly twenty characterized enediyne scaffolds, the antibody-drug conjugate payload calicheamicin from *Micromonospora echinospora* is most similar structurally (Supplementary Fig. 32). ^54^ To identify the *esp* BGC, if present in the NPDC genomes, the NPDC BGCs were queried with PKSE and E4, and the “Download BGC hits multiFASTA” option was selected to obtain the 31,029 protein sequences from all 659 BGCs. The protein sequences were clustered using the EFI SSN tool, and each node was mapped to one of the 93 enediyne GCFs (Fig. 6). ^39^ Key moieties shared between ESP A_1_ and calicheamicin include a methyltrisulfide (methionine γ-lyase), an alkylaminosugar (radical SAM *C*-methyltransferase), and a hydroxylaminosugar (sugar *N*-oxidase) (Fig. 6 and Supplementary Fig. 32), and fortunately, only one additional enediyne GCF could be associated with all three moieties. This GCF was found in four *Spirillospora* genomes: NPDC000929, NPDC048021, NPDC048032, and NPDC050156. In this BGC, genes could be predicted for the core, all four sugars, and the peripheral aromatic moiety (Fig. 6, Supplementary Fig. 32, and Supplementary Table 7); therefore, the strains were fermented to confirm the production of ESP A_1_. Two of the strains, NPDC000929 and NPDC048032, produced a compound with a mass consistent with ESP A_1_, so the fermentation of NPDC048032 was scaled up and 15.8 mg ESP A_1_ was purified and characterized by 1D and 2D NMR and HRMS (Supplementary Fig. 33-39 and Supplementary Table 8). In addition to identifying the *esp* BGC, this method identified four alternative producers of ESP A_1_, including NPDC048032, which had a titer (32.1 mg L^-1^) more than double that of the original producer (14.5 mg L^-1^) (Fig. 6).

## Discussion

The NPDC marks a transformative initiative by tapping into the vast biosynthetic potential of Actinobacteria for NP discovery. With over 122,000 strains, of which more than 84% are expected to be Actinobacteria, the NPDC collection represents a repository rich in genomic diversity. The analysis of the first 8,490 released genomes reveals the expansive taxonomic range within Actinobacteria, underscoring the significance of genome sequencing in preserving and exploring biodiversity.

One key aspect of genome sequencing is the reliable taxonomic classification of the strains in the collection, which significantly increases the value of the collection for microbiological and biosynthetic studies as there is a notable discrepancy between the phenotype- and genome-based taxonomic assignment. For many rare genera, this disparity can be explained by the evolving nature of taxonomic nomenclature or the close relatedness between genera. For example, among the 86 strains assigned as *Actinomadura*, nearly half can now be assigned as *Nonomuraea*, a genus that was not defined until the 1990s. ^55^ Others belong to *Spirillospora*, a genus whose members share very high 16S rRNA sequence similarity to some *Actinomadura* strains. ^56^ The updated taxonomies thus emphasize the critical importance of genome sequencing and the preservation of the genomic diversity encapsulated within the NPDC collection. Despite the differences between the phenotype- and genome-based taxonomies, this survey confirms that the NPDC collection has successfully prioritized environmental over health-associated Actinobacteria and therefore is complementary to existing genomes in RefSeq. Overall, the untapped diversity awaiting exploration within many rarer genera stored in the NPDC collection emphasizes the need for further sequencing and cataloguing efforts. By prioritizing these genera based on their observed intrageneric diversity, we can shed more light on this crucial branch of the tree of life while identifying promising candidates for future discovery campaigns. Within ∼7% of the NPDC collection, three new genera could be identified, serving as compelling examples of the microbiological discoveries possible through the NPDC and as inspiration for future studies enabled by the NPDC collection. However, while the microbiological facet of the NPDC holds substantial value, its greatest strength arguably resides in the biosynthetic machineries encoded by these strains.

Consistent with previous studies emphasizing the value of taxonomic distance for NP discovery, ^6^ the percentage of antiSMASH-annotated BGCs with 100% identity to MIBiG BGCs is lower in proposed new species compared to known species (Fig. 4D), but there is only a weak correlation with taxonomic novelty at the genus level (Fig. 4B-C). Instead, we observe that biosynthetic specialization occurs mostly at the individual strain level, as more than 80% of all GCFs are found in fewer than 10 strains, though continued sequencing of the strain collection is expected to improve this ratio (Extended Data Fig. 5). With more than 200,000 BGCs assigned in the initial NPDC genome release, prioritization of BGCs of interest and determination of the rate of return on sequencing for novel biosynthesis require grouping into GCFs. As structural diversity often is introduced by downstream modifications using single enzymes, grouping into GCFs is not a perfect measurement for NP type and scaffolds, but it does provide an approximation of the diversity of the BGCs and, to a lesser degree, NPs. Most of the GCFs that occur across species and genera often encode NPs such as siderophores or other common metabolites, e.g., geosmin, that are often critical for a strain’s survival independent of most environmental factors. ^57, 58^ Outside of these highly prevalent GCFs however, the 5% cut-off utilized to define distinct species provides sufficient genomic space for harboring new BGCs even within known species. For the median NPDC Actinobacteria genome, 5% is equivalent to more than 400 kb or twelve average-sized BGCs (33 kb in NPDC genomes). This observation is in line with the relatively consistent levels of biosynthetic novelty observed across Actinobacteria genera (Fig. 4B) by assuming a stable set of core GCFs and a more variable set of GCFs for rapid ecological adaptation. However, the argument for sequencing beyond known species and *Streptomyces* remains, as it is still rare for these non-core GCFs to cross between genera. ^6^

While the cost of genome sequencing has been dropping dramatically in recent years, it is still not considered cost-nor time-efficient for every lab to sequence all strains that have been collected—leading to sequencing primarily of individual pre-selected strains. Herein, we have shown the power of the approach of sequencing at scale, without being biased towards niche uses but also having the genotype-phenotype linkage lacking in large-scale metagenomic sequencing efforts. Our results underpin that broad sequencing efforts are especially valuable for studying less prevalent GCFs that will otherwise be missed. The presented online NPDC Portal relies on making robust and established methods easily applicable to this large dataset for expert and non-expert users alike, with the added benefit of increasing use of valuable strains that might otherwise stay hidden away, unsequenced, in a freezer in an individual lab. As maximizing the value of this resource goes beyond any single research group, our goal is that by (i) enabling queries previously inaccessible to non-computational researchers and (ii) making available strains that are prioritized by these queries, we can democratize genome mining for NP chemists and other non-specialists. Our presented resistance gene mining example (Fig. 5) shows that, while not replacing existing resistance-guided tools such as ARTS, ^59^ resistance gene-mining via the NPDC Portal offers a rapid survey of NPDC genomes without an advanced bioinformatics background and avoids the pitfall of a standard BLAST database that will return mostly primary metabolism hits.

The scope of this tool also goes far beyond resistance-guided genome mining, as highlighted by each of the remaining examples. The multi-BLAST function could, in addition to the presented example for mining *N-N* containing NPs (Extended Data Fig. 7A), be applied to many other strategies, such as looking for specific combinations of NRPS and PKS domains or biosynthetic enzymes responsible for installation of one or more pharmacophores or functional groups. For synthetic chemists, the access to a wide range of strains encoding a large selection of enzymes as potential biocatalysts (Extended Data Fig. 7B) allows circumvention of issues with stability, solubility, promiscuity, or other characteristics without resorting to enzyme engineering. Further, the ability to map biosynthetic proteins associated by a conserved chemical moiety or scaffold by GCF membership (Fig. 6) enables researchers to draw conclusions on the relative importance of specific proteins that might otherwise go unnoticed through examination of hundreds of BGC sequences or unbiased collections of protein sequences from large genome datasets. Application of this tool to any set of BGCs related by one or more genes therefore provides a more holistic view of BGC diversification, especially in larger NP families, to be harnessed for NP discovery and production. In the identification of the *esp* BGC and additional ESP A_1_ producers, the value of having multiple producers was highlighted, given the wide range of titers across different species.

With so many uncharacterized GCFs and low overlap with BGCs from the public databases, NPDC strains encode vast potential for NP and enzyme discovery. The NPDC Portal serves as a gateway to these resources, facilitating access to strains and associated metadata, while offering a suite of tools to expedite the functional or fundamental exploration of BGCs. By democratizing access to this wealth of strains and genomic data, the NPDC is positioned to accelerate advancements in drug discovery, biocatalysis, microbiology, and biotechnology.

## Materials and Methods

### General experimental procedures, bacterial strains, and chemicals

All 8490 strains were obtained from the Natural Products Discovery Center (NPDC) strain collection located at The Herbert Wertheim UF Scripps Institute for Biomedical Innovation & Technology in Jupiter, Florida. All strains were originally stored as either frozen glycerol spore stocks or lyophilized stocks. Most strains were cultured for 7 days in 5 mL modified tryptic soy broth (TSB) supplemented with optimized levels of trace elements (TSB+): ZnCl_2_ (40 mg L^-1^), FeCl_3_ • 6H_2_O (200 mg L^-1^), CuCl_2_ • 2H_2_O (10 mg L^-1^), MnCl_2_ • 4H_2_O (10 mg L^-1^), Na_2_B_4_O_7_ • 10H_2_O (10 mg L^-1^), and (NH_4_)_6_Mo_7_O_24_ • 4H_2_O (10 mg L^-1^). ^60^ When strains were grown on solid media, either ISP2 or ISP4 media were used. ^61^ Solid and liquid media were supplemented with nalidixic acid (30 mg L^-1^) and cycloheximide (50 mg L^-1^).

DNA concentrations were determined fluorometrically using the Quant-iT Broad Range and High Sensitivity dsDNA assays (Invitrogen) on an infinite M1000Pro plate reader (Tecan). DNA quality was determined using a NanoDrop 2000C spectrophotometer (Thermo Scientific). The Lotus library preparation kits were purchased from Integrated DNA Technologies (IDT). The Agencourt AMPure XP-PCR purification beads were purchased from Beckman-Coulter. All solvents (molecular biology grade) or chemicals were purchased from standard commercial sources.

^1^H, ^13^C, and 2D (COSY, HSQC, and HMBC) NMR spectra were recorded at 25 °C with an Avance 600 MHz spectrometer instrument (Bruker). Ultra-high performance liquid chromatography electrospray ionization high resolution mass spectrometry with UV detection (UHPLC-ESI-HRMS-UV) measurements were carried out on a Vanquish UHPLC system (Thermo Scientific) combined with an Orbitrap Exploris 120 mass spectrometer (Thermo Scientific) equipped with an electrospray ion (ESI) source, an Accucore^TM^ C18 column (100 mm × 2.1 mm, 2.6 μm) at 35 °C, and a diode array detector HL. Optical rotations were obtained using an AUTOPOL IV automatic polarimeter (Rudolph Research Analytical).

Plasmids and bacterial strains used in this study are listed in Supplementary Table 9. *E. coli* strains were grown on Luria-Bertani (LB) medium at 37 °C; kanamycin (Kan) was added as necessary at 50 μg mL^-1^. PCR primers were synthesized by Integrated DNA Technologies. High-fidelity DNA polymerase (Q5), restriction endonucleases, and T4 DNA ligase were purchased from NEB. Plasmid and gel extraction kits were acquired from Omega Bio-tek Inc. Electrophoresis was carried out using a Bio-Rad PowerPac 300. Plasmid sequencing was performed by Genewiz, Inc.

### Genomic DNA isolation

To isolate gDNA from bacterial cultures, samples were processed in batches of 96 cell pellets dissolved in 740 μL TE buffer (original dilutions based on culture density). Cells were lysed using 20 μL lysozyme (100 mg mL^-1^) and 10 μL achromopeptidase (5 mg mL^-1^) in the presence of 5 μL RNase A (20 mg mL^-1^) at 37 °C for 2 h with periodic inversions. Next, 40 μL sodium dodecyl sulfate (10% w/v) and 10 μL proteinase K (8 mg mL^-1^) were added, mixed, and incubated at 56 °C for 16-24 h until samples were clear. 100 μL NaCl (5 M) was then added to the samples and mixed, followed by the addition of 500 μL 25:24:1 phenol:chloroform:isoamyl alcohol. Centrifugal phase separation (10 min at 13.2k rpm) was aided by the addition of 75 μL Molykote high vacuum grease (DuPont). A second extraction was performed using 500 μL 24:1 chloroform:isoamyl alcohol and separated by centrifugation (10 min at 13.2k rpm). A final ethanol precipitation with sodium acetate (final concentration of 0.3 M) was performed, and the final gDNAs were resuspended with 100 μL low-EDTA TE buffer (10 mM Tris-HCl, pH 8.0, 0.1 mM EDTA). gDNA qualities were determined spectrophotometrically, and gDNA concentrations were determined fluorometrically.

### Genome sequencing, assembly, and annotation

Most library preparations for short-read sequencing were carried out using IDT Lotus reagents with a modified automated protocol on an Agilent Bravo liquid handling robot. The manufacturer’s instructions were utilized at a quarter scale except for the final bead purification and elution step, which was conducted at full volume. The libraries were PCR-amplified. Each library was indexed with IDT UDI 10 bp barcodes and pooled in batches of 384 normalized samples each. Genomes were sequenced using 2×150 sequencing on an Illumina NovaSeq 6000 with an S4 flow cell (384 samples per lane).

To process sequencing reads, BBDuk was used to remove contaminants, trim reads that contained adapter sequence and terminal homopolymers of five or more ‘G’ bases, and remove reads containing one or more ’N’ bases or having length ≤ 51 bp or 33% of the full read length. ^62^ Reads mapped with BBMap to masked human references at 93% identity were separated into a chaff file. ^62^ Further, reads aligned to masked common microbial contaminants were separated into a chaff file.

The following steps were performed for genome assembly: (1) artifact-filtered and normalized Illumina reads were assembled with SPAdes (version v3.14.1; –phred-offset 33 –cov-cutoff auto -t 16 -m 64 –careful -k 25,55,95); (2) contigs were discarded if the length was < 1 kb (BBTools reformat.sh: minlength=1000 ow=t). ^62, 63^ CheckM then was used to calculate the contamination and completeness level of genomes. ^64^

Genomes having ≥ 95% completeness and ≤ 10% contamination were kept, while others are discarded. GTDB-Toolkit was used to annotate the taxonomy of genomes. ^24^ Prokka was used to predict and annotate coding sequences in the genomes. ^65^

### Construction of NPDC Portal

The NPDC Portal is written in Python 3.11 using Flask (v2.3) and Jinja2 (v3.1) libraries. The web server is linked to three separate SQLite databases for storing genomic, users, and BLAST query data. The BLAST function is provided via a separate Python script running in the background, calling DIAMOND-BLASTP (v2.0.15) on a pre-built NPDC protein database (default sensitivity settings with modified query parameters: e-value = 1e^-10^, query-cover = 80%, identity-threshold = 40%).^38^ Raw text results from DIAMOND were then parsed and stored accordingly in the query database to facilitate interactive viewing and downloading. The source code for the NPDC Portal is available at https://github.com/shenlabnp/npdc_portal.

For querying the Strain, Genome, and BGC metadata tables on the NPDC Portal, the column being searched should be specified in brackets (“query[category name]”), and multiple categories may be queried concurrently using “and” to connect the queries. For queries involving multiple BGC classes or dates, the terms are separated by “|”. Numerical queries can be expanded to ranges using comparison operators “>”, “<”, or “-“. For querying MIBiG hits, the MIBiG BGC number must be used, e.g., “BGC0000271[MIBiG hit]”.

### Bioinformatics analysis

Assembled genome data were downloaded in GenBank format (.gbk) from the NCBI RefSeq FTP server using the ncbi-genome-download tool (April 2022). ^66^ The list of RefSeq accessions was cross-referenced with the GTDB taxonomy RS-207 (April 2022) to get the whole-genome-based taxonomic classification of each genome. ^16, 24^ NCBI RefSeq genomes without any GTDB annotation (i.e., more recent and low-quality genomes) were filtered out, leaving a final list of 215,240 genomes. These genomes were selected due to their high quality, enabling a useful comparative BGC analysis, and due to being included in the current version of GTDB, enabling a useful taxonomic comparison.

Assembled NPDC genomes were processed using the GTDB-TK pipeline (v2.1.0) to assign taxonomy based on the same GTDB RS-207 database. ^24^ Additionally, an internal taxonomic grouping was performed by calculating pairwise ANI-like distances between NPDC genomes using Mash (v2.3). ^26^ Then, a complete-linkage hierarchical clustering was run with a threshold value of 0.05 (roughly equivalent to 95% ANI value, a general cutoff for species demarcation. ^67^ Finally, the genomes were categorized into "known species" for genomes successfully annotated by GTDB down to the species level and "new species" for genomes without corresponding GTDB species annotation. The Mash-derived genome clusters were used to assign putative species grouping to all genomes categorized as the latter.

AntiSMASH (v5.1.1) was run for each genome from NPDC and NCBI RefSeq datasets to identify BGCs. ^68^ The two datasets were complemented by another set of 1910 experimentally-validated BGCs from the MIBiG v2.0 database. ^36^ All GenBank BGC files were run using a modified version of BiG-SLiCE which uses cosine-based distance to enable accurate BGC clustering. ^69^ A cosine distance threshold of 0.4 was used to group BGCs into GCFs. The absence and presence of GCFs across the three datasets could then be tracked, with a GCF determined to be present in a dataset if at least one member BGC is from the given dataset.

To create the phylogenetic trees, the GTDB-Tk-v2 toolkit was utilized using the identify and align commands with combinations of NPDC and RefSeq genomes. ^24^ The GTDB-Tk-v2 outputs were converted into phylogenetic trees using fasttree and visualized on iTOL. ^70, 71^ For further taxonomy-based analyses, genera grouped in GTDB under the same parent name but considered different genera (e.g., *Arthrobacter*, *Arthrobacter_C*) were compared together to be consistent with the taxonomic assignments provided in RefSeq (Supplementary Table 10).

For the rarefaction curve in Extended Data Fig. 5C, starting with pairs of NPDC IDs and GCF IDs, the fraction of GCFs found in only a single genome were calculated at different numbers of randomly selected NPDC Actinobacteria genomes (N = 500, 1000, 2000, 3000, 4000, 5000, 6000, and 7139). This calculation was performed 100 times for each value of N. From there, a rarefaction analysis was performed, with curve fitting used to extrapolate beyond the current dataset out to 120,000 genomes. A 95% confidence interval was also calculated for the extrapolated portion of the dataset.

### Genome mining

For DnaN-encoding BGC mining, the amino acid sequence of GriR encoded by *Streptomyces muensis* was used as the query sequence for the NPDC BGC Database (sequence retrieved from the griselimycin BGC; MIBiG BGC0001414). ^36, 40^ For each GCF associated with a NPDC BLASTP hit, the DnaN homologue was manually compared with any other homologues encoded by the same genome (if any) and DnaN copies not associated with BGCs from related strains. All antiSMASH-annotated BGCs encoding divergent DnaN homologues were downloaded as .gbk files from the NPDC Portal and aligned with the griselimycin BGC (MIBiG BGC0001414) using Clinker. ^43^

For mining for *N*-*N* bond-encoding BGCs, the five proteins responsible for installation of the *N*-*N* bond in fosfazinomycin biosynthesis (FzmN/FzmO/FzmP/FzmQ/FzmR) were used as simultaneous queries of the NPDC BGC Database (sequences retrieved from the *Streptomyces* sp. WM6372 fosfazinomycin BGC; MIBiG BGC0000937). ^36, 44^ All BLASTP BGC hits were downloaded as .gbk files from the NPDC Portal and clustered using BiG-SCAPE with standard parameters and visualized using Cytoscape. ^45, 72^ BGC sequences from each BiG-SCAPE cluster were aligned together with the fosfazinomycin BGC and either the *Streptomyces murayamaensis* kinamycin BGC (MIBiG BGC0000236) or the *Salinispora pacifica* lomaiviticin BGC (MIBiG BGC0000240) using Clinker. ^43, 44, 73^ For the example of searching for BGCs that also encoded a phosphonate NPs, the protein FzmC from the fosfazinomycin BGC was used in conjunction with FzmN/FzmO/FzmP/FzmR to query the NPDC BGC Database. For the biocatalysts, the AsnO protein sequence was used as a query against the NPDC BGC database, and selected representative homologues were aligned using Clustal Omega. ^46^

For mining enediyne BGCs, two conserved core enediyne proteins, TnmE and TnmE4, from the tiancimycin A BGC (MIBiG BGC0001378) were used as simultaneous queries for the NPDC BGC Database. ^48, 49^ The resulting BGC hits multiFASTA file was downloaded, containing the sequences of all proteins encoded by any BGC containing homologues of both TnmE and TnmE4. The multiFASTA file was uploaded to the EFI SSN tool using an e-value of e^-100^ and visualized in Cytoscape. ^39, 72^ The resulting SSN was color-coded in Cytoscape using a presence-absence map of key conserved proteins associated with either calicheamicin-type enediynes, anthraquinone-fused enediynes, or 9-membered enediynes.

### Electron microscopy

Colonies were fixed in 4% paraformaldehyde and 2.5% glutaraldehyde in 1X PBS, containing 0.1% Triton X-100 pH: 7.2. Fixed cells were processed in 78 μm microporous specimen capsules (EMS) and with the aid of a Pelco BioWave laboratory microwave (Ted Pella). Fixed colonies were washed with 1X PBS, pH 7.22, post fixed with buffered 2% OsO_4_, water washed and dehydrated in a graded ethanol series 25% through 100%, with 10% increments. Samples were critical point dried using the Tousimis Autosamdri-815 (Tousimis). Once dried, the samples were mounted onto a 12 mm Carbon Conductive Adhesive Tab and aluminum stub (EMS) and then sputter coated with gold/palladium under argon gas, using the Denton Desk V sputter coater (Denton Vacuum). Samples were imaged with Hitachi SU-5000 FE-SEM (Hitachi High Technologies in America) with use of EM Wizard software.

### Production and isolation of esperamicin A_1_

Fresh spores of *Spirillospora* sp. NPDC048032 from ISP2 plates were used to inoculate seed cultures (50 mL in 250 mL baffled flasks, 0.5% glucose, 1% pharmamedia, 2% corn starch, 1% primary yeast, 0.2% calcium carbonate, pH 7.2) grown for 4 days at 28 °C, 250 rpm. ^74^ Production cultures were fermented under the same conditions in production medium (6% cane molasses, 2% corn starch, 2% fish meal, 0.01% CuSO_4_ x 5 H_2_O, 0.2% CaCO_3_, 0.00005% NaI, pH 7.2) inoculated with 8% (v/v) seed culture. ^74^ Small scale cultivation was performed using 50 mL in 250 mL, and large-scale cultivations using 12 x 400 mL in 2 L baffled flasks. All cultures were grown for 8 days and supplemented with 2% (w/v) Diaion HP-20 resin on day 4. Resin and cells were collected by centrifugation and extracted with EtOAc. The small-scale extracts were concentrated to dryness, redissolved in MeOH (1 mL), cooled to -80 °C for 1 h, and centrifuged at 18.100 x g for 15 minutes. 10 µL of the obtained extract was analyzed on an Agilent 1260 Infinity LC coupled to a 6230 TOF equipped with an Agilent Poroshell 120 EC-C18 column (50 mm × 4.6 mm, 2.7 µm) at 40 °C using a linear gradient from 5% to 100% MeCN (0.1% formic acid)/H_2_O (0.1% formic acid) over 8 min, a flow rate of 0.4 mL min^-1^ and UV detection at 254 nm. 1.23 g crude material containing 208 mg of ESP A_1_ was obtained from the large-scale fermentations. The sample was loaded onto a 3 x 40 cm LH-20 Sephadex column and eluted with 10% EtOAc in MeOH. Fractions were monitored by TLC and ESP A_1_ containing fractions subjected to preparative HPLC after solvent reduction under reduced pressure. Separation was achieved on an Agilent 1260 Infinity system equipped with an Eclipse XDB-C18 column (7 µm, 21.2 x 250 mm) employing the following conditions: 5% to 40% MeCN + 0.1% formic acid in H_2_O + 0.1% formic acid over 5 min, 40% to 65% over 15 min, 65% to 95% over 1 min, flow rate 17.0 mL min^-1^, UV detection at 254 nm. Fractions near *t_R_* = 12.3 min were frozen immediately and lyophilized to yield 15.8 mg ESP A_1_ as an off-white solid (11.92 µmol, 7.6% isolated yield).

To compare ESP A_1_ titers, *Spirillospora* sp. NPDC000929, *Spirillospora* sp. NPDC048021, *Spirillospora* sp. NPDC048032, and *Actinomadura verrucosospora* ATCC 39334 were cultivated in 250 mL flasks as described above. Each strain was grown in triplicate, and after fermentation for eight days, the cultures were extracted. The extracts were analyzed as described above. The UV signal for ESP A_1_ at 254 nm was integrated, and the amount of ESP A_1_ in each extract was calculated based on a calibration curve using the NMR-confirmed isolated ESP A_1_ from NPDC048032.

### Production and isolation of bonnevillamide A

Fresh spores of *Streptomyces* sp. NPDC056627 from GYM plates were used to inoculate seed cultures (50 mL in 250 mL baffled flasks, TSB medium) grown for 3 days at 28 °C, 220 rpm. Production cultures (40 x 50 mL) were fermented under the same conditions in production medium (40 g L^-1^ dextrin, 7.5 g L^-1^ tomato paste, 2.5 g L^-1^ N-Z-amine, 5 g L^-1^ primary yeast, pH 7.0) inoculated with 2% (v/v) seed culture. ^75^ All cultures were grown for 9 d and supplemented with 2% (v/v) of a Diaion HP-20/Amberlite XAD-16 resin mixture (1:1) on day 3. Resin and cells were collected by centrifugation and consecutively extracted with MeOH and acetone. After evaporation of the organic solvents, the extract was partitioned between water and chloroform. The obtained chloroform layer was concentrated under reduced pressure, loaded onto a 4 x 50 cm LH-20 Sephadex column and eluted with MeOH. Fractions were monitored by HPLC-MS and bonnevillamide A containing fractions subjected to semipreparative HPLC after solvent reduction under reduced pressure. Separation was achieved on an Agilent 1260 Infinity II system equipped with a Zorbax SB-C18 column (5 µm, 9.4 x 250 mm) employing the following conditions: 25% to 65% MeCN + 0.1% formic acid in H_2_O + 0.1% formic acid over 30 min, flow rate 5.0 mL min^-1^, UV detection at 294 nm. Fractions near *t_R_* = 19.9 min were lyophilized, yielding 3.3 mg bonnevillamide A as a red-brown amorphous solid.

### Production of DnaN

The *dnaN* gene was PCR-amplified from genomic DNA prepared from *M. luteus* NPDC049463 using primers Mluteus_DnaN.FOR (AAAACCTCTATTTCCAGTCGATGCAGCTCGTGAAGTTCA CCGTC) and Mluteus_DnaN.REV (TACTTACTTAAATGGAGGTGGGACAGGTACACGG). The PCR product was purified, treated with T4 polymerase, and cloned into pBS3080 according to ligation-independent procedures to afford pBS24001. ^76^ The cloned product was transformed first into *E. coli* NEB Turbo cells, and after sequencing verification, into *E. coli* BL21(DE3) to yield strain SB24001. SB24001 was cultured in 3 L LB broth inoculated with 1% overnight culture at 37 °C and 220 rpm until an OD_600_ was reached. The culture was cooled to 4 °C, gene expression was induced with the addition of 0.125 mM isopropyl β-d-1-thiogalactopyranoside (IPTG), and the cells were grown around 18 h at 18 °C and 220 rpm. After harvesting the cells by centrifugation at 4000 g for 20 min at 4 °C, the pellet was resuspended in lysis buffer (50 mM Tri, pH 8.0, containing 300 mM NaCl and 10 mM imidazole), lysed by sonication, and centrifuged at 15,000 g for 30 min at 4 °C. The supernatant was purified in two steps using an ÄKTA FPLC system (GE Healthcare Biosciences): a) nickel-affinity chromatography equipped with a HisTrap HP, 5 mL column (GE Healthcare Life Sciences), which was first washed with 100 mL Wash buffer (50 mM Tris, pH 8.0, containing 300 mM NaCl and 20 mM imidazole) and then eluted with a gradient increasing the concentration (0–50%, 120 mL) of Elution buffer (50 mM Tris, pH 8.0, containing 100 mM NaCl and 500 mM imidazole); b) size-exclusion chromatography equipped with a Superdex 200 16/600 column (GE Healthcare Life Sciences) pre-equilibrated with SEC buffer (50 mM sodium phosphate, pH 7.4, 150 mM NaCl), then eluted with one column volume of SEC buffer at 1 mL min^-1^. The resultant protein with an *N*-terminal His_6_-tag was concentrated using a 10 kDa Amicon Ultra-15 concentrator (Millipore). Protein concentrations were determined from the absorbance at 280 nm using a molar absorptivity constant (ε_280_ = 22,920 M^-1^ cm^-1^) and molar mass (42.278 kDa). Individual aliquots of DnaN were stored at -80 °C until use.

### Microscale thermophoresis of DnaN

Microscale thermophoresis binding experiments were performed according to the standard protocol provided by NanoTemper on a Monolith NT.115Pico. *M. luteus* DnaN was labeled using the Monolith Series Protein Labeling Kit Red-NHS and diluted to 10 nM with MST buffer (20 mM Tris, pH 7.4, 150 mM NaCl, 0.05% Tween-20). A serially diluted bonnevillamide A DMSO stock solution was added to obtain the indicated concentrations and incubated with the protein for 10 min before loading the capillaries. Data analysis was carried out using MO.Affinity Analysis v2.3 (NanoTemper) and visualized in GraphPad Prism.

### MIC determinations

*Kocuria rhizophila* ATCC 9341*, Micrococcus luteus* NPDC049463, *Streptomyces albidoflavus* J1074, and *Mycobacterium smegmatis* ATCC 607 were diluted to approx. 5 × 10^5^ cells mL^-1^ in TSB medium (120 μL, 96 well-plates) and subjected to a serial dilution of bonnevillamide A at a concentration range from 50 to 0.4 μM in triplicate. After 16 h incubation at 28 °C for *K. rhizophila*, *M. luteus*, *S. albidoflavus* and 37 °C for *M. smegmatis,* OD_600_ was assessed on a SpectraMax M5 plate reader for calculation of the minimal inhibitory concentration.

*Mycobacterium tuberculosis* R37a ATCC 25177, *M. intracellulare* DSM 43223, and *Mycobacterium abscessus* ATCC 19977 were grown on separate plates in 8 × 8 well grids, with 200 µL of Middlebrook 7H10 agar with 10 % OADC supplement, 0.2 % glycerol per well. Each row of wells was incubated with 10 µL of BVA, serially diluted in DMSO from 625 µM down to 5 µM. 5 µL of bacterial culture (100 colony forming units - CFUs) was added to each well. Two plates were independently prepared for each experiment. Plates were incubated in a CO_2_ incubator at 37 °C until visible growth was observed in control wells (without BVA). Minimal inhibitory concentration was identified as the lowest concentration of BVA that prevented bacterial growth.

## Data Availability

All data are available in the main text, the supplementary materials, or by request of the corresponding author. DNA sequences from NPDC genomes are available through the NPDC Portal (https://npdc.rc.ufl.edu/home).

## Acknowledgments

The project was supported in part by the Natural Products Discovery Center at the Herbert Wertheim UF Scripps Institute for Innovation and Technology and NIH grant GM134954 (B.S.). E.K., C.N.T., and A.D.S. were supported in part by NIH postdoctoral fellowships GM134688, GM128345, and GM133114, respectively. C.D.B., A.F.K., and L.L. were supported in part by the German Research Foundation (DFG) Walter-Benjamin postdoctoral fellowships 501989175, 514898299, and 509137346, respectively. C.-J.Z. was supported in part by a scholarship from the Chinese Scholarship Council (201803170010). B.-Y.F. was supported in part by the Jiangsu Overseas Visiting Scholar Program for University Prominent Young & Middle-aged Teachers and Presidents. Whole genome sequencing was carried out in part by the U.S. Department of Energy Joint Genome Institute (https://ror.org/04xm1d337), a DOE Office of Science User Facility that is supported by the Office of Science of the U.S. Department of Energy operated under Contract No. DE-AC02-05CH11231.JGI), under the Community Science Program (proposal: 10.46936/10.25585/60001355), by Corteva Agriscience^TM^ under a collaborative project, and by the UF ICBR NextGen DNA Sequencing Core Facility (RRID:SCR_019152). We would like to acknowledge Karen Kelly at UF ICBR Electron Microscopy for help with imaging of the *Spongisporangium*, *Streptodolium*, and *Uniformispora* strains (RRID:SCR_019146).

## Author Contributions

Conceptualization (EK, SAK, DY, CNT, CDB, MJS, BS); Methodology (EK, SAK, DY, CNT); Software (SAK, DY, CMD); Formal analysis and visualization (EK, SAK, CDB, LL); Investigation (EK, SAK, DY, CNT, CMD, LLF, CDB, JRS, ADS, CG, SM, GL, KV, FY, PS, LL, AN, AFK, AJC, WY, BF, YJ, AA, CJZ, LS, TMC, MJS, MGC, LPSC); Writing – Original Draft (EK, CDB, BS); Writing – Review & Editing (EK, CDB, BS); Supervision, project administration, and funding acquisition (BS).

## Extended Data Figures

**Extended Data Fig. 1.**
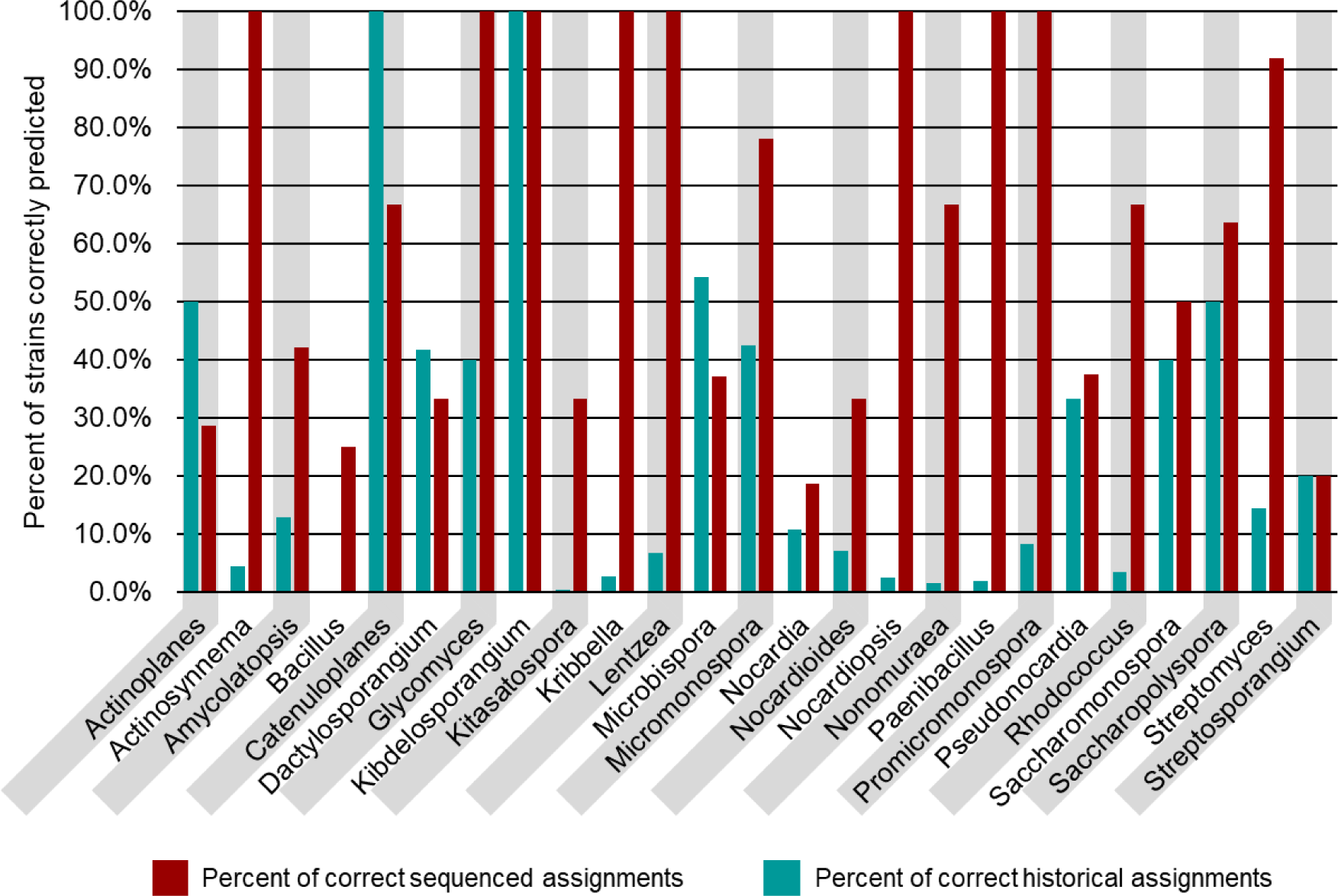
Comparison of historical and sequence-based taxonomies of NPDC strains. Approximately 10% of NPDC strains had morphology-based taxonomies assigned. See Supplementary Table 3 for additional values. The teal bars represent the percentage of isolates in a whole-genome sequencing-determined genus that were properly assigned based on historical classification. The red bars represent the percentage of historically classified isolates whose genus was confirmed based on whole-genome sequencing.

**Extended Data Fig. 2.**
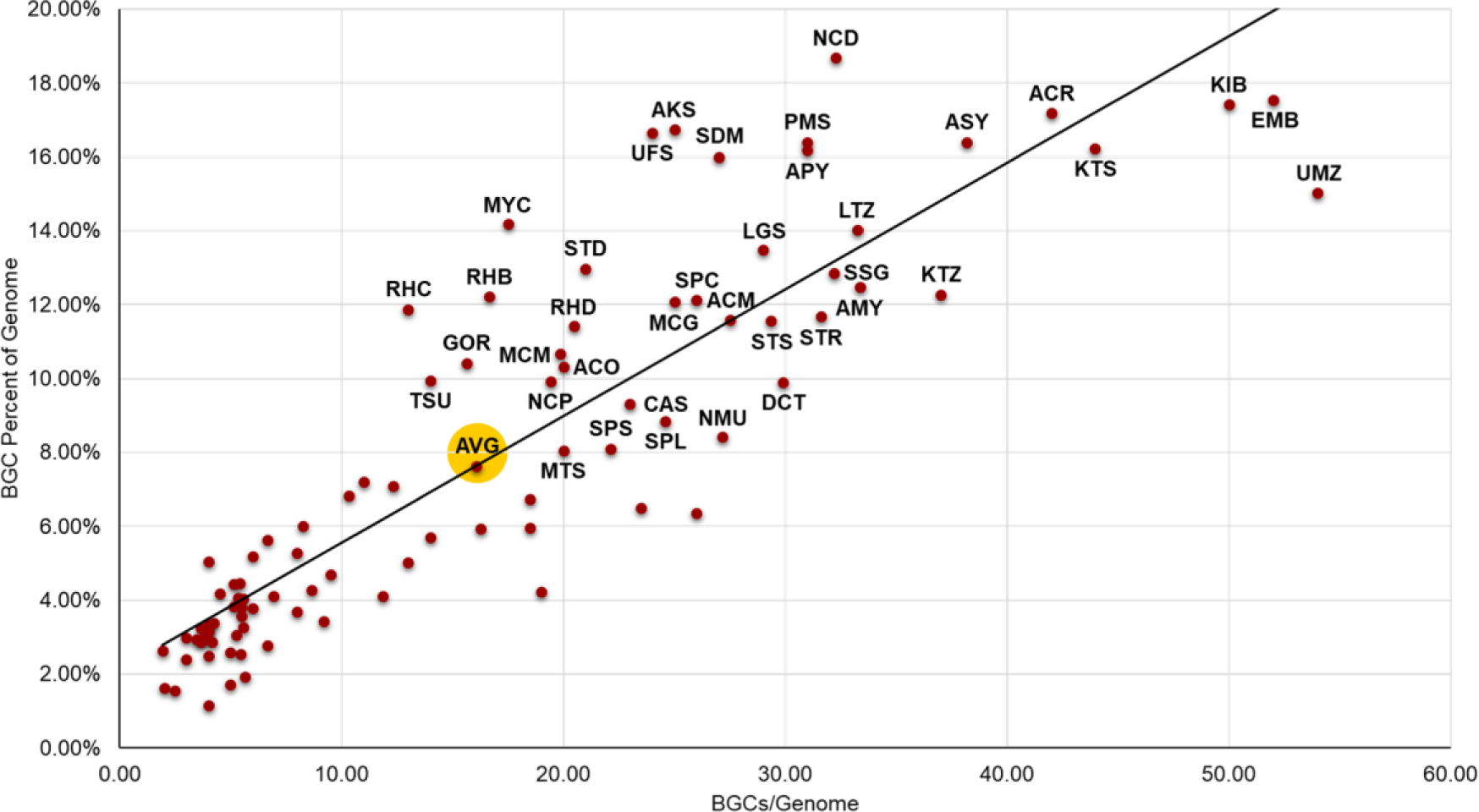
BGC dedication by Actinobacteria genus. The number of BGCs per genome follows a roughly linear relationship with the percentage of genome encoding BGCs, with some notable outliers showing up as the number of BGCs increases. All genera with above average percentages of the genome dedicated to BGCs are labeled. The average is indicated by AVG (highlighted in yellow). See Supplementary Table 10 for genus abbreviations.

**Extended Data Fig. 3.**
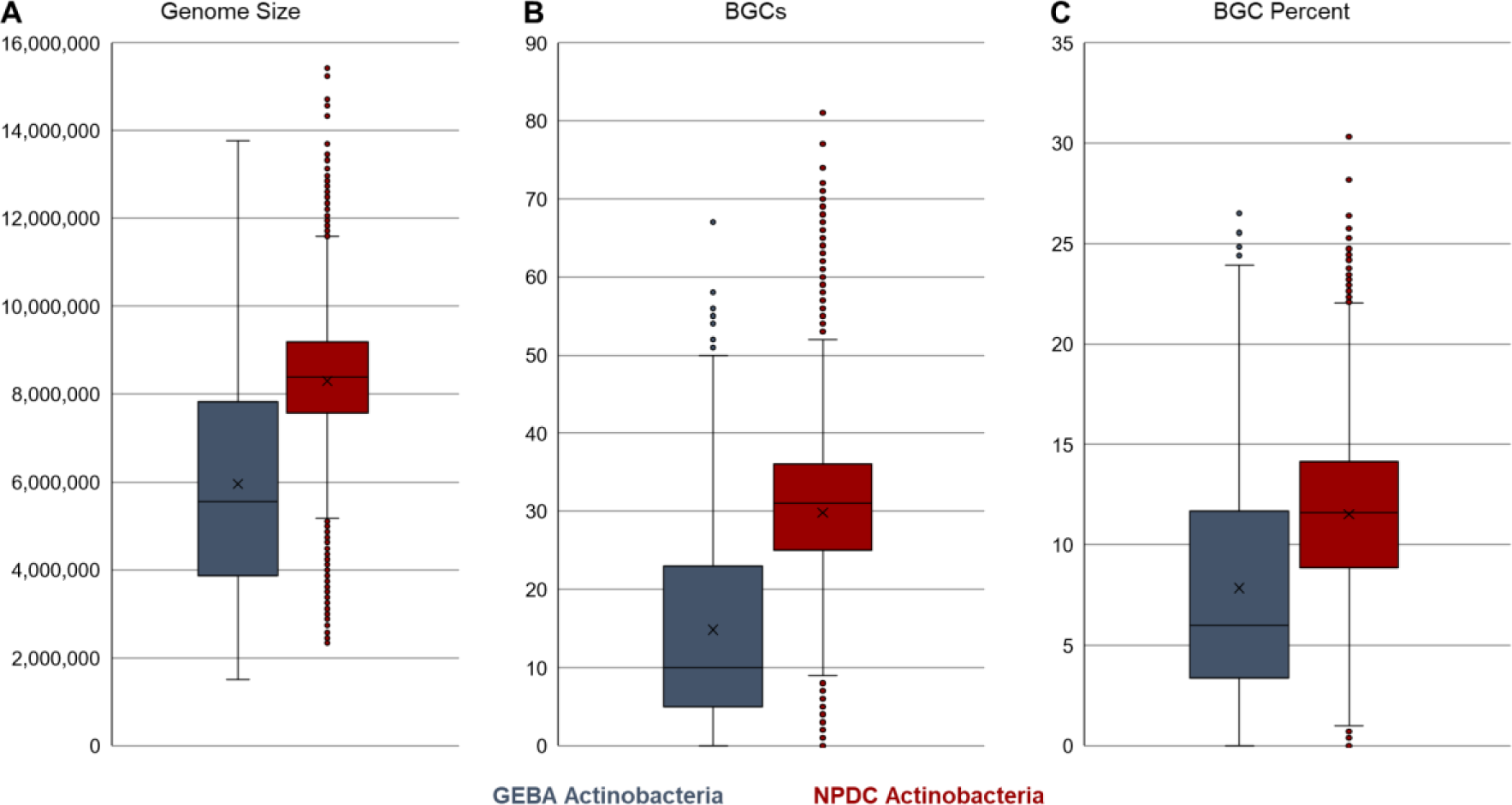
Comparison of GEBA and NPDC Actinobacteria genomes. (A) Comparison of genome sizes shows that the NPDC strains, collected for their NP biosynthetic potential, typically have much larger genomes than those from the Genomic Encyclopedia of Bacteria and Archaea (GEBA) Actinobacteria, which were primarily sequenced based on taxonomic diversity rather than biosynthetic potential. ^15^ Similar trends were observed for the number of antiSMASH-predicted BGCs per genome (B) and the percentage of the genome dedicated to those BGCs (C).

**Extended Data Fig. 4.**
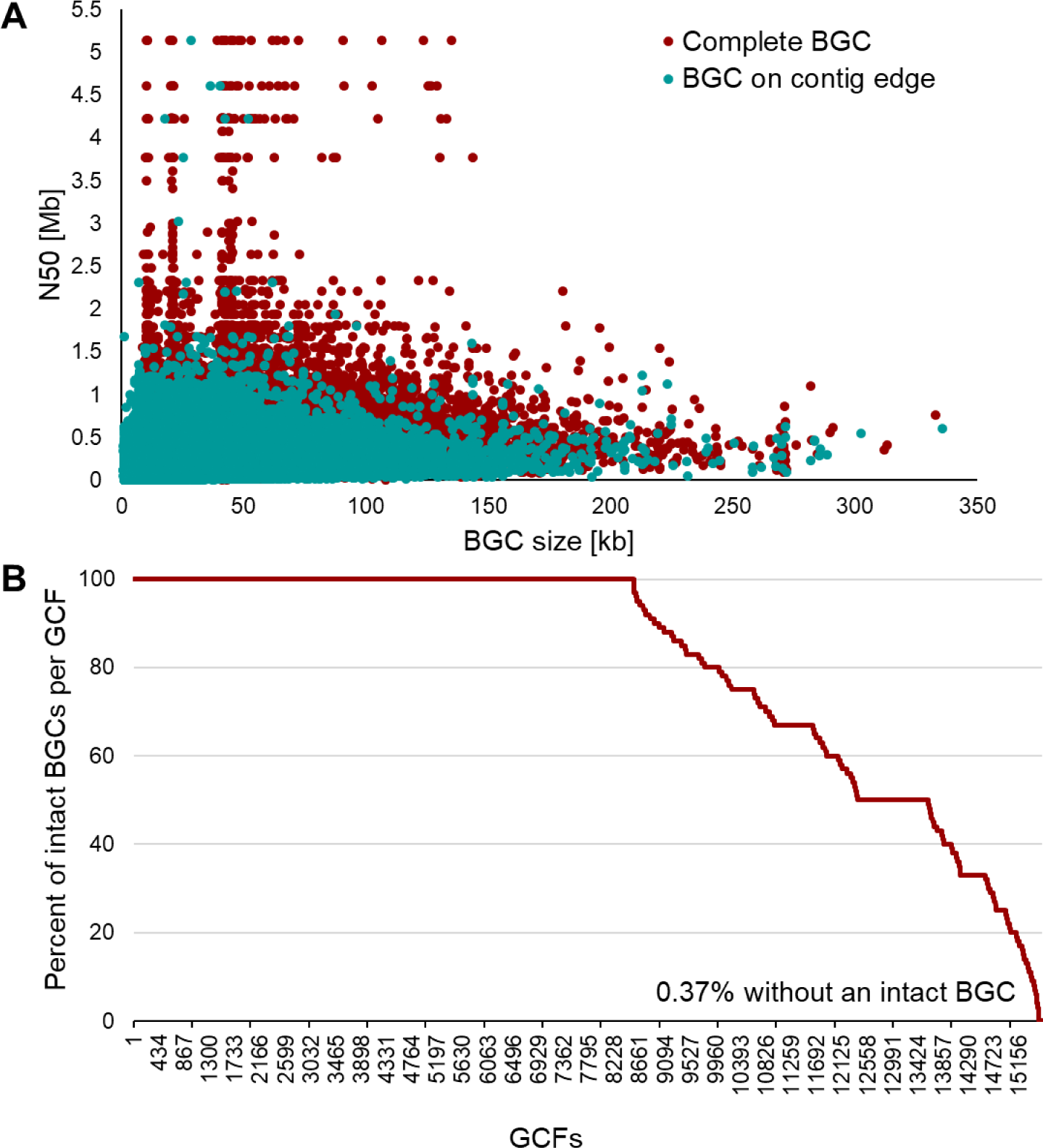
Completeness of NPDC BGCs and GCFs. (A) Relationship between intact BGCs and genome quality. In general, genomes with larger N50 values display a higher rate of BGCs predicted to be intact by antiSMASH. ^68^ However, even genomes with small N50 values contain large intact BGCs. BGCs are predicted as intact if the antiSMASH-annotated boundaries do not stop at a contig edge. (B) Proportion of intact BGCs per GCF in NPDC Actinobacteria genomes. Most NPDC GCFs consist only of intact BGCs, while >99% of NPDC GCFs contain at least one BGC predicted to be intact. BGCs are predicted as intact if the antiSMASH-annotated boundaries do not stop at a contig edge. By using GCFs in place of BGCs for measuring biosynthetic diversity, GCFs containing at least one putative intact BGC can minimize the impact of fragmented BGCs on downstream analyses.

**Extended Data Fig. 5.**
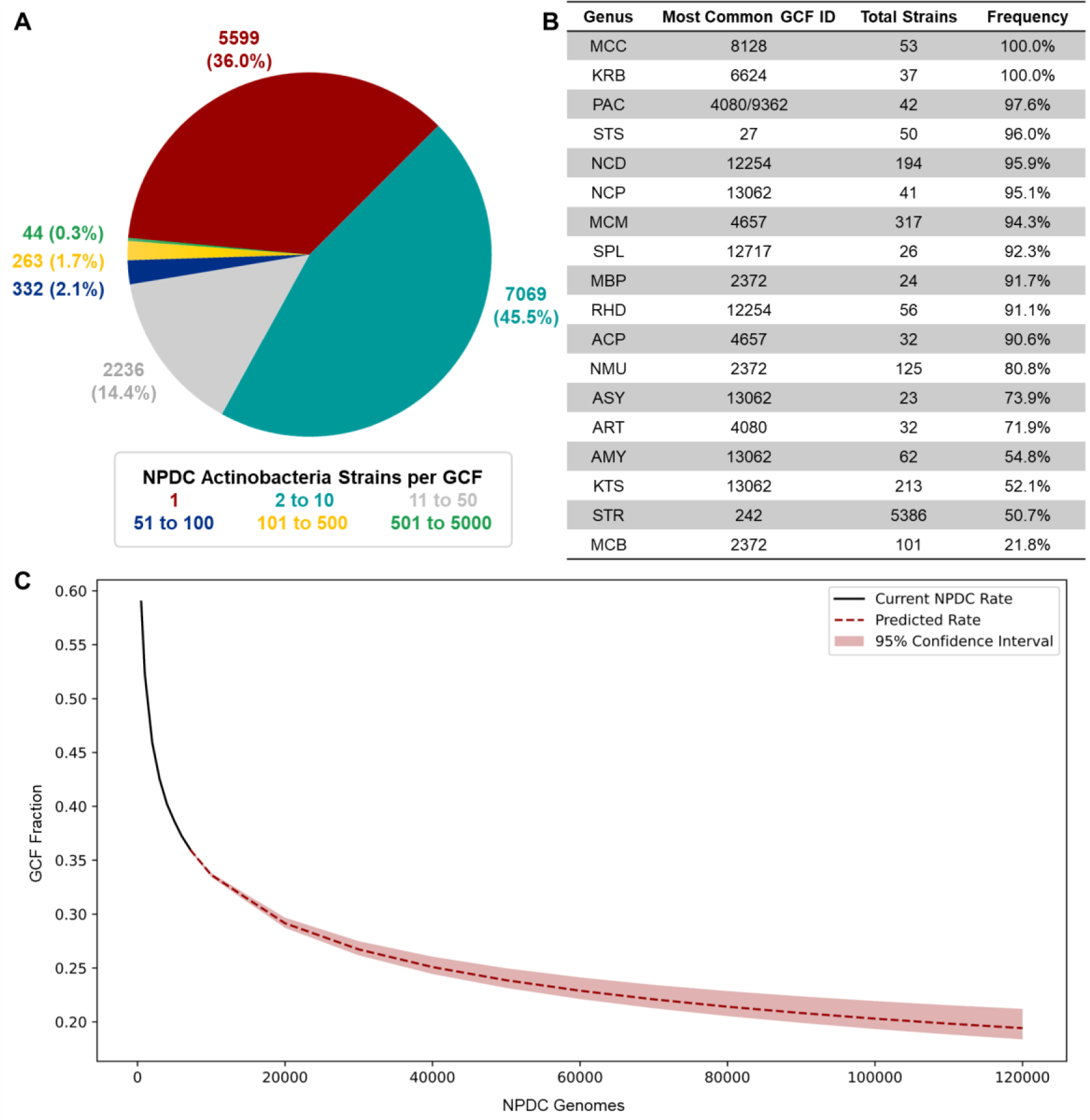
Distribution of NPDC Actinobacteria GCFs. (A) Distribution of GCFs were colored based on how many NPDC Actinobacteria genomes they were found in (e.g., GCFs with 1 strain = red), showing that most GCFs are found in ten or fewer NPDC genomes. (B) In genera with at least 20 strains, the number of instances of the most common GCF in each genus was listed by ID. In most genera, the most common GCF appears in >90% of genomes in that genus. In some cases, BGCs belonging to the most common GCF appear multiple times within a single genome, but these instances were only counted once. (C) A rarefaction analysis was performed for all NPDC Actinobacteria GCFs and extrapolated out to 120,000 genomes (approximately the size of the NPDC collection). The fraction of GCFs found in only a single genome are plotted here, indicating the value of continued genome sequencing for biosynthetic diversity. To determine the current NPDC rate, the fraction of Actinobacteria GCFs found in only a single genome was averaged at 500, 1000, 2000, 3000, 4000, 5000, 6000, and 7139 NPDC genomes (randomly sampled 100 times per number of genomes). See Supplementary Table 10 for genus abbreviations.

**Extended Data Fig. 6.**
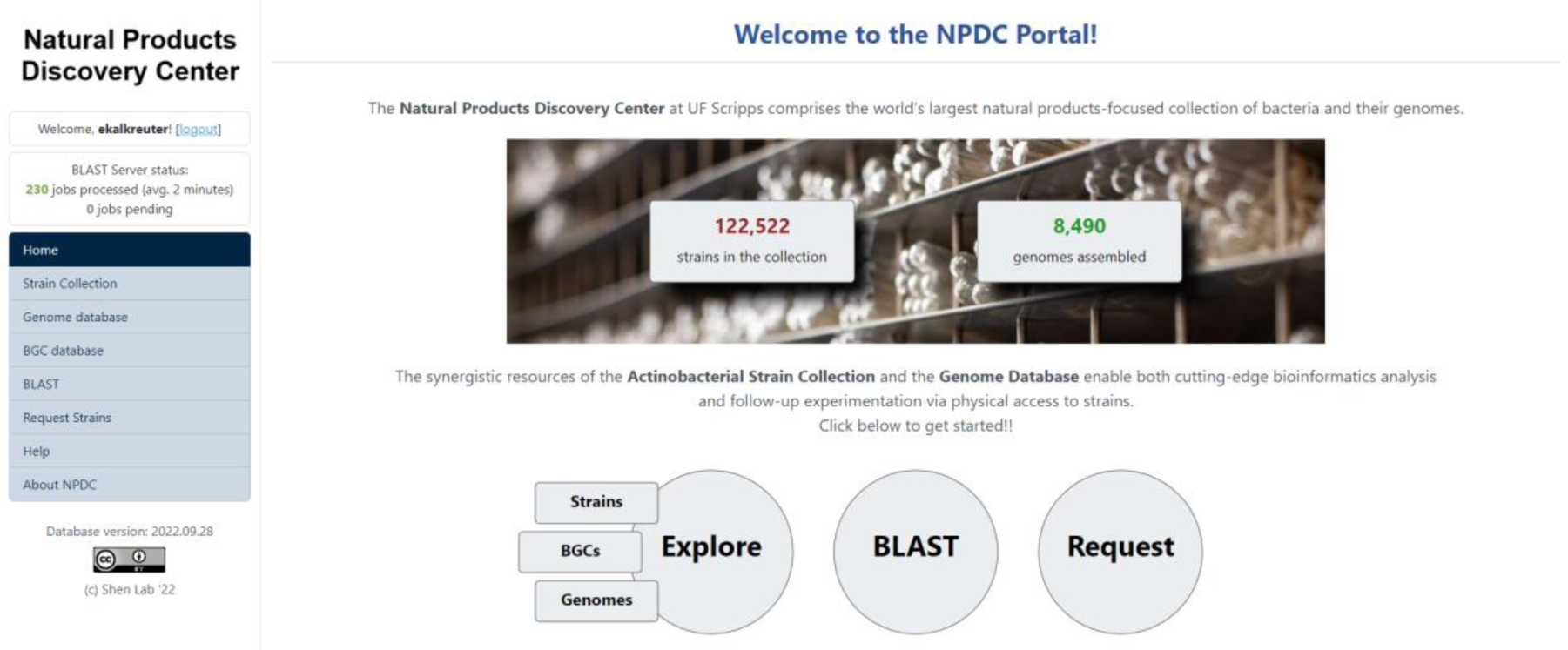
NPDC Portal home page. From the home page, users can access the strain, BGC, and genome databases, the BLASTP tool, and can request strains. Image pulled March 31, 2023.

**Extended Data Fig. 7.**
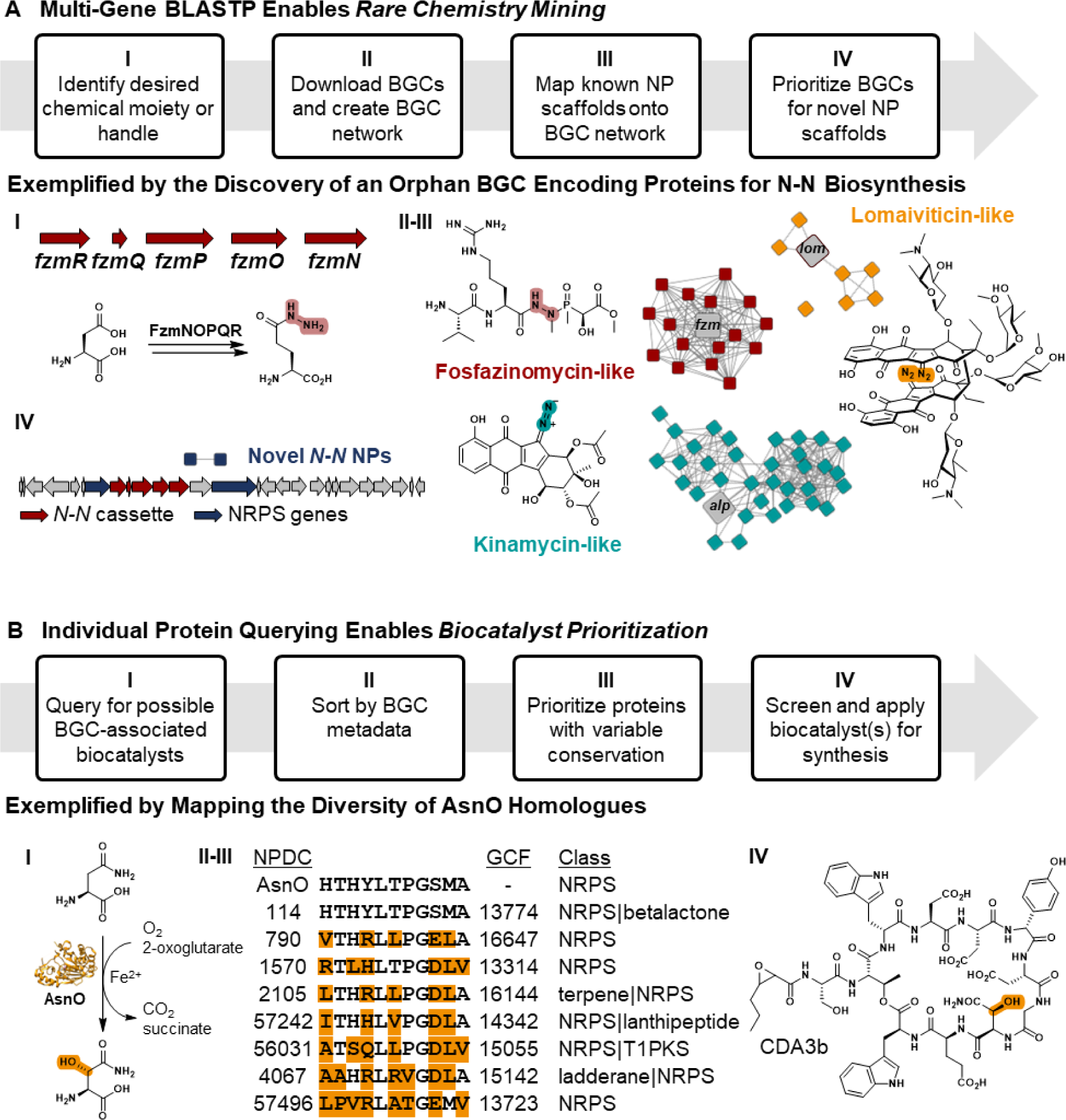
Genome mining using the NPDC Portal. (A) The NPDC Portal enables up to five simultaneous BLAST queries for NP diversity and novelty. (I) The five gene cassette required for *N*-*N* bond formation in fosfazinomycin was queried against the NPDC BGC database. (II) The resultant BGCs were clustered by BiG-SCAPE and (III) assigned to known and (IV) unknown NP families. ^45^ For the unknown family of BGCs containing this five-gene cassette, a representative BGC is depicted. Also see Supplementary Fig. 30. (B) The NPDC Portal enables searching enzyme families for biocatalyst discovery and development. (I) Sequences of BGC-associated α-KG-dependent dioxygenases related to the asparaginyl oxygenase AsnO (PDB:2OG5) were identified with the NPDC BLASTP tool. (II) The sequences were sorted by the NPDC-provided and antiSMASH-generated BGC classes and BiG-SLICE-generated GCFs and (III) aligned to identify homologues with varying conservation, allowing (IV) development of the optimal biocatalyst for synthetic applications. ^39, 47^

## Table of Contents Figure

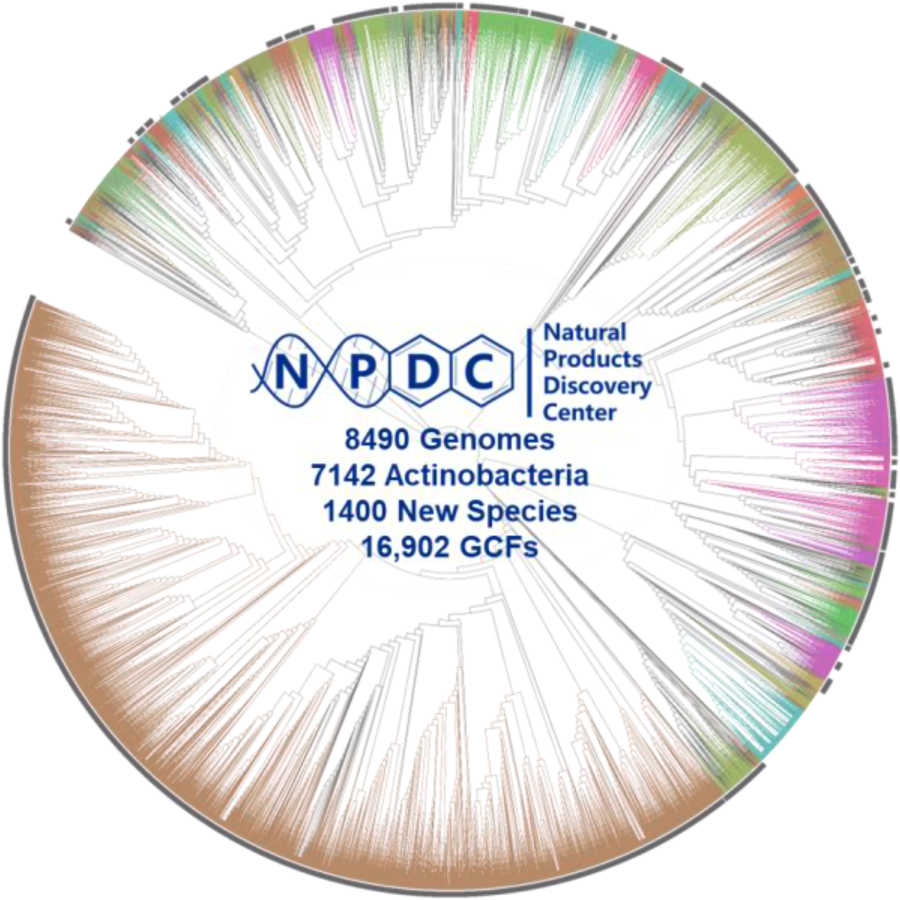

