## Supplementary Information for "The Natural Products Discovery Center: Release of the First 8490 Sequenced Strains for Exploring Actinobacteria Biosynthetic Diversity"

**This PDF file includes:**

Supplementary Tables 1 to 10
Supplementary Figures 1 to 39
SI References

**Supplementary Table 1.** Sample collection locations for sequenced NPDC strains with existing metadata (50% of sequenced strains).

| Country | Number of strains |
|---|---|
| Argentina | 3 |
| Australia | 1 |
| Belgium | 1 |
| Canada | 57 |
| Central African Republic | 22 |
| Chile | 33 |
| China | 3110 |
| Colombia | 13 |
| Comoros | 42 |
| Curaçao | 1 |
| Fiji | 62 |
| France | 6 |
| French Guiana | 1 |
| India | 5 |
| Indonesia | 226 |
| Italy | 50 |
| Malaysia | 6 |
| Paraguay | 2 |
| Peru | 8 |
| Portugal | 7 |
| Russia | 44 |
| Singapore | 2 |
| Solomon Islands | 10 |
| South Africa | 4 |
| Saint Lucia | 20 |
| Taiwan | 13 |
| Thailand | 1 |
| Trinidad and Tobago | 1 |
| Togo | 85 |
| United Arab Emirates | 1 |
| United Kingdom | 1 |
| United States of America | 398 |
| Uruguay | 13 |

**Supplementary Table 2.** Breakdown of 1348 non-Actinobacteria NPDC genomes from initial 8490 genomes.

| Phylum | Strains | Genera | Species (New) | Largest Genus | Strains | Species (New) |
|---|---|---|---|---|---|---|
| Firmicutes | 904 | 42 | 132 (45) | *Bacillus* | 427 | 11 (2) |
| Bacteroides | 25 | 3 | 7 (2) | *Pedobacter* | 19 | 1 (0) |
| Proteobacteria | 419 | 61 | 216 (57) | *Pseudomonas_E* | 152 | 80 (24) |

**Supplementary Table 3.** Comparison of historical morphology-based and sequence-based taxonomies of sequenced NPDC bacteria.

| Genus | Sequenced | Historical | Matched | Percent |
|---|---|---|---|---|
| *Actinoplanes* | 32 | 56 | 16 | 28.6% |
| *Actinosynnema* | 23 | 1 | 1 | 100.0% |
| *Amycolatopsis* | 62 | 19 | 8 | 42.1% |
| *Bacillus* | 609 | 4 | 1 | 25.0% |
| *Catenuloplanes* | 2 | 3 | 2 | 66.7% |
| *Dactylosporangium* | 12 | 15 | 5 | 33.3% |
| *Glycomyces* | 5 | 2 | 2 | 100.0% |
| *Kibdelosporangium* | 1 | 1 | 1 | 100.0% |
| *Kitasatospora* | 213 | 3 | 1 | 33.3% |
| *Kribbella* | 37 | 1 | 1 | 100.0% |
| *Lentzea* | 15 | 1 | 1 | 100.0% |
| *Microbispora* | 24 | 35 | 13 | 37.1% |
| *Micromonospora* | 317 | 173 | 135 | 78.0% |
| *Nocardia* | 194 | 113 | 21 | 18.6% |
| *Nocardioides* | 14 | 3 | 1 | 33.3% |
| *Nocardiopsis* | 41 | 1 | 1 | 100.0% |
| *Nonomuraea* | 125 | 3 | 2 | 66.7% |
| *Paenibacillus* | 52 | 1 | 1 | 100.0% |
| *Promicromonospora* | 12 | 1 | 1 | 100.0% |
| *Pseudonocardia* | 9 | 8 | 3 | 37.5% |
| *Rhodococcus* | 56 | 3 | 2 | 66.7% |
| *Saccharomonospora* | 5 | 4 | 2 | 50.0% |
| *Saccharopolyspora* | 14 | 11 | 7 | 63.6% |
| *Streptomyces* | 5386 | 842 | 774 | 91.9% |
| *Streptosporangium* | 50 | 50 | 10 | 20.0% |

**Supplementary Table 4.** FastANI of strains from new genera. [1]

| NPDC | 2781 | 48946 | 49639 | 59210 | 59280 |
|---|---|---|---|---|---|
| 2781 | - | - | - | - | - |
| 48946 | 98.23% | - | - | - | - |
| 49639 | < 80% | < 80% | - | - | - |
| 59210 | 82.66% | 82.64% | < 80% | - | - |
| 59280 | 82.70% | 82.60% | < 80% | 99.97% | - |

**Supplementary Table 5.** Number of antiSMASH-assigned MIBiG Actinobacteria BGC hits by threshold.

| MIBiG name | 100% hits | 80% hits | 50% hits |
|---|---|---|---|
| (-)-δ-cadinene | 36 | 36 | 39 |
| (+)-T-muurolol | 0 | 0 | 21 |
| 10,11-dihydro-8-deoxy-12,13-deepoxy-12,13-dihydrochalcomycin | 0 | 2 | 59 |
| 1-heptadecene | 11 | 11 | 11 |
| 1-nonadecene | 8 | 8 | 8 |
| 2-methylisoborneol | 1263 | 1263 | 1597 |
| 3,7-dihydroxytropolone | 0 | 0 | 85 |
| 4-hexadecanoyl-3-hydroxy-2-(hydroxymethyl)-2H-furan-5-one | 31 | 48 | 445 |
| 4-hydroxy-3-nitrosobenzamide | 0 | 0 | 19 |
| 4-Z-annimycin | 0 | 0 | 179 |
| 5-acetyl-5,10-dihydrophenazine-1-carboxylic acid | 25 | 34 | 166 |
| 5-isoprenylindole-3-carboxylate β-D-glycosyl ester | 0 | 0 | 486 |
| 67-121C | 0 | 0 | 103 |
| 6-methylsalicyclic acid | 1 | 1 | 1 |
| 7-prenylisatin | 41 | 52 | 111 |
| 9-methylstreptimidone | 0 | 0 | 73 |
| A33853 | 3 | 19 | 266 |
| A40926 | 0 | 0 | 410 |
| A-47934 | 1 | 5 | 31 |
| A-74528 | 0 | 1 | 11 |
| A-90289 A | 0 | 1 | 2 |
| A-94964 | 0 | 1 | 74 |
| aborycin | 80 | 86 | 105 |
| AbT1 | 2 | 2 | 2 |
| abyssomicin C | 0 | 0 | 41 |
| abyssomicin M | 0 | 2 | 18 |
| acarviostatin I03 | 0 | 0 | 151 |
| achromosin | 21 | 21 | 23 |
| aclacinomycin | 0 | 1 | 74 |
| actagardine | 0 | 0 | 44 |
| actinoallolide A | 0 | 0 | 135 |
| actinokineosin | 0 | 1 | 42 |
| actinomycin D | 0 | 17 | 29 |
| actinonin | 1 | 1 | 2 |
| actinorhodin | 23 | 31 | 86 |
| actinospectacin | 1 | 4 | 38 |
| aculeximycin | 0 | 0 | 51 |
| aerobactin | 1 | 1 | 2 |
| A-factor | 67 | 67 | 67 |

**Supplementary Table 5. (cont.)**

| MIBiG name | 100% hits | 80% hits | 50% hits |
|---|---|---|---|
| ajudazol A | 0 | 0 | 18 |
| alanylclavam | 1 | 3 | 336 |
| albachelin | 11 | 37 | 144 |
| albaflavenone | 2190 | 2190 | 2190 |
| albonoursin | 0 | 3 | 449 |
| albusnodin | 121 | 121 | 156 |
| aldgamycin J | 0 | 0 | 120 |
| alkyl-O-dihydrogeranyl-methoxyhydroquinones | 11 | 43 | 384 |
| alkylresorcinol | 2083 | 2083 | 2173 |
| allocyclinone | 0 | 0 | 194 |
| alnumycin A | 7 | 17 | 121 |
| altemicidin | 0 | 0 | 147 |
| althiomycin | 11 | 35 | 101 |
| AmfS | 595 | 773 | 821 |
| amicetin | 0 | 0 | 2 |
| amipurimycin | 0 | 22 | 171 |
| amonabactin P 750 | 0 | 0 | 3 |
| amychelin | 0 | 36 | 717 |
| amycolamycin A | 0 | 1 | 3 |
| amycomicin | 13 | 35 | 93 |
| anabaenopeptin NZ857 | 48 | 48 | 48 |
| anantin B1 | 0 | 0 | 80 |
| anantin C | 31 | 31 | 199 |
| angolamycin | 5 | 11 | 8 |
| anisomycin | 11 | 14 | 33 |
| ansamitocin P-3 | 0 | 2 | 45 |
| anthrabenzoxocinone | 6 | 6 | 9 |
| anthracimycin | 0 | 0 | 54 |
| anthramycin | 0 | 11 | 42 |
| antibiotic HKI 10311129 | 0 | 7 | 132 |
| antimycin | 380 | 499 | 595 |
| apoptolidin | 0 | 0 | 381 |
| apramycin | 0 | 0 | 18 |
| aranciamycin | 0 | 1 | 102 |
| arenimycin A | 0 | 1 | 34 |
| arimetamycin B | 0 | 1 | 42 |
| arixanthomycin A | 0 | 0 | 168 |
| arsono-polyketide | 5 | 42 | 306 |
| arylomycin | 10 | 24 | 198 |
| ashimides | 20 | 20 | 475 |
| atratumycin | 0 | 0 | 950 |

**Supplementary Table 5. (cont.)**

| MIBiG name | 100% hits | 80% hits | 50% hits |
|---|---|---|---|
| aurantimycin A | 1 | 1 | 67 |
| aureothin | 7 | 13 | 17 |
| auricin | 12 | 33 | 165 |
| auroramycin | 0 | 0 | 210 |
| avermectin | 0 | 2 | 4 |
| avermitilol | 265 | 265 | 265 |
| avilamycin A | 0 | 2 | 2 |
| avoparcin | 0 | 0 | 25 |
| azalomycin F3a | 4 | 6 | 94 |
| azinomycin B | 0 | 1 | 33 |
| azomycin | 0 | 10 | 26 |
| bacillibactin | 8 | 9 | 33 |
| bacillomycin D | 1 | 1 | 275 |
| bacilysin | 0 | 0 | 21 |
| bafilomycin B1 | 57 | 111 | 228 |
| bagremycin A | 0 | 0 | 193 |
| balhimycin | 0 | 1 | 327 |
| BD-12 | 0 | 2 | 79 |
| BE-14106 | 9 | 16 | 114 |
| BE-24566B | 3 | 3 | 5 |
| BE-43547A1 | 0 | 6 | 104 |
| BE-54017 | 0 | 2 | 12 |
| belactosin A | 0 | 0 | 338 |
| benastatin A | 5 | 7 | 15 |
| berninamycin A | 4 | 11 | 155 |
| bezastatin derivates | 60 | 69 | 75 |
| bicornutin A1 | 10 | 10 | 10 |
| bicyclomycin | 31 | 33 | 33 |
| bisucaberin B | 0 | 0 | 47 |
| blasticidin S | 0 | 0 | 190 |
| bleomycin | 0 | 3 | 28 |
| borrelidin | 0 | 0 | 2 |
| bottromycin A2 | 1 | 2 | 74 |
| bottromycin D | 0 | 0 | 9 |
| branched-chain fatty acids | 24 | 24 | 49 |
| butyrolactol A | 0 | 27 | 1 |
| C-1027 | 0 | 0 | 34 |
| caboxamycin | 28 | 70 | 105 |
| cacibiocin B | 0 | 0 | 19 |
| cadaside A | 0 | 0 | 114 |
| caerulomycin A | 0 | 0 | 57 |

**Supplementary Table 5. (cont.)**

| MIBiG name | 100% hits | 80% hits | 50% hits |
|---|---|---|---|
| calicheamicin | 0 | 0 | 21 |
| candicidin | 50 | 160 | 407 |
| caniferolide A | 0 | 0 | 1616 |
| capreomycin IA | 0 | 0 | 305 |
| carbapenem MM4550 | 0 | 0 | 31 |
| carotenoid | 64 | 66 | 1332 |
| catenulipeptin | 0 | 0 | 165 |
| catenulisporolides | 0 | 0 | 331 |
| cathomycin | 36 | 41 | 59 |
| cattlecin | 0 | 0 | 8 |
| CDA1b | 0 | 33 | 136 |
| celesticetin | 1 | 1 | 1 |
| cephamycin C | 0 | 0 | 82 |
| chartreusin | 1 | 1 | 19 |
| chaxapeptin | 0 | 0 | 108 |
| chloramphenicol | 21 | 38 | 212 |
| chlorizidine A | 0 | 0 | 3 |
| chlorothricin | 0 | 0 | 84 |
| chlortetracycline | 0 | 14 | 767 |
| chromomycin A3 | 8 | 9 | 45 |
| chrysomycin | 0 | 1 | 95 |
| chuangxinmycin | 0 | 0 | 34 |
| cinerubin B | 32 | 42 | 226 |
| cinnamycin | 0 | 0 | 484 |
| citrulassin A | 0 | 0 | 59 |
| citrulassin B | 33 | 42 | 66 |
| citrulassin D | 363 | 408 | 440 |
| citrulassin E | 4 | 4 | 10 |
| citrulassin F | 12 | 12 | 33 |
| clavulanic acid | 0 | 0 | 20 |
| clifednamide A | 0 | 1 | 1 |
| coelibactin | 278 | 314 | 418 |
| coelichelin | 892 | 1594 | 1855 |
| coelimycin P1 | 0 | 0 | 271 |
| colabomycin E | 0 | 1 | 13 |
| collinomycin | 0 | 31 | 143 |
| collismycin A | 0 | 33 | 6 |
| combamide | 2 | 2 | 47 |
| concanamycin A | 0 | 2 | 440 |
| cosmomycin D | 3 | 3 | 83 |
| coumermycin A1 | 1 | 11 | 69 |

**Supplementary Table 5. (cont.)**

| MIBiG name | 100% hits | 80% hits | 50% hits |
|---|---|---|---|
| cremimycin | 0 | 0 | 61 |
| curacomycin | 1 | 1 | 115 |
| curacozole | 0 | 2 | 7 |
| curamycin | 21 | 23 | 39 |
| cyclizidine | 0 | 2 | 185 |
| cycloheximide | 0 | 4 | 143 |
| cyclomarin D | 0 | 0 | 284 |
| cyclooctatin | 11 | 11 | 16 |
| cyclothiazomycin | 0 | 0 | 174 |
| cyclothiazomycin b1 | 2 | 2 | 8 |
| cyclothiazomycin C | 12 | 12 | 12 |
| cylindrospermopsin | 0 | 0 | 37 |
| cypemycin | 27 | 36 | 138 |
| cyphomycin | 0 | 0 | 209 |
| cyslabdan | 1 | 57 | 80 |
| cystargolide A | 0 | 3 | 75 |
| cysteoamide | 2 | 2 | 215 |
| cytorhodin | 0 | 28 | 191 |
| dactylocycline A | 1 | 1 | 535 |
| decaplanin | 1 | 2 | 38 |
| dehydrophos | 0 | 10 | 147 |
| deimino-antipain | 0 | 3 | 344 |
| desferrioxamin B | 1730 | 4106 | 4339 |
| desferrioxamine | 1 | 173 | 10 |
| desferrioxamine E | 452 | 452 | 522 |
| desosamine | 0 | 1 | 13 |
| desotamide | 11 | 13 | 38 |
| diazaquinomycin A | 2 | 2 | 4 |
| diazaquinomycin H | 0 | 16 | 82 |
| diazepinomicin | 0 | 0 | 481 |
| difficidin | 0 | 0 | 309 |
| diisonitrile antibiotic SF2768 | 0 | 0 | 25 |
| divergolide A | 1 | 9 | 60 |
| dynemicin A | 0 | 0 | 19 |
| E-837 | 12 | 18 | 54 |
| ebelactone | 0 | 0 | 25 |
| ebelactone A | 0 | 0 | 12 |
| echinomycin | 0 | 19 | 362 |
| echoside A | 24 | 69 | 406 |
| ECO-02301 | 0 | 1 | 40 |
| ectoine | 5106 | 5106 | 5586 |

**Supplementary Table 5. (cont.)**

| MIBiG name | 100% hits | 80% hits | 50% hits |
|---|---|---|---|
| elaiophylin | 0 | 12 | 49 |
| elloramycin | 1 | 1 | 22 |
| endophenazine A | 0 | 0 | 78 |
| enduracidin | 0 | 0 | 105 |
| enterocin | 9 | 30 | 114 |
| epilancin 15x | 0 | 0 | 2 |
| eponemycin | 0 | 0 | 99 |
| epoxomicin | 0 | 0 | 26 |
| erdacin | 0 | 0 | 3490 |
| erdasporine A | 0 | 0 | 8 |
| Ery-9 | 33 | 33 | 77 |
| erythrochelin | 0 | 19 | 615 |
| erythromycin A | 0 | 0 | 14 |
| esmeraldin | 1 | 4 | 111 |
| esperamicin | 0 | 0 | 41 |
| ethylenediaminesuccinic acid hydroxyarginine (EDHA) | 29 | 37 | 67 |
| FD-594 | 0 | 5 | 281 |
| FD-891 | 0 | 1 | 35 |
| feglymycin | 1 | 20 | 321 |
| fengycin | 0 | 5 | 1073 |
| filipin | 17 | 27 | 56 |
| flaviolin | 1 | 1 | 235 |
| fluostatin | 1 | 1 | 49 |
| fluostatins M-Q | 0 | 3 | 68 |
| fluvirucin B2 | 0 | 1 | 64 |
| fogacin A | 1 | 8 | 11 |
| formicamycins A-M | 0 | 3 | 369 |
| fosfazinomycin A | 3 | 46 | 168 |
| fosfomycin | 0 | 5 | 36 |
| fostriecin | 0 | 0 | 56 |
| foxicins A-D | 0 | 0 | 1 |
| FR-900098 | 0 | 1 | 11 |
| FR900359 | 0 | 0 | 3 |
| FR-900520 | 0 | 0 | 76 |
| frankiamicin | 0 | 0 | 508 |
| fredericamycin A | 0 | 26 | 2 |
| frenolicin B | 0 | 7 | 91 |
| frigocyclinone | 0 | 0 | 88 |
| friulimicin A | 0 | 0 | 353 |
| frontalamide B | 0 | 25 | 66 |
| funisamine | 0 | 0 | 1 |

**Supplementary Table 5. (cont.)**

| MIBiG name | 100% hits | 80% hits | 50% hits |
|---|---|---|---|
| fuscachelin A | 3 | 10 | 58 |
| fusilassin | 0 | 0 | 13 |
| GE2270 | 0 | 0 | 72 |
| GE37468 | 2 | 5 | 13 |
| geldanamycin | 1 | 1 | 78 |
| geosmin | 4323 | 4323 | 4354 |
| germicidin | 589 | 589 | 589 |
| gilvocarcin V | 0 | 20 | 68 |
| gilvusmycin | 0 | 4 | 32 |
| glycinocin A | 0 | 0 | 165 |
| glycopeptidolipid | 0 | 3 | 1 |
| gobichelin A | 0 | 0 | 31 |
| granaticin | 0 | 4 | 256 |
| grincamycin | 0 | 6 | 6 |
| griseochelin | 1 | 48 | 27 |
| griseorhodin A | 29 | 37 | 74 |
| griseoviridin | 0 | 0 | 121 |
| grixazone | 0 | 0 | 67 |
| guadinomine | 0 | 0 | 29 |
| guangnanmycin | 9 | 29 | 200 |
| halstoctacosanolide A | 1 | 1 | 371 |
| hatomarubigin A | 0 | 1 | 101 |
| heat-stable antifungal factor | 0 | 0 | 10 |
| hedamycin | 0 | 39 | 306 |
| herbimycin A | 0 | 0 | 23 |
| herboxidiene | 0 | 0 | 15 |
| heronamide A | 0 | 0 | 369 |
| heterobactin A | 9 | 20 | 43 |
| himastatin | 0 | 0 | 94 |
| hiroshidine | 2 | 5 | 100 |
| hitachimycin | 0 | 1 | 84 |
| holomycin | 0 | 13 | 167 |
| hopene | 249 | 1889 | 6353 |
| hormaomycin | 0 | 0 | 395 |
| huanglongmycin A | 0 | 0 | 44 |
| hydroxystreptomycin | 0 | 2 | 36 |
| hygrocin A | 0 | 3 | 185 |
| hygromycin A | 0 | 5 | 44 |
| α,β-epoxyketone | 3 | 4 | 30 |
| α-lipomycin | 1 | 1 | 305 |
| β-D-galactosylvalidoxylamine-A | 0 | 0 | 2 |

**Supplementary Table 5. (cont.)**

| MIBiG name | 100% hits | 80% hits | 50% hits |
|---|---|---|---|
| ibomycin | 0 | 0 | 66 |
| icosalide A | 104 | 104 | 104 |
| ikarugamycin | 0 | 8 | 164 |
| ilamycins | 0 | 6 | 120 |
| indigoidine | 15 | 135 | 330 |
| informatipeptin | 398 | 480 | 2089 |
| isatropolone A | 4 | 8 | 45 |
| ishigamide | 37 | 76 | 349 |
| isocomplestatin | 24 | 86 | 131 |
| isofuranonaphthoquinone | 6 | 7 | 21 |
| isoindolinomycin | 1 | 2 | 47 |
| iso-migrastatin | 21 | 22 | 25 |
| isorenieratene | 1648 | 2142 | 3676 |
| jadomycin | 3 | 25 | 45 |
| jawsamycin | 0 | 1 | 109 |
| JBIR-06 | 0 | 0 | 174 |
| JBIR-100 | 0 | 4 | 2 |
| JBIR-126 | 72 | 148 | 300 |
| JBIR-34 | 0 | 6 | 332 |
| JBIR-76 | 0 | 2 | 339 |
| JBIR-78 | 0 | 3 | 166 |
| jinggangmycin | 5 | 5 | 5 |
| julichrome Q3-3 | 1 | 1 | 11 |
| kanamycin | 0 | 1 | 11 |
| kasugamycin | 0 | 0 | 3 |
| kedarcidin | 0 | 0 | 29 |
| kendomycin | 0 | 1 | 262 |
| keratinimicin A | 0 | 1 | 45 |
| ketomemicin B3 | 6 | 34 | 60 |
| keywimysin | 436 | 446 | 449 |
| kiamycin | 0 | 1 | 68 |
| kijanimicin | 1 | 1 | 35 |
| kinamycin | 0 | 5 | 104 |
| kirromycin | 0 | 0 | 747 |
| kistamicin A | 0 | 0 | 425 |
| kitasetaline | 0 | 0 | 8 |
| komodoquinone B | 15 | 19 | 136 |
| kosinostatin | 0 | 0 | 28 |
| labyrinthopeptin A2 | 2 | 2 | 119 |
| lactazole | 1 | 1 | 232 |
| lactimidomycin | 0 | 0 | 10 |

**Supplementary Table 5. (cont.)**

| MIBiG name | 100% hits | 80% hits | 50% hits |
|---|---|---|---|
| laidlomycin | 0 | 0 | 38 |
| landomycin A | 3 | 14 | 90 |
| lankacidin C | 5 | 5 | 7 |
| lankamycin | 2 | 3 | 11 |
| lasalocid | 0 | 0 | 121 |
| lavendiol | 0 | 1 | 159 |
| lazarimide A | 0 | 0 | 1 |
| legonaridin | 0 | 0 | 84 |
| leinamycin | 0 | 3 | 261 |
| lichenysin | 0 | 1 | 6 |
| lidamycin | 4 | 21 | 242 |
| limazepine C | 0 | 0 | 240 |
| lincomycin | 0 | 0 | 35 |
| linfuranone B | 0 | 0 | 421 |
| lipopeptide 8D1-1 | 0 | 7 | 35 |
| liposidomycin B | 0 | 7 | 20 |
| lipstatin | 1 | 1 | 551 |
| lividomycin | 0 | 1 | 90 |
| livipeptin | 87 | 87 | 156 |
| LL-D49194α1 (LLD) | 0 | 2 | 141 |
| LL-F28249α | 0 | 0 | 27 |
| lobophorin A | 0 | 0 | 636 |
| lobophorin B | 0 | 0 | 174 |
| lobosamide A | 0 | 1 | 150 |
| lomaiviticin A | 0 | 0 | 730 |
| lomofungin | 0 | 3 | 1886 |
| LP2006 | 15 | 15 | 23 |
| lugdunomycin | 0 | 6 | 7 |
| luminmide | 5 | 5 | 5 |
| lydicamycin | 4 | 14 | 173 |
| lysolipin I | 0 | 19 | 1355 |
| macbecin | 1 | 2 | 175 |
| macrotermycins | 0 | 5 | 687 |
| macrotetrolide | 0 | 1 | 48 |
| maduropeptin | 0 | 0 | 1234 |
| maklamicin | 0 | 0 | 266 |
| malonomycin | 2 | 9 | 45 |
| mannopeptimycin | 0 | 10 | 114 |
| marformycin A | 0 | 0 | 270 |
| marineosin A | 9 | 23 | 161 |
| marinopyrrole A | 0 | 0 | 82 |

**Supplementary Table 5. (cont.)**

| MIBiG name | 100% hits | 80% hits | 50% hits |
|---|---|---|---|
| matlystatin A | 0 | 0 | 110 |
| mayamycin | 2 | 6 | 125 |
| medermycin | 0 | 1 | 549 |
| mediomycin A | 0 | 2 | 161 |
| melanin | 1392 | 1483 | 3687 |
| mensacaricin | 0 | 1 | 8 |
| meoabyssomicin | 0 | 0 | 94 |
| meridamycin | 0 | 1 | 161 |
| merochlorin A | 0 | 5 | 62 |
| mersacidin | 0 | 1 | 1 |
| metatricycloene | 0 | 0 | 88 |
| methylenomycin A | 8 | 8 | 155 |
| methylpendolmycin | 0 | 0 | 3 |
| methymycin | 0 | 0 | 27 |
| michiganin A | 0 | 1 | 2 |
| microansamycin | 0 | 0 | 101 |
| microbisporicin A2 | 0 | 0 | 14 |
| microtermolide A | 0 | 0 | 577 |
| miharamycin A | 3 | 30 | 17 |
| mirubactin | 0 | 1 | 66 |
| mithramycin | 0 | 11 | 100 |
| mitomycin | 0 | 0 | 30 |
| ML-449 | 0 | 0 | 209 |
| moenomycin | 2 | 2 | 5 |
| monensin | 0 | 2 | 459 |
| monobactam | 0 | 1 | 2 |
| moomysin | 0 | 0 | 906 |
| MS-271 | 39 | 41 | 72 |
| murayaquinone | 0 | 1 | 589 |
| muraymycin C1 | 0 | 0 | 75 |
| mycinamicin II | 0 | 0 | 85 |
| mycobactin | 0 | 3 | 43 |
| mycotrienin I | 0 | 0 | 346 |
| myxochelin A | 0 | 0 | 59 |
| myxothiazol | 0 | 0 | 6 |
| nanchangmycin | 1 | 5 | 288 |
| naphthomycin A | 0 | 0 | 188 |
| naphthyridinomycin | 23 | 48 | 190 |
| napsamycin A | 0 | 12 | 143 |
| napyradiomycin | 0 | 0 | 138 |
| nargenicin | 0 | 0 | 110 |

**Supplementary Table 5. (cont.)**

| MIBiG name | 100% hits | 80% hits | 50% hits |
|---|---|---|---|
| naseseazine C | 1 | 1 | 8 |
| natamycin | 0 | 1 | 131 |
| nataxazole | 0 | 1 | 488 |
| nenestatin | 0 | 0 | 4 |
| neoantimycin | 0 | 1 | 86 |
| neocarazostatin A | 18 | 28 | 39 |
| neocarzilin A | 6 | 28 | 135 |
| neocarzinostatin | 0 | 3 | 90 |
| neomycin | 0 | 1 | 219 |
| neopolyoxin C | 16 | 16 | 33 |
| neothioviridamide | 12 | 14 | 26 |
| netropsin | 19 | 44 | 96 |
| niddamycin | 0 | 0 | 1 |
| nigericin | 5 | 58 | 147 |
| niphimycins C-E | 0 | 2 | 309 |
| nocamycin | 0 | 0 | 130 |
| nocardicin A | 1 | 1 | 5 |
| nocobactin NA | 19 | 120 | 192 |
| nogabecin | 0 | 0 | 11 |
| nogalamycin | 0 | 0 | 357 |
| nonactin | 0 | 2 | 32 |
| nosiheptide | 0 | 7 | 29 |
| nostopeptolide A2 | 0 | 0 | 182 |
| nucleocidin | 0 | 1 | 630 |
| nybomycin | 0 | 0 | 23 |
| nystatin | 0 | 0 | 417 |
| nystatin A1 | 0 | 16 | 100 |
| nystatin-like Pseudonocardia polyene | 3 | 3 | 39 |
| ochronotic pigment | 2 | 2 | 92 |
| oligomycin | 8 | 15 | 127 |
| olimycin A | 8 | 19 | 137 |
| oronofacic acid | 1 | 1 | 1 |
| oviedomycin | 0 | 6 | 71 |
| oxalomycin B | 0 | 1 | 160 |
| oxazolepoxidomycin A | 0 | 7 | 168 |
| oxytetracycline | 0 | 25 | 111 |
| pacidamycin 1 | 0 | 0 | 61 |
| pactamides | 4 | 5 | 122 |
| pactamycin | 0 | 1 | 45 |
| paenibactin | 0 | 235 | 21 |
| paerucumarin | 0 | 0 | 60 |

**Supplementary Table 5. (cont.)**

| MIBiG name | 100% hits | 80% hits | 50% hits |
|---|---|---|---|
| paromomycin | 0 | 11 | 335 |
| paulomycin | 0 | 0 | 33 |
| pellasoren | 0 | 0 | 56 |
| pentalenolactone | 17 | 24 | 286 |
| pentamycin | 16 | 37 | 98 |
| pepticinnamin E | 34 | 51 | 338 |
| perquinoline A | 1 | 8 | 95 |
| petrobactin | 2 | 2 | 3 |
| pheganomycin | 0 | 4 | 29 |
| phenalamide | 0 | 0 | 9 |
| phenalamide A2 | 0 | 0 | 7 |
| phenalinolactone A | 0 | 11 | 510 |
| phoslactomycin B | 5 | 9 | 42 |
| phosphinothricintripeptide | 0 | 4 | 119 |
| phosphonoacetic Acid | 3 | 6 | 34 |
| physostigmine | 0 | 1 | 1 |
| piericidin A1 | 9 | 32 | 126 |
| pimaricin | 0 | 0 | 210 |
| pladienolide B | 0 | 0 | 37 |
| planosporicin | 76 | 81 | 122 |
| polyketomycin | 0 | 2 | 268 |
| polyoxin A | 0 | 0 | 403 |
| polyoxypeptin | 0 | 9 | 160 |
| porothramycin A | 0 | 0 | 25 |
| prejadomycin | 0 | 7 | 137 |
| PreQ0 Base | 6 | 6 | 6 |
| pristinol | 161 | 161 | 161 |
| pseudomonine | 1 | 1 | 1 |
| pseudouridimycin | 0 | 1 | 367 |
| purincyclamide | 11 | 11 | 43 |
| puromycin | 4 | 12 | 16 |
| putisolvin | 0 | 0 | 2 |
| pyocyanine | 7 | 7 | 13 |
| pyralomicin 1a | 0 | 1 | 303 |
| pyridomycin | 0 | 0 | 318 |
| pyrrolizixenamide A | 3 | 3 | 3 |
| qinichelins | 0 | 8 | 1 |
| quartromicin A1 | 0 | 0 | 122 |
| quinocarcin | 0 | 0 | 120 |
| radamycin | 3 | 25 | 91 |
| raimonol | 33 | 138 | 152 |

**Supplementary Table 5. (cont.)**

| MIBiG name | 100% hits | 80% hits | 50% hits |
|---|---|---|---|
| ralsolamycin | 0 | 0 | 42 |
| rapamycin | 0 | 0 | 93 |
| ravidomycin | 0 | 4 | 63 |
| reductasporine | 3 | 5 | 86 |
| resistomycin | 0 | 36 | 236 |
| rhizomide A | 237 | 237 | 237 |
| rhodochelin | 13 | 13 | 14 |
| rifamorpholine A | 0 | 0 | 74 |
| rifamycin | 0 | 0 | 253 |
| rimocidin | 0 | 2 | 71 |
| rimosamide | 13 | 21 | 742 |
| rishirilide B | 10 | 12 | 55 |
| ristocetin | 1 | 1 | 6 |
| ristomycin A | 4 | 5 | 19 |
| rosamicin | 0 | 0 | 131 |
| RP-1776 | 0 | 42 | 66 |
| rubradirin | 1 | 1 | 11 |
| rubrolone A | 0 | 0 | 97 |
| rufomycin | 0 | 0 | 9 |
| s56-p1 | 0 | 1 | 87 |
| saalfelduracin | 0 | 1 | 2 |
| saframycin A | 0 | 0 | 60 |
| SAL-2242 | 105 | 106 | 234 |
| salinamide A | 10 | 13 | 173 |
| salinichelins | 0 | 0 | 662 |
| salinilactam | 0 | 2 | 184 |
| salinipostin G | 0 | 0 | 2 |
| salinomycin | 8 | 8 | 562 |
| salinosporamide A | 0 | 0 | 49 |
| SapB | 926 | 926 | 1176 |
| saprolmycin E | 0 | 5 | 407 |
| saquayamycin A | 0 | 0 | 107 |
| saquayamycin Z | 0 | 2 | 91 |
| sarpeptin A | 3 | 26 | 13 |
| SBI-06990 A1 | 45 | 45 | 128 |
| scabichelin | 252 | 301 | 414 |
| sceliphrolactam | 3 | 137 | 633 |
| Sch 18640 | 5 | 5 | 44 |
| Sch-47554 | 0 | 2 | 118 |
| scleric acid | 0 | 3 | 18 |
| SCO-2138 | 1 | 134 | 29 |

**Supplementary Table 5. (cont.)**

| MIBiG name | 100% hits | 80% hits | 50% hits |
|---|---|---|---|
| setomimycin | 11 | 12 | 36 |
| SF2575 | 0 | 0 | 14 |
| SGR PTMs | 928 | 969 | 979 |
| showdomycin | 0 | 0 | 349 |
| siamycin | 0 | 0 | 3 |
| siamycin I | 0 | 17 | 102 |
| sibiromycin | 0 | 6 | 14 |
| simocyclinone D8 | 0 | 2 | 23 |
| siomycin A | 3 | 6 | 64 |
| sipanmycin | 0 | 0 | 377 |
| spectinabilin | 2 | 12 | 42 |
| spectinomycin | 0 | 0 | 9 |
| sphaericin | 0 | 0 | 2 |
| spicamycin | 0 | 0 | 19 |
| spiramycin | 0 | 0 | 90 |
| spore pigment | 9 | 2204 | 3679 |
| sporolide A | 0 | 0 | 276 |
| SRO15-2005 | 0 | 2 | 24 |
| SRO15-3108 | 64 | 64 | 71 |
| SSV-2083 | 0 | 2 | 34 |
| stambomycin A | 3 | 3 | 133 |
| staphyloferrin A | 3 | 3 | 3 |
| staurosporine | 19 | 36 | 59 |
| steffimycin D | 0 | 0 | 266 |
| stenothricin | 0 | 11 | 14 |
| streptazone E | 0 | 18 | 361 |
| streptobactin | 54 | 392 | 1083 |
| streptolydigin | 0 | 22 | 802 |
| streptomonomicin | 0 | 7 | 21 |
| streptomycin | 0 | 1 | 1230 |
| streptophenazine B | 14 | 35 | 115 |
| streptothricin | 30 | 110 | 168 |
| streptovaricin | 0 | 0 | 414 |
| surugamide A | 61 | 100 | 454 |
| SWA-2138 | 0 | 0 | 16 |
| syringopeptin 25A | 1 | 1 | 3 |
| tacrolimus | 0 | 0 | 4 |
| tallysomycin A | 0 | 0 | 110 |
| tautomycetin | 0 | 1 | 82 |
| tautomycin | 0 | 0 | 10 |
| teichomycin | 0 | 11 | 690 |

**Supplementary Table 5. (cont.)**

| MIBiG name | 100% hits | 80% hits | 50% hits |
|---|---|---|---|
| telomestatin | 23 | 40 | 92 |
| telomycin | 0 | 0 | 318 |
| tetracenomycin C | 10 | 11 | 15 |
| tetrocarcin A | 0 | 0 | 92 |
| thiazostatin | 16 | 29 | 148 |
| thiocoraline | 0 | 0 | 59 |
| thioholgamide A | 3 | 3 | 64 |
| thiolactomycin | 0 | 2 | 17 |
| thiomuracin | 0 | 0 | 13 |
| thiomuracin A | 5 | 6 | 9 |
| thioplatensimycin | 0 | 0 | 399 |
| thiostrepton | 2 | 2 | 4 |
| thiotetroamide | 0 | 3 | 9 |
| thiovarsolin A | 2 | 3 | 20 |
| tiacumicin B | 0 | 0 | 73 |
| tiancilactone | 1 | 6 | 43 |
| tiancimycin | 0 | 11 | 232 |
| tirandamycin | 1 | 25 | 168 |
| TLN-05220 | 0 | 1 | 58 |
| tolaasin I | 0 | 0 | 6 |
| tomaymycin | 30 | 44 | 56 |
| totopotensamide A | 0 | 1 | 96 |
| toxoflavin | 0 | 0 | 97 |
| toyocamycin | 18 | 57 | 244 |
| TP-1161 | 0 | 0 | 71 |
| trehangelin | 0 | 0 | 4 |
| triacsins | 0 | 0 | 25 |
| triostin A | 2 | 2 | 58 |
| tubercidin | 7 | 32 | 57 |
| tunicamycin B1 | 0 | 13 | 82 |
| tylactone | 0 | 0 | 317 |
| tyrobetaine | 0 | 35 | 57 |
| ulleungmycin | 0 | 4 | 757 |
| uncialamycin | 0 | 1 | 51 |
| undecylprodigiosin | 77 | 128 | 184 |
| valclavam | 0 | 3 | 161 |
| validamycin A | 0 | 0 | 205 |
| valinomycin | 0 | 53 | 25 |
| vancosamine | 2 | 3 | 3 |
| vazabitide A | 0 | 0 | 70 |
| venemycin | 2 | 3 | 32 |

**Supplementary Table 5. (cont.)**

| MIBiG name | 100% hits | 80% hits | 50% hits |
|---|---|---|---|
| versipelostatin | 0 | 0 | 15 |
| vicenistatin | 0 | 5 | 40 |
| viguiepinol | 4 | 4 | 165 |
| violapyrone B | 0 | 0 | 48 |
| viomycin | 34 | 46 | 74 |
| virginiamycin S1 | 0 | 38 | 689 |
| warkmycin CS1 | 0 | 84 | 11 |
| weishanmycin | 0 | 10 | 127 |
| WS79089A | 1 | 10 | 616 |
| WS9326 | 0 | 31 | 46 |
| X-14547 | 0 | 0 | 56 |
| xantholipin | 0 | 2 | 1031 |
| xanthothricin | 0 | 0 | 6 |
| xenematide | 8 | 8 | 8 |
| xenotetrapeptide | 10 | 10 | 10 |
| xiamycin A | 0 | 0 | 520 |
| zorbamycin | 1 | 3 | 46 |
| zwittermicin A | 0 | 0 | 7 |

**Supplementary Table 6.** Comparison of observed bonnevillamide A chemical shifts in MeOD-$d_4$ ($^1$H NMR at 600 MHz and $^{13}$C NMR at 151 MHz) in comparison to values previously reported in DMSO-$d_6$. Optical rotation was determined as $[\propto]_D^{23}$ -5 ° g$^{-1}$ mL dm$^{-1}$ (c = 0.1, DMSO) compared to the literature reported $[\propto]_D^{20}$ -30 ° g$^{-1}$ mL dm$^{-1}$ (c = 0.1, $CHCl_3$). [2]

| Amino-acid | # | $^1$H observed [ppm] | $^1$H literature[2] [ppm] | $^{13}$C observed [ppm] | $^{13}$C literature[2] [ppm] | Δ $^1$H | Δ $^{13}$C |
|---|---|---|---|---|---|---|---|
| *BVA* | **1** | | | 166.8 | 163.1 | | 3.7 |
| | **2** | | | 151.8 | 149.5 | | 2.3 |
| | **3** | 6.74 | 6.66 | 119.5 | 115.9 | 0.08 | 3.6 |
| | **4** | | | 128.4 | 127.0 | | 1.4 |
| | **5/5'** | 7.67 | 7.73 | 131.6 | 129.8 | 0.03 | 1.8 |
| | **6/6'** | | | 124.3 | 122.7 | | 1.6 |
| | **7** | | | 150.7 | 149.4 | | 1.3 |
| | **7-OMe** | 3.73 | 3.67 | 60.8 | 60.1 | 0.06 | 0.7 |
| *Thr* | **1** | | | 172.9 | 170.2 | | 2.7 |
| | **2** | 4.60 | 4.26 | 61.1 | 59.1 | 0.34 | 2.0 |
| | **3** | 4.16 | 3.93 | 69.4 | 67.2 | 0.24 | 2.2 |
| | **4** | 1.19 | 1.05 | 21.3 | 20.8 | 0.14 | 0.5 |
| *Leu$_1$* | **1** | | | 176.3 | 173.8 | | 2.5 |
| | **2** | 5.09 | 4.98 | 50.7 | 48 | 0.13 | 2.7 |
| | **3** | 1.56 | 1.37, 1.52 | 42 | 40.1 | 0.04 | 1.9 |
| | **4** | 1.79 | 1.66 | 26.7 | 24.4 | 0.12 | 2.3 |
| | **5** | 0.94 | 0.83 | 22.8 | 22.1 | 0.11 | 0.7 |
| | **6** | 0.98 | 0.83 | 24.7 | 22.4 | 0.15 | 2.3 |
| *HMPro* | **1** | | | 173.5 | 170.7 | | 2.8 |
| | **2** | 4.41 | 4.43 | 60.8 | 58.4 | 0.02 | 2.4 |
| | **3** | 2.35 | 2.15 | 33.9 | 32.1 | 0.20 | 1.8 |
| | **4** | 4.98 | 4.88 | 80.7 | 78.5 | 0.10 | 2.2 |
| | **5** | 4.66 | 4.49 | 62.8 | 60.1 | 0.17 | 2.7 |
| | **6** | 1.34 | 1.18 | 20.2 | 19.1 | 0.16 | 1.1 |
| | **7** | | | 173.3 | 170.5 | | 2.8 |
| | **8** | 2.08 | 1.99 | 22 | 21.3 | 0.09 | 0.7 |
| *Leu$_2$* | **1** | | | 173.8 | 171.2 | | 2.6 |
| | **2** | 4.64 | 4.49 | 51 | 48.6 | 0.15 | 2.4 |
| | **3** | 1.50, 1.77 | 1.31, 1.62 | 42.7 | 41.3 | 0.19, 0.15 | 1.4 |
| | **4** | 1.75 | 1.62 | 26.6 | 24.7 | 0.13 | 1.9 |
| | **5** | 0.95 | 0.82 | 22.4 | 21.5 | 0.13 | 0.9 |
| | **6** | 0.98 | 0.88 | 24.6 | 23.7 | 0.10 | 0.9 |
| *N-OH-Val* | **1** | | | 171.7 | 168.3 | | 3.4 |
| | **2** | 4.57 | 4.43 | 63.5 | 61.2 | 0.12 | 2.3 |
| | **3** | 2.46 | 2.31 | 28.2 | 26.1 | 0.15 | 2.1 |
| | **4** | 0.92 | 0.77 | 20.2 | 18.9 | 0.15 | 1.3 |
| | **5** | 0.98 | 0.84 | 20.5 | 19.7 | 0.14 | 0.8 |
| *MACME* | **1** | | | 174.8 | 171.6 | | 3.2 |
| | **2** | 4.64 | 4.49 | 58.5 | 56.2 | 0.15 | 2.3 |
| | **3** | 1.84, 2.77 | 1.72, 2.69 | 29.6 | 27.9 | 0.08, 0.08 | 1.7 |
| | **4** | 4.59 | 4.36 | 60.1 | 57.3 | 0.23 | 2.8 |
| | **5** | 1.59 | 1.47 | 23.5 | 22.4 | 0.12 | 1.1 |
| | **6** | 3.75 | 3.67 | 53.6 | 52.4 | 0.08 | 1.2 |

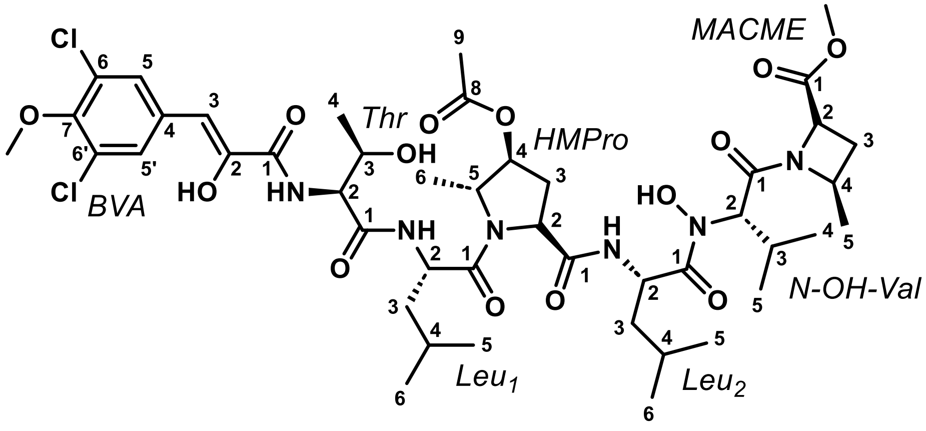

MACME
Thr
HMPro
BVA
N-OH-Val
Leu1
Leu2
Cl
Cl
OH
HO
HN
NH
N
H
HO

**Supplementary Table 7.** Annotation of *esp* BGC from *Spirillospora* sp. NPDC050156.

| Gene | AA | Putative function | CAL homologue | % Identity/ Similarity | Protein homologue | % Identity/ Similarity |
|---|---|---|---|---|---|---|
| *espU1* | 400 | Hypothetical protein | CalU10 | 70/82 | WP_185024253.1 | 100/100 |
| *espE1* | 294 | SAM-dependent methyltransferase | CalE3 | 63/72 | MBB6394621.1 | 97/98 |
| *espU2* | 222 | Hypothetical protein | CalU9 | 43/63 | WP_089315999.1 | 96/99 |
| *espR1* | 444 | AarF/UbiB kinase family protein | CalR3 | 63/74 | WP_185024256.1 | 95/96 |
| *espR2* | 119 | DNA-binding ArsR family transcriptional regulator | CalR8 | 67/82 | MBB6394624.1 | 95/97 |
| *espG1* | 383 | Glycosyltransferase | CalG3 | 51/66 | WP_185024258.1 | 98/98 |
| *espC* | 159 | Hypothetical protein | -- | -- | MCR3742863.1 | 92/95 |
| *espU3* | 162 | Hypothetical protein | -- | -- | WP_179834670.1 | 92/98 |
| *espT1* | 278 | ABC transporter permease | CalT4 | 58/73 | GGQ36575.1 | 97/98 |
| *espA1* | 303 | Arylamine *N*-acetyltransferase | -- | -- | WP_185024261.1 | 90/92 |
| *espS1* | 351 | Glucose-1-phosphate thymidylyltransferase | CalS7 | 38/54 | WP_179834673.1 | 93/96 |
| *espS2* | 197 | NDP-sugar epimerase | CalS1 | 45/59 | WP_258943028.1 | 92/94 |
| *espU4* | 242 | FkbM family methyltransferase | -- | -- | WP_185024624.1 | 96/97 |
| *espS3* | 500 | B12-binding domain-containing radical SAM protein | CalU22 | 79/89 | WP_185024265.1 | 94/97 |
| *espR3* | 216 | DNA-binding HxlR family transcriptional regulator | CalU8 | 52/64 | MBB6394634.1 | 98/99 |
| *espU5* | 155 | Hypothetical protein | -- | -- | WP_185024267.1 | 92/96 |
| *espR4* | 411 | AarF/UbiB kinase family protein | CalR3 | 42/55 | MBB6394636.1 | 89/93 |
| *IS* | 136 | Transposase | -- | -- | GGQ36627.1 | 93/93 |
| *espR5* | 181 | LmbU family transcriptional regulator | -- | -- | MCR3742851.1 | 97/98 |
| *espE3* | 318 | Enediyne biosynthesis protein | CalU15 | 42/54 | WP_185024270.1 | 94/96 |
| *espE4* | 648 | Enediyne biosynthesis protein | CalU14 | 46/57 | WP_258943020.1 | 93/95 |
| *espE5* | 337 | Enediyne biosynthesis protein | CalT3 | 60/71 | WP_185024272.1 | 95/96 |
| *espE* | 1864 | Enediyne polyketide synthase | CalE8 | 45/54 | MBB6394642.1 | 90/92 |
| *espE10* | 156 | Enediyne biosynthesis thioesterase | CalE7 | 50/64 | WP_179834686.1 | 94/96 |
| *espE2* | 228 | F420-dependent oxidoreductase | CalS6 | 37/50 | WP_258943016.1 | 91/93 |
| *espA2* | 407 | Cytochrome P450 | CalE10 | 34/49 | WP_185024276.1 | 97/97 |
| *espA3* | 361 | SAM-dependent methyltransferase | CalO6 | 41/53 | WP_229810767.1 | 93/93 |
| *espA4* | 529 | Aryl-CoA transferase | -- | -- | WP_179834689.1 | 95/97 |
| *espR6* | 498 | TldD/PmbA family protein | CalR6 | 53/64 | MBB6394648.1 | 95/97 |
| *espR7* | 468 | TldD/PmbA family protein | CalR5 | 52/62 | MBB6394649.1 | 92/93 |
| *espS4* | 337 | NDP-sugar dehydratase | CalS3 | 62/72 | WP_185024280.1 | 98/98 |
| *espA5* | 421 | Acyltransferase | -- | -- | WP_244993777.1 | 95/95 |
| *espS5* | 333 | Oxidoreductase | CalS12 | 72/83 | WP_179834693.1 | 96/96 |
| *espS6* | 257 | NDP-sugar *O*-methyltransferase | CalS11 | 63/77 | WP_179834694.1 | 96/98 |
| *espS7* | 245 | SAM-dependent methyltransferase | CalS10 | 62/71 | WP_179834695.1 | 92/95 |
| *espS8* | 476 | NDP-hexose 2,3-dehydratase | CalS14 | 64/73 | GGQ36729.1 | 95/96 |
| *espT2* | 275 | ABC transporter permease | CalT4 | 56/73 | WP_185024285.1 | 96/97 |
| *espT3* | 337 | ABC transporter ATP-binding protein | CalT5 | 58/75 | MBB6394657.1 | 96/96 |
| *espG3* | 413 | Glycosyltransferase | CalG4 | 52/69 | WP_089316031.1 | 91/95 |
| *espS9* | 365 | NAD-dependent epimerase/dehydratase | CalS9 | 61/74 | WP_089316032.1 | 96/97 |
| *espU6* | 218 | Hypothetical protein | CalU13 | 58/68 | WP_185024288.1 | 96/98 |
| *espB* | 223 | DNA alkylation repair protein | -- | -- | NDU71393.1 | 94/96 |
| *espU7* | 233 | Hypothetical protein | CalU12 | 68/77 | WP_179834702.1 | 94/95 |

### Supplementary Table 7. (cont.)

| Gene | AA | Putative function | CAL homologue | % Identity/ Similarity | Protein homologue | % Identity/ Similarity |
|---|---|---|---|---|---|---|
| *espS10* | 407 | Cytochrome P450 | CalE10 | 60/69 | WP_179834703.1 | 96/97 |
| *espG4* | 377 | Glycosyltransferase | CalG3 | 60/75 | WP_179834704.1 | 96/98 |
| *espR8* | 412 | Hypothetical Protein | CalR9 | 64/74 | WP_185024293.1 | 93/96 |
| *espT4* | 590 | ABC transporter substrate-binding protein | CalT6 | 48/61 | WP_179834705.1 | 92/95 |
| *espE6* | 441 | Cysteine desulfurase | CalE9 | 74/87 | MBB6394667.1 | 98/99 |
| *espT5* | 567 | ABC transporter substrate-binding protein | CalT6 | 70/79 | MCR3742820.1 | 97/99 |
| *espA6* | 334 | SAM-dependent methyltransferase | CalO1 | 64/76 | WP_185024296.1 | 96/97 |
| *espS11* | 253 | SAM-dependent methyltransferase | CalS10 | 47/64 | WP_089316042.1 | 92/97 |
| *espG2* | 399 | Glycosyltransferase | CalG2 | 55/65 | GGQ36818.1 | 92/94 |
| *espS12* | 382 | Sugar aminotransferase | CalS13 | 73/83 | WP_185024299.1 | 97/97 |
| *espS13* | 91 | Sulfur carrier protein | CalU18 | 29/47 | WP_179848726.1 | 95/95 |
| *espU8* | 295 | Formylglycine-generating enzyme sulfatase | CalU17 | 58/67 | MBB6394674.1 | 93/97 |
| *espS14* | 450 | NDP-sugar dehydrogenase | CalS8 | 65/73 | NDU71379.1 | 92/96 |
| *espU9* | 265 | Type III CoA transferase | CalU11 | 56/66 | WP_185024302.1 | 95/96 |
| *espE7* | 436 | Cysteine desulfurase | CalE4 | 71/80 | WP_258942986.1 | 93/95 |
| *espE8* | 352 | Methionine γ-lyase | CalE6 | 62/72 | WP_179834715.1 | 96/97 |
| *espE9* | 317 | SAM-dependent methyltransferase | CalE2 | 54/62 | WP_179834716.1 | 90/94 |
| *espS15* | 328 | NDP-sugar dehydratase | CalS3 | 68/77 | WP_179848732.1 | 97/98 |
| *espE11* | 290 | SAM-dependent methyltransferase | CalE5 | 48/61 | WP_179834718.1 | 91/93 |

**Supplementary Table 8.** Comparison of observed esperamicin $A_1$ chemical shifts in $CDCl_3$ ($^1H$ NMR at 600 MHz and $^{13}C$ NMR at 151 MHz) in comparison to previously reported values in MeOD-$d_4$ for $^1H$ shifts (360 MHz) and in $CDCl_3$ for $^{13}C$ shifts (90 MHz).[3] Optical rotation was determined as $[\propto]_D^{23}$ -113 ° $g^{-1}$ mL $dm^{-1}$ (c = 0.37, $CHCl_3$) compared to the literature reported $[\propto]_D^{27}$ -191 ° $g^{-1}$ mL $dm^{-1}$ (c = 0.5, $CHCl_3$).[3,4] nd = not detected.

| Moiety | # | $^1H$ observed [ppm] | $^1H$ literature [ppm] | $^{13}C$ observed [ppm] | $^{13}C$ literature[2] [ppm] | Δ $^1H$ | Δ $^{13}C$ |
|---|---|---|---|---|---|---|---|
| *Core* | **1** | 4.17, 3.78 | 4.11, 3.85 | 40.0 | 39.5 | 0.06, 0.07 | 0.5 |
| | **2** | 6.70 | 6.66 | 131.4 | 130.1 | 0.04 | 1.3 |
| | **3** | | | 133.0 | 135.0 | | 2.0 |
| | **4** | | | 77.8 | 76.8 | | 1.0 |
| | **5** | | | 97.5 | 98.2 | | 0.7 |
| | **6** | | | 83.8 | 83.3 | | 0.5 |
| | **7** | 5.93 | 6.06 | 123.4 | 123.1 | 0.13 | 0.3 |
| | **8** | 5.82 | 5.94 | 125.2 | 124.9 | 0.12 | 0.3 |
| | **9** | | | 87.1 | 88.3 | | 1.2 |
| | **10** | | | 98.3 | 97.5 | | 0.8 |
| | **11** | 6.18 | 6.17 | 70.9 | 68.0 | 0.01 | 2.9 |
| | **12** | | | 130.5 | 131.0 | | 0.5 |
| | **13** | | | 145.9 | 147.0 | | 1.1 |
| | **14** | | | 190.8 | 191.3 | | 0.5 |
| | **15** | 4.28 | 4.31 | 88.3 | 86.0 | 0.03 | 2.3 |
| | **16** | | | 155.2 | 155.0 | | 0.2 |
| | **17** | 3.88 | 3.71 | 56.3 | 52.5 | 0.17 | 3.8 |
| | **1-S-S-S-Me** | 2.56 | 2.54 | 23.1 | 22.6 | 0.02 | 0.5 |
| *4-deoxy-4-methyl thio-α-D-digitoxose (Dig)* | **1** | 4.99 | 4.98 | 99.8 | 99.5 | 0.01 | 0.3 |
| | **2** | 2.16, 1.52 | 1.96, 1.56 | 35.2 | 35.1 | 0.20, 0.04 | 0.1 |
| | **3** | 4.12 | 4.20 | 64.6 | 64.5 | 0.08 | 0.1 |
| | **4** | 2.52 | 2.37 | 55.8 | 55.6 | 0.15 | 0.2 |
| | **5** | 4.06 | 3.87 | 71.5 | 69.2 | 0.19 | 2.3 |
| | **6** | 1.41 | 1.38 | 19.9 | 19.8 | 0.03 | 0.1 |
| | **4-S-Me** | 2.11 | 2.14 | 14.0 | 13.7 | 0.03 | 0.3 |
| *4,6-dideoxy-4-hydroxylamino-α-D-glucose (Glu)* | **1** | 4.57 | 4.57 | 100.4 | 99.5 | 0.00 | 0.9 |
| | **2** | 3.69 | 3.56 | 79.7 | 79.7 | 0.13 | 0.0 |
| | **3** | 3.80 | 3.93 | 69.5 | 69.5 | 0.13 | 0.0 |
| | **4** | 2.29 | 2.26 | 68.2 | 68.1 | 0.03 | 0.1 |
| | **5** | 3.69 | 3.62 | 71.5 | 71.7 | 0.07 | 0.2 |
| | **6** | 1.31 | 1.30 | 17.8 | 17.5 | 0.01 | 0.3 |
| *Amino-pentose (Pen)* | **1** | 5.36 | 5.54 | 98.8 | 97.2 | 0.18 | 1.6 |
| | **2** | 2.34, 1.63 | 2.47, 1.60 | 33.5 | 34.0 | 0.13, 0.03 | 0.5 |
| | **3** | nd | 3.74 | 73.1 | 75.8 | | 2.7 |
| | **4** | 3.15 | 3.15 | 58.8 | 57.1 | 0.00 | 1.7 |
| | **5** | 4.17, 3.90 | 3.84, 3.70 | 58.9 | 62.3 | 0.33, 0.20 | 3.4 |
| | **6** | 3.48 | 3.30 | 49.8 | 47.2 | 0.18 | 2.6 |
| | **7** | 1.46 | 1.23 | 18.5 | 22.2 | 0.23 | 3.7 |
| | **8** | 1.37 | 1.21 | 20.2 | 23.4 | 0.16 | 3.2 |
| | **3-O-Me** | 3.43 | 3.42 | 56.6 | 56.0 | 0.01 | 0.6 |
| *2-Desoxy-Fucose (Fuc)* | **1** | 5.36 | 5.48 | 99.7 | 99.0 | 0.12 | 0.7 |
| | **2** | 2.38, 2.30 | 2.32, 2.15 | 29.2 | 29.0 | 0.06, 0.15 | 0.2 |
| | **3** | 5.51 | 5.48 | 70.0 | 70.2 | 0.03 | 0.2 |
| | **4** | 4.03 | 3.90 | 68.9 | 66.7 | 0.13 | 2.2 |
| | **5** | 4.57 | 4.62 | 67.6 | 68.8 | 0.05 | 1.2 |

**Supplementary Table 7. (cont.)**

| Moiety | # | $^{1}H$ observed [ppm] | $^{1}H$ literature [ppm] | $^{13}C$ observed [ppm] | $^{13}C$ literature[2] [ppm] | Δ $^{1}H$ | Δ $^{13}C$ |
|---|---|---|---|---|---|---|---|
| | **6** | 1.39 | 1.25 | 16.8 | 16.5 | 0.14 | 0.3 |
| *Amino-veratric acid (Ver)* | **1** | | | 166.7 | 166.4 | | 0.3 |
| | **2** | | | 107.6 | 107.6 | | 0.0 |
| | **3** | 7.49 | 7.63 | 112.7 | 112.5 | 0.14 | 0.2 |
| | **4** | | | 144.3 | 144.0 | | 0.3 |
| | **5** | | | 154.3 | 153.8 | | 0.5 |
| | **6** | 8.60 | 8.44 | 104.1 | 103.7 | 0.16 | 0.4 |
| | **7** | | | 137.2 | 136.7 | | 0.5 |
| | **4-O-Me** | 3.88 | 3.91 | 56.3 | 56.0 | 0.03 | 0.3 |
| | **5-O-Me** | 3.96 | 3.85 | 56.3 | 56.0 | 0.11 | 0.3 |
| *2-Methoxy-acrylic acid (Acr)* | **1** | | | 161.1 | 160.7 | | 0.4 |
| | **2** | | | 154.7 | 154.4 | | 0.3 |
| | **3** | 5.46, 4.54 | 5.38, 4.67 | 91.0 | 90.5 | 0.08, 0.13 | 0.5 |
| | **2-O-Me** | 3.81 | 3.79 | 56.3 | 56.0 | 0.03 | 0.3 |

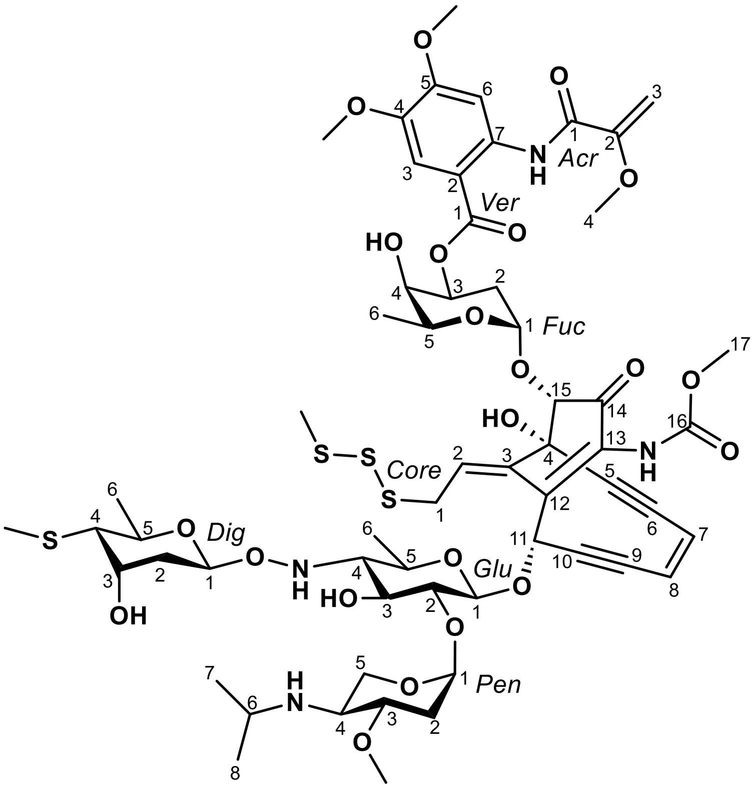

**Supplementary Table 9.** Strains and plasmids used in this study.

| Name | Description | Reference or source |
|---|---|---|
| **Strains** | | |
| Turbo | *E. coli* strain for general cloning | Life Technologies |
| BL21(DE3) | *E. coli* host for protein overproduction | Life Technologies |
| SB24001 | BL21(DE3) cells containing pBS24001 for protein production | This study |
| *Kocuria rhizophila* ATCC 9341 | Strain for testing antibacterial activity | ([5]) |
| *Micrococcus luteus* NPDC049463 | Strain used for cloning *dnaN* from | This study |
| *Mycobacterium smegmatis* ATCC 607 | Strain for testing antibacterial activity | ([6]) |
| *Streptomyces albidoflavus* J1074 | Strain for testing antibacterial activity | ([7]) |
| *Mycobacterium tuberculosis* R37a ATCC 25177 | Strain for testing antibacterial activity | ([8]) |
| *Mycobacterium intracellulare* DSM 43223 | Strain for testing antibacterial activity | ([9]) |
| *Mycobacterium abscessus* ATCC 19977 | Strain for testing antibacterial activity | ([10]) |
| **Plasmids** | | |
| pBS3080 | pRSFDuet-1 derived plasmid containing a *Bsm*FI site for ligation-independent cloning (LIC) and encodes a TEV protease site after the *N*-terminal $His_6$-tag | ([11]) |
| pBS24001 | $dnaN_{M.luteus}$ cloned into pBS3080 | This study |

**Supplementary Table 10.** Genus abbreviations for Extended Data and SI figures.

| Genus | Abbreviation |
|---|---|
| *Actinoallomurus* | AAM |
| *Actinocorallia* | ACO |
| *Actinocrispum* | ACR |
| *Actinokineospora* | AKS |
| *Actinomadura* (B) | ACM |
| *Actinophytocola* | APY |
| *Actinoplanes* | ACP |
| *Actinopolymorpha* | APM |
| *Actinosynnema* | ASY |
| *Aeromicrobium* | AEM |
| *Agrococcus* | AGC |
| *Agromyces* | AGR |
| *Amycolatopsis* | AMY |
| *Arthrobacter* (B,D,E,F,G,I,K) | ART |
| *Asanoa* | ASN |
| *Brachybacterium* | BRB |
| *Brevibacterium* | BRE |
| *Catellatospora* | CAS |
| *Catenuloplanes* | CAP |
| *Cellulomonas* | CEL |
| *Cellulosimicrobium* | CLM |
| *Citricoccus* | CIT |
| *Corynebacterium* | COR |
| *Cryptosporangium* | CRP |
| *Curtobacterium* | CRB |
| *Dactylosporangium* | DCT |
| *Dietzia* | DTZ |
| *Embleya* | EMB |
| *Enteractinococcus* | EAC |
| *Frigoribacterium* | FRG |
| *Glutamicibacter* | GMB |
| *Glycomyces* | GLY |
| *Gordonia* | GOR |
| *Hamadaea* | HAM |
| *Isoptericola* | ITC |
| *Janibacter* | JAN |
| *Kibdelosporangium* | KIB |
| *Kitasatospora* | KTS |
| *Kocuria* | KOC |
| *Kribbella* | KRB |
| *Kutzneria* | KTZ |

**Supplementary Table 10. (cont.)**

| Genus | Abbreviation |
|---|---|
| *Leifsonia* | LFS |
| *Lentzea* | LTZ |
| *Leucobacter* | LEU |
| *Longispora* | LGS |
| *Microbacterium* | MCB |
| *Microbispora* | MBP |
| *Micrococcus* | MCC |
| *Micromonospora* (G) | MCM |
| *Microtetraspora* | MTS |
| *Modestobacter* | MOD |
| *Mycobacterium* | MYC |
| *Nesterenkonia* | NEK |
| *Nocardia* | NCD |
| *Nocardioides* | NCS |
| *Nocardiopsis* | NCP |
| *Nonomuraea* | NMU |
| *Oerskovia* | OER |
| *Paenarthrobacter* | PAE |
| *Patulibacter* | PAT |
| *Polymorphospora* | PMS |
| *Promicromonospora* | PMC |
| *Pseudarthrobacter* | PAC |
| *Pseudonocardia* | PSN |
| *Rhodococcus* (B,C) | RHD |
| *Rhodoglobus* | RHG |
| *Rothia* | RTH |
| *Saccharomonospora* | SMS |
| *Saccharopolyspora* (C) | SPS |
| *SCUT-3* | SCU |
| *Specibacter* | SPB |
| *Sphaerisporangium* | SSG |
| *Spirillospora* | SPL |
| *Spongisporangium* | SSP |
| *Streptodolium* | SDM |
| *Streptomyces* (D) | STR |
| *Streptosporangium* | STS |
| *Terrabacter* | TER |
| *Tsukamurella* | TSU |
| *Umezawaea* | UMZ |
| *Uniformispora* | UFS |

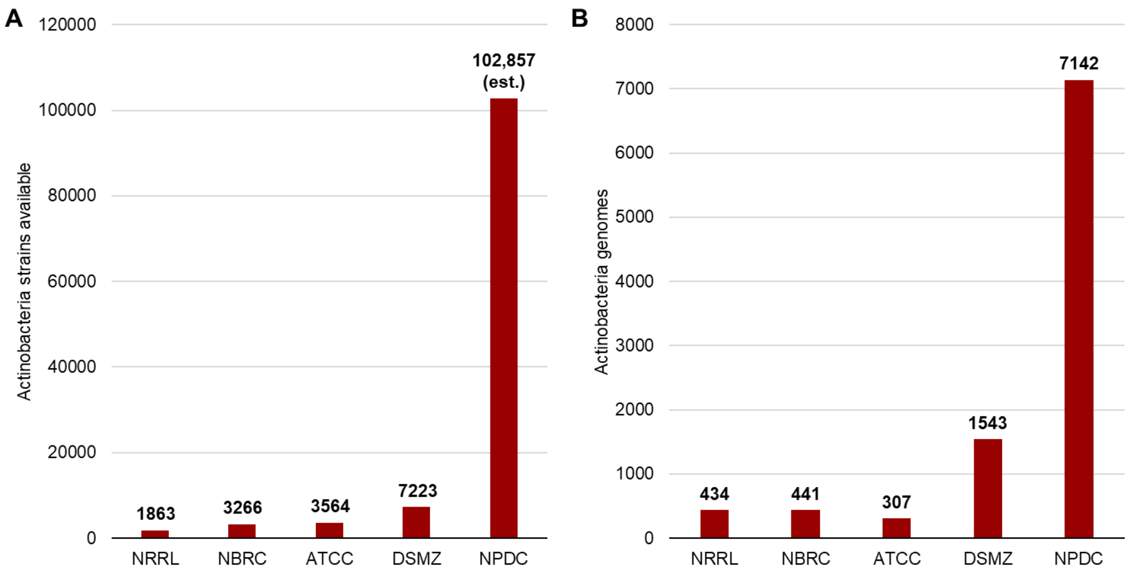

**Supplementary Fig. 1.** Comparison of the NPDC collection with other Actinobacteria strain collections. (A) The number of Actinobacteria strains listed on websites of major worldwide strain collections range from less than 2000 strains to just over 7000 strains, in comparison to more than 102,000 predicted Actinobacteria in the NPDC collection based on current sequencing ratios (84.0% Actinobacteria) and a total NPDC collection size of 122,449 strains. (B) Using data in the NCBI genome database and any genomic information stored on collection websites, published genomes were totaled. Any overlaps between collections were counted for each collection, so the total number of sequenced strains between collections is lower than the sum. All data collected May 2023. [12-15] NRRL = Agricultural Research Service Culture Collection; NBRC = NITE Biological Resource Center; ATCC = American Type Culture Collection; DSMZ = German Collection of Microorganisms and Cell Cultures; NPDC = Natural Products Discovery Center.

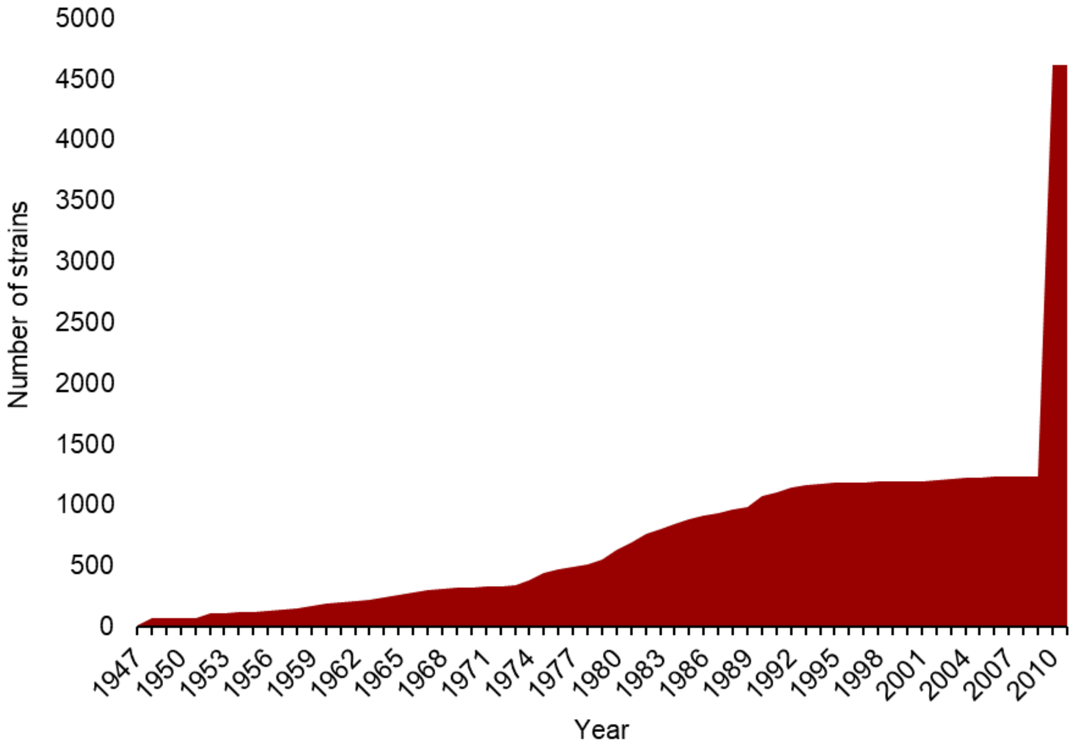

**Supplementary Fig. 2.** Sample isolation years for sequenced NPDC strains with existing metadata. 34.6% of NPDC strains have associated isolation dates. The spike in strains isolated during the 2010s is exaggerated in this subset of sequenced strains and is not representative of the entire collection. [16]

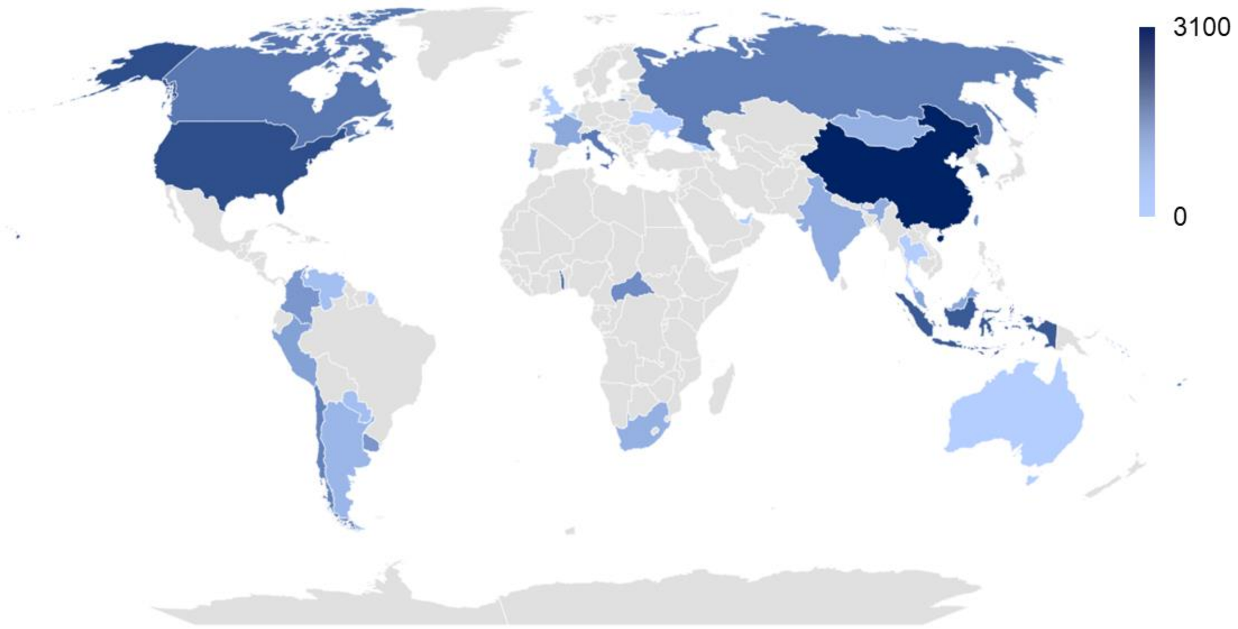

**Supplementary Fig. 3.** Sample collection locations for sequenced NPDC strains with existing metadata. Color intensity correlates with the number of strains. 6.7% of NPDC strains have associated isolation locations. See Supplementary Table 1 for exact values. The spike in strains isolated from China is exaggerated in this subset of sequenced strains and is not representative of the entire collection. [16]

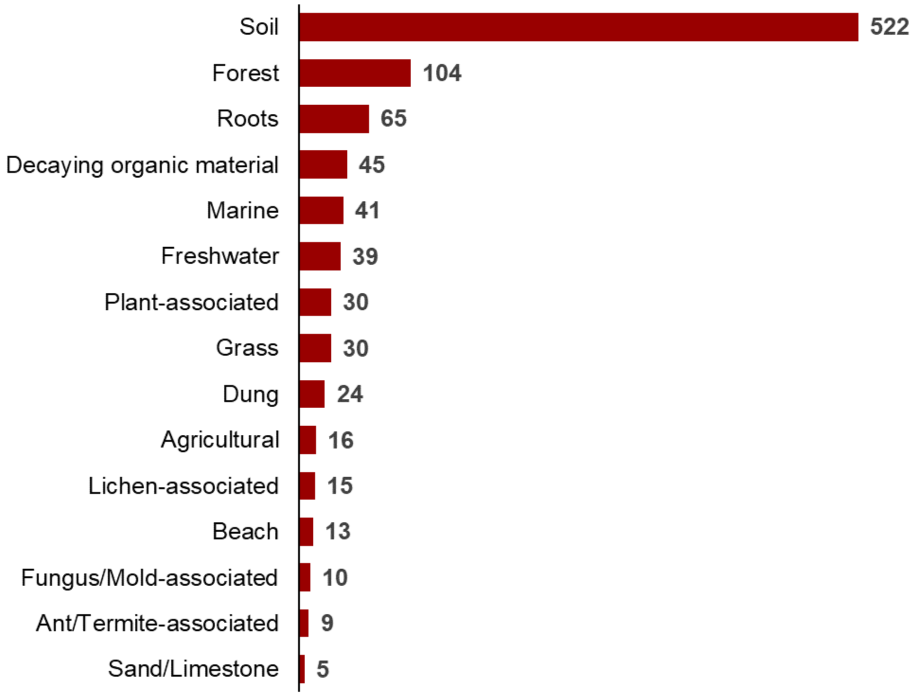

**Supplementary Fig. 4.** Sample collection general environmental categories for sequenced NPDC strains with existing metadata. 7.1% of NPDC strains have associated isolation environments.

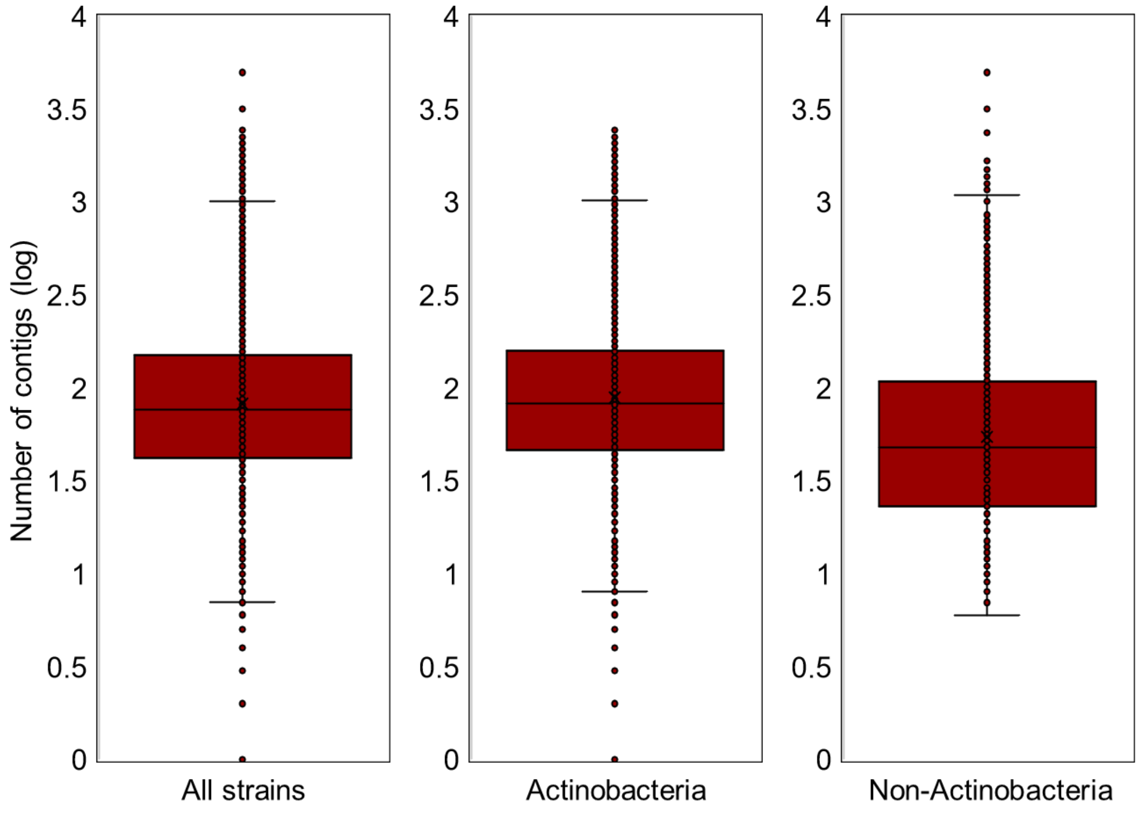

**Supplementary Fig. 5.** Number of contigs for the initial 8490 assembled NPDC genomes. Distributions of contig numbers were plotted for all strains, all Actinobacteria, and all non-Actinobacteria. Lower values indicate higher quality genomes. Non-Actinobacteria genomes are typically smaller than Actinobacteria genomes (Supplementary Fig. 7), leading to improved assembly criteria.

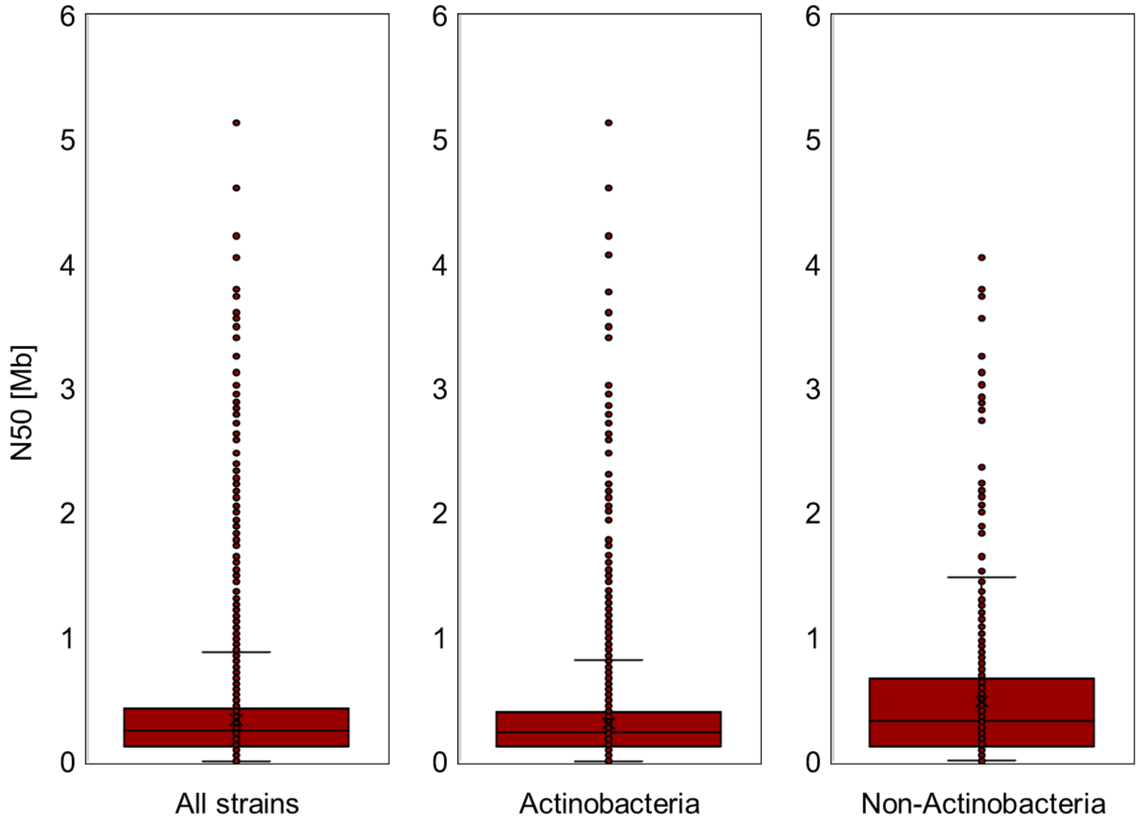

**Supplementary Fig. 6.** N50 values for the initial 8490 assembled NPDC genomes. Distributions of N50 values were plotted for all strains, all Actinobacteria, and all non-Actinobacteria. Non-Actinobacteria genomes are typically smaller than Actinobacteria genomes (Supplementary Fig. 7), leading to improved assembly criteria.

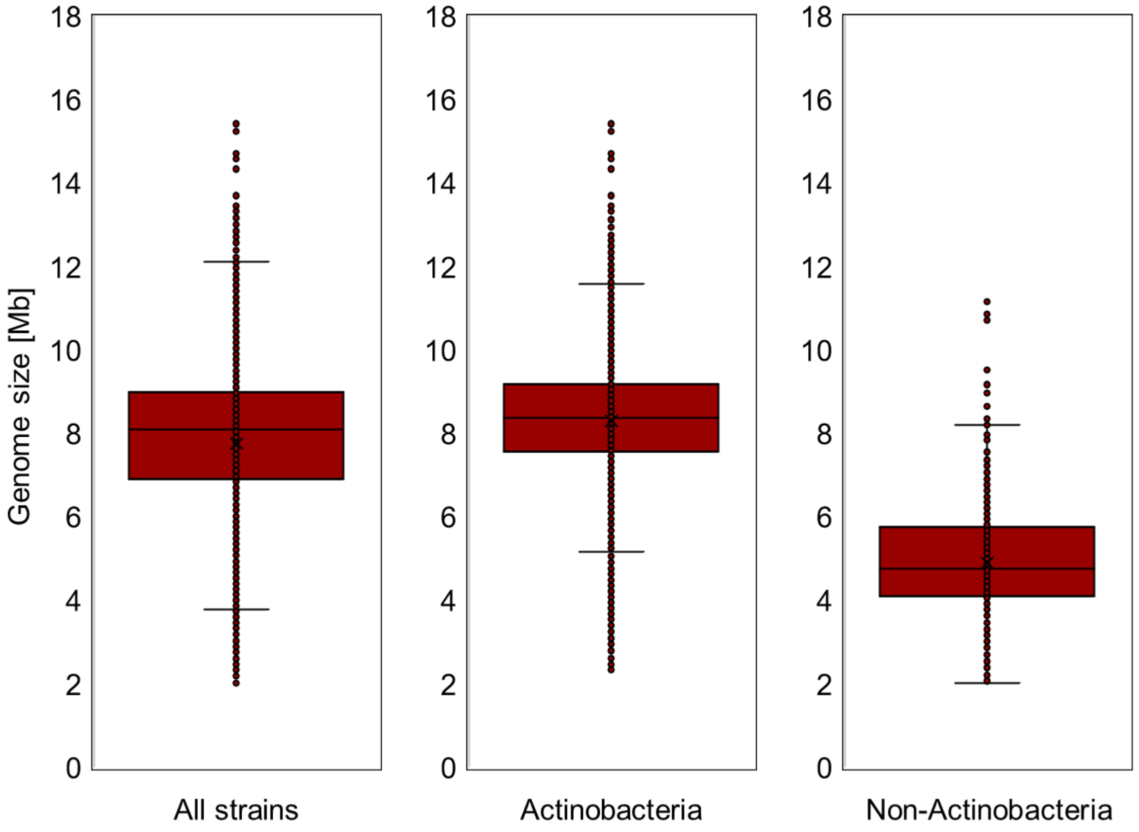

**Supplementary Fig. 7.** Genome sizes for the initial 8490 assembled NPDC genomes. Distributions of genome sizes were plotted for all strains, all Actinobacteria, and all non-Actinobacteria.

A

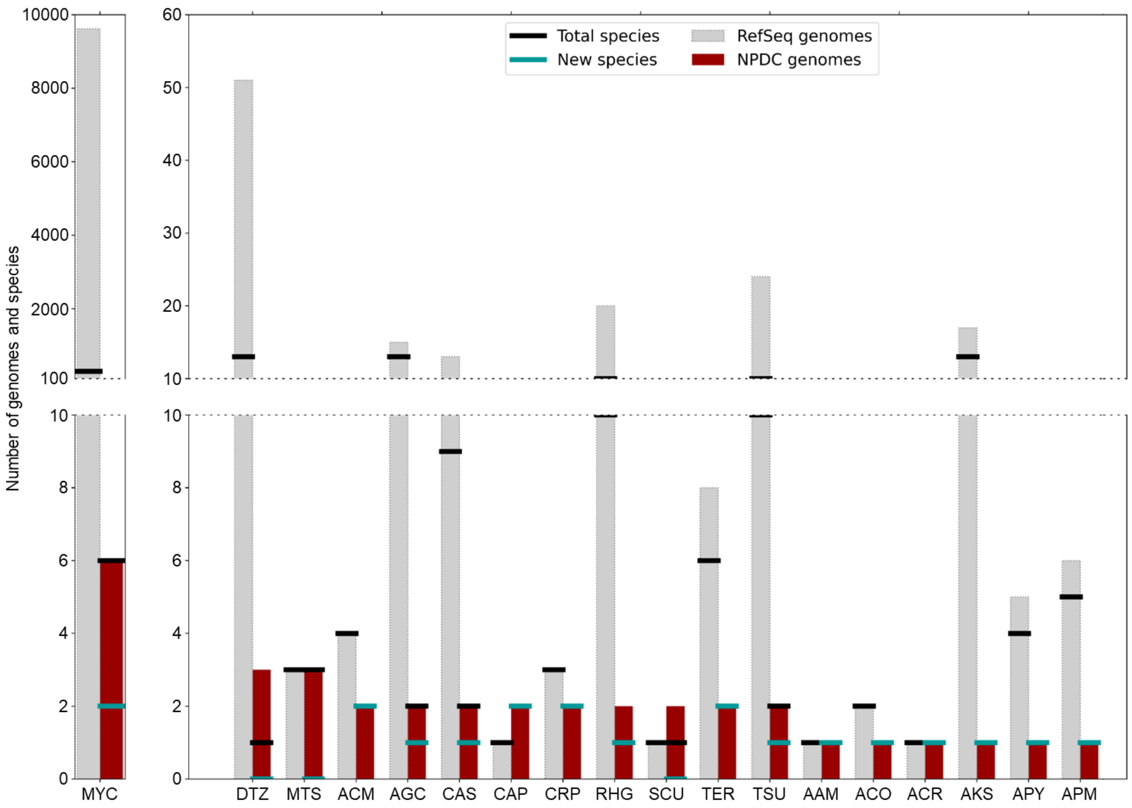
Number of genomes and species
Total species
New species
RefSeq genomes
NPDC genomes
MYC
DTZ
MTS
ACM
AGC
CAS
CAP
CRP
RHG
SCU
TER
TSU
AAM
ACO
ACR
AKS
APY
APM

B

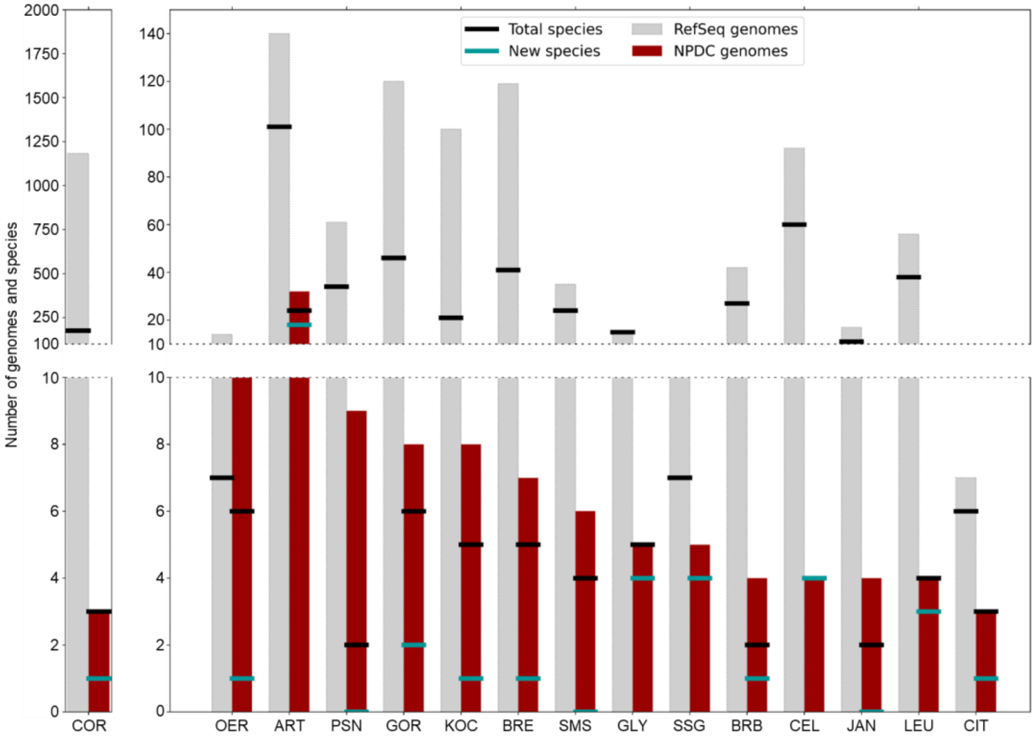

Total species
New species
RefSeq genomes
NPDC genomes
Number of genomes and species
2000
1750
1500
1250
1000
750
500
250
100
10
8
6
4
2
0
140
120
100
80
60
40
20
10
COR
OER
ART
PSN
GOR
KOC
BRE
SMS
GLY
SSG
BRB
CEL
JAN
LEU
CIT

C

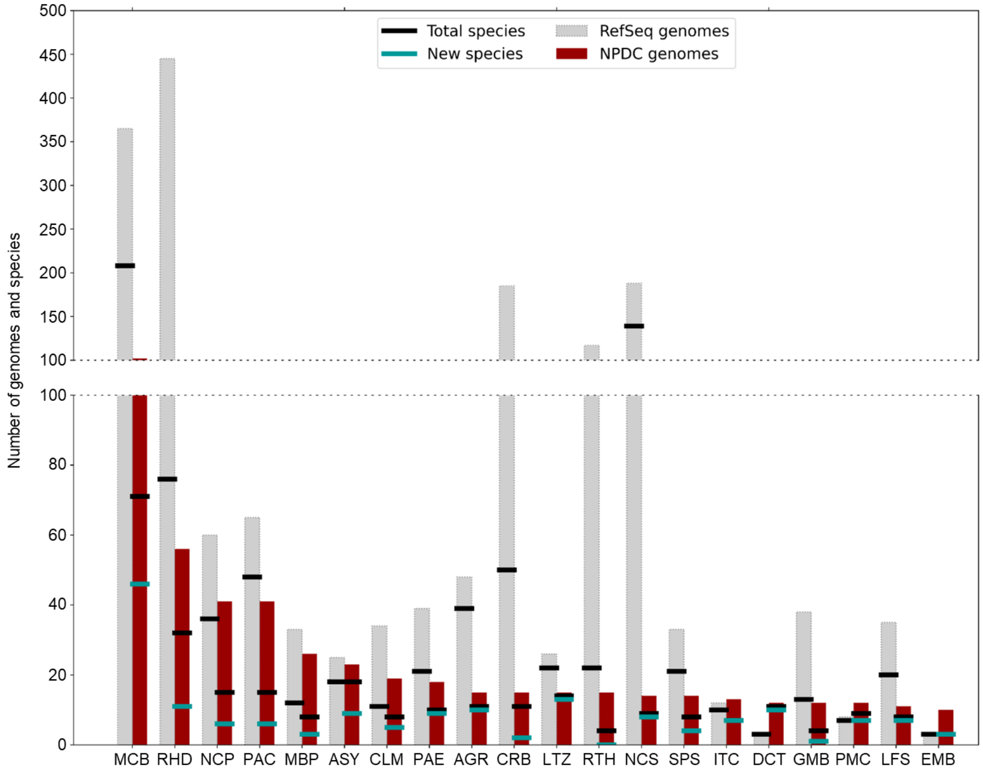

Total species
New species
RefSeq genomes
NPDC genomes
Number of genomes and species
500
450
400
350
300
250
200
150
100
80
60
40
20
0
MCB
RHD
NCP
PAC
MBP
ASY
CLM
PAE
AGR
CRB
LTZ
RTH
NCS
SPS
ITC
DCT
GMB
PMC
LFS
EMB

D

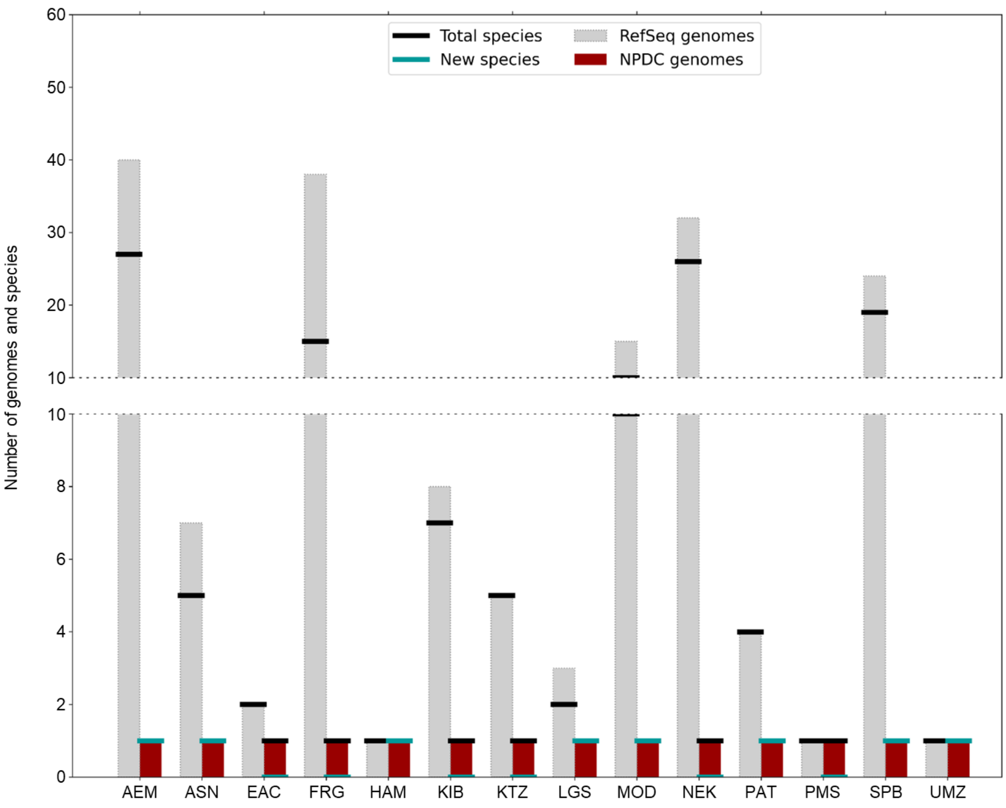

**Supplementary Fig. 8.** NPDC and RefSeq genomes and species by Actinobacteria genus. Actinobacteria genomes from genera in the NPDC (red bars) are compared to genomes from RefSeq (grey bars). Total number of species per genus based on a 0.05 Mash distance cut-off are indicated by black lines. Numbers of new species in NPDC lacking a closely related representative in GTDB are indicated by teal lines. Also see Fig. 1D. See Supplementary Table 10 for abbreviations. Genera are grouped based on the total number of genomes.

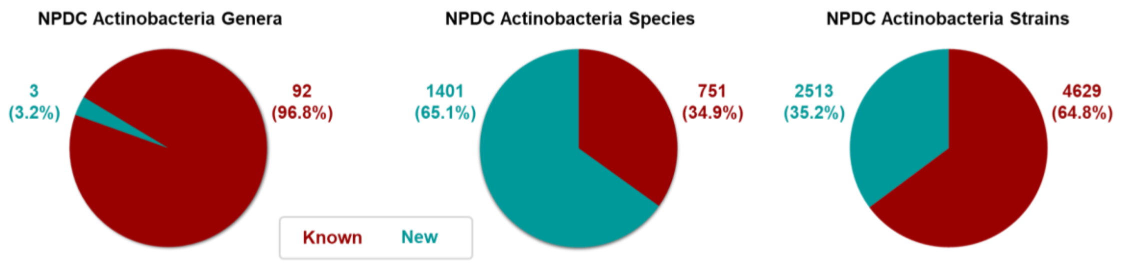

**Supplementary Fig. 9.** NPDC Actinobacteria novelty at different taxonomic levels. Known indicates genera, species, or strains belonging to species that have non-NPDC genomes in GTDB. 'New' indicates these genera, species, or strains could not be assigned to a previously assigned genus or species using GTDB.

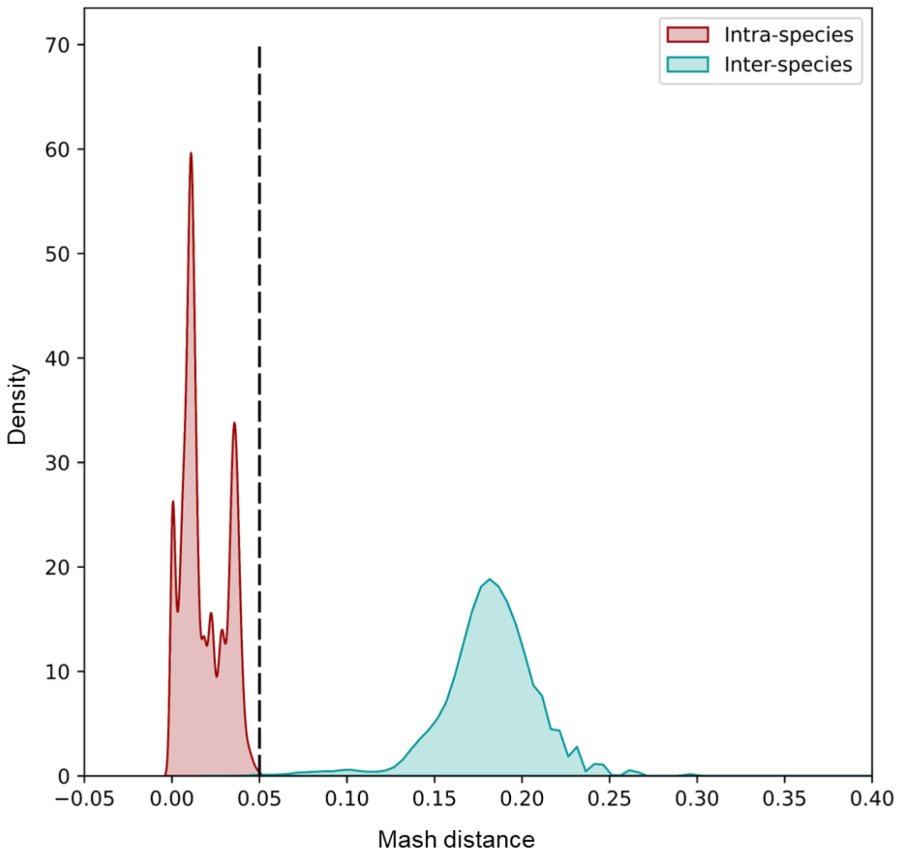

**Supplementary Fig. 10.** Mash distance distribution among Actinobacteria within the same species and across species. Mash distances between 0 and 0.05 were used to delineate species cut-offs for this study.

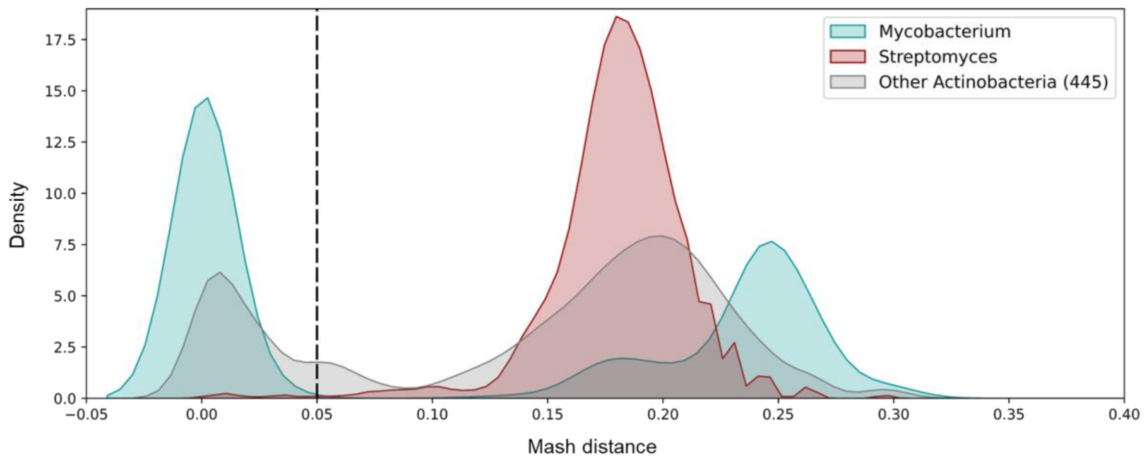

**Supplementary Fig. 11.** Mash distance distribution within Actinobacteria. Mash distances less than 0.05 indicate same strains within the same species. Most *Mycobacterium* belong to a single species, in contrast to most *Streptomyces*, which display far greater genomic diversity.

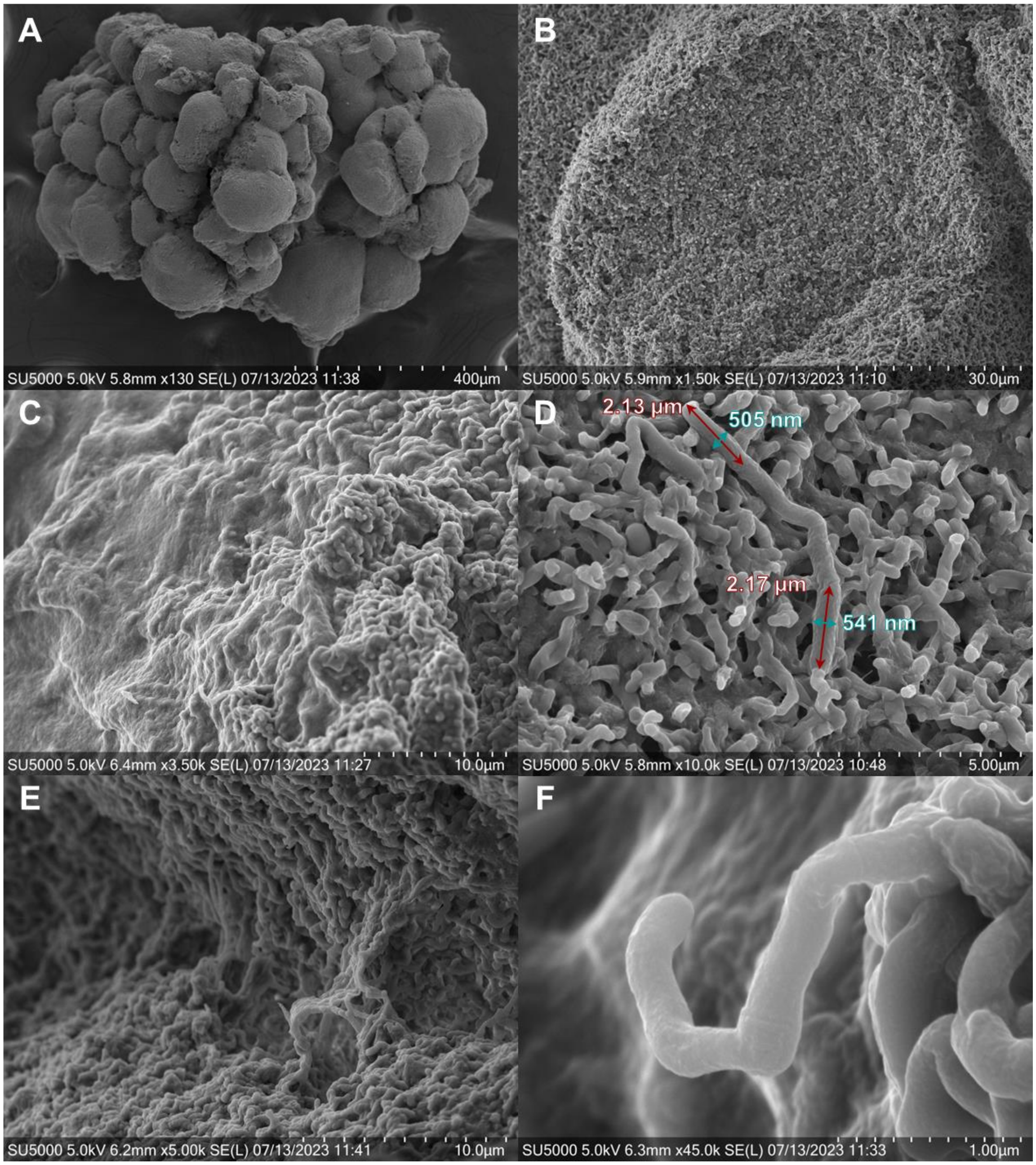

**Supplementary Fig. 12.** SEM images of *Spongisporangium articulatum* NPDC049639. (A) The strain presents characteristic spherical conglomerates, which were used as a basis for the genus name ("*sporangium"* = deriving from the Greek σπορά *(sporá)* "seed" and ἀγγεῖον *(angeîon)* "vessel"). (B) Fractured edge of a colony surface highlighting the uniform, spongelike structure inside and on the surface of the conglomerates. (C) Biofilm covering hyphae and spores. (D) Segmented aerial hyphae on colony surface. The longest segments observed display lengths of up to 2.17 µm and widths of 505 to 541 nm. (E) Kinked hyphae forming arched structures connecting the surfaces of two spherical conglomerates. (F) Aerial hypha with alternating small segments and large segments with up to 90° angled junctions, which were used as a basis for the species name ("*articulatum*" = Latin for "joint"). Also see Fig. 3B.

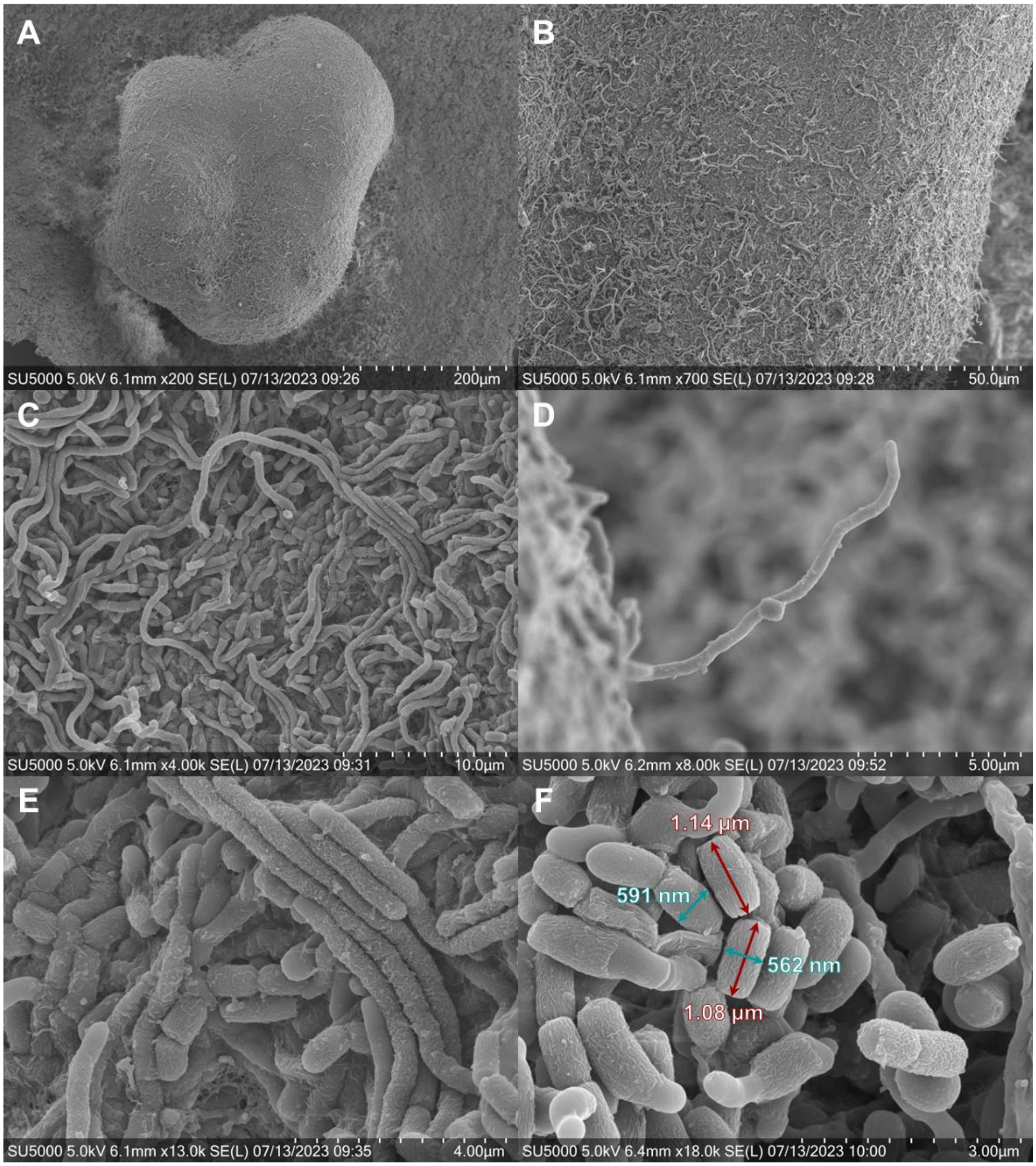

**Supplementary Fig. 13.** SEM images of *Streptodolium elevatio* NPDC002781. (A) The strain, isolated from a forest soil sample in Qinhai, China, presents characteristic conglomerates on an otherwise flat bacterial lawn, serving as inspiration for the species name (“*elevatio”* = Latin for “raising” or “elevation”). (B) Colony surface with long protruding aerial hyphae. (C) Typical aerial hyphae and spores on the colony surface similar to those found in *Streptomyces*, which inspired the first part of the genus name. (D) Protruding aerial hyphae on the conglomerate presented in (A). (E) Zoomed view of (C) highlighting the fibrous sheath coating aerial hyphae and spores. (F) Barrel-shaped spores inspiring the second half of the genus name (“*dolium”* = Latin for “large jar” or “vessel”). Spores display widths between 560 and 590 nm with lengths of 1.06 to 1.16 µm. Also see Fig. 3D.

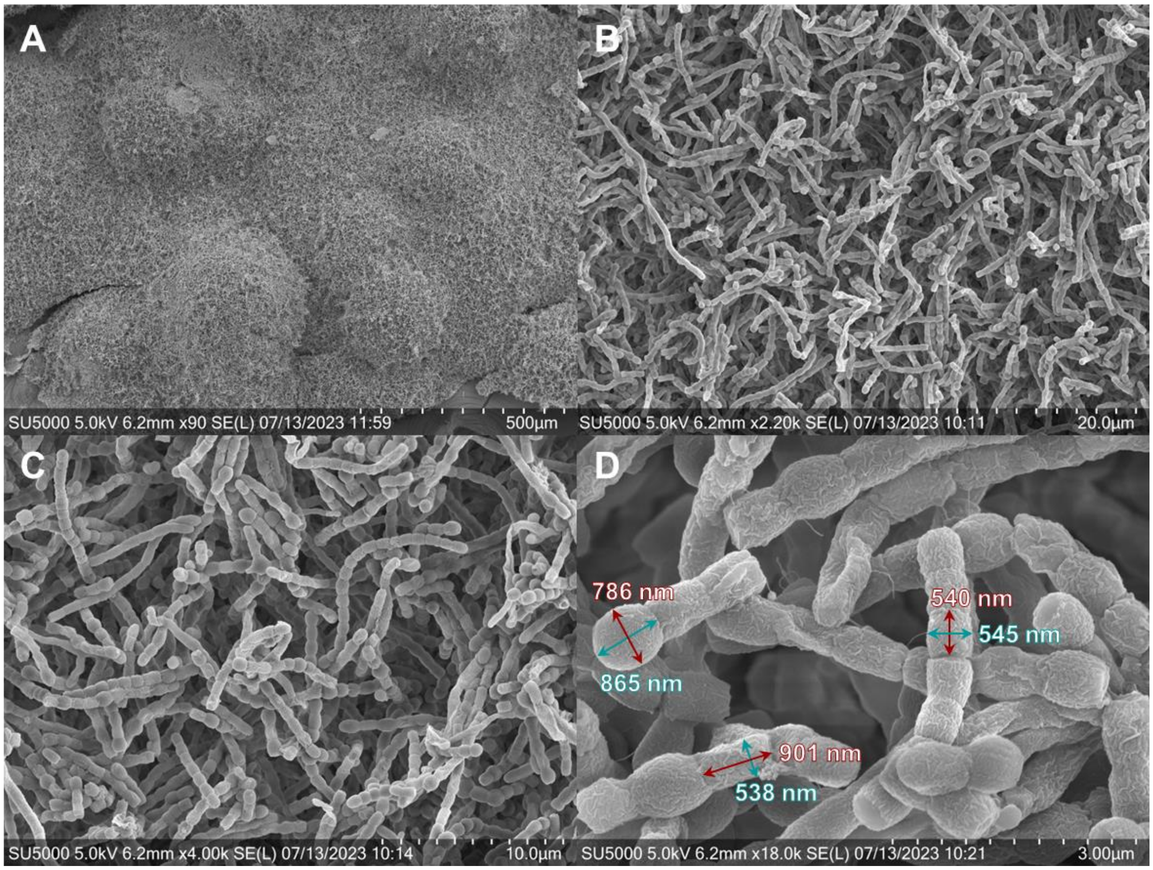

**Supplementary Fig. 14.** SEM images of *Uniformispora flossi* NPDC059280. (A) The strain grows as a relatively flat bacterial lawn with slight mounds on the surface. (B) and (C) Uniform spores are presented as long chains on the colony surface, which were used as inspiration for the genus name. (D) Zoomed view of spore chains containing nearly spherical terminal spores (786 x 865 nm) with elongated adjacent spores (538 x 901 nm) and otherwise almost cubical spores displaying widths comparable to the elongated spores of 545 to 561 nm but lengths between 481 and 535 nm. The species name was assigned in honor of Professor Heinz Floss and his contributions to the field of natural products and natural product biosynthesis. Also see Fig. 3E.

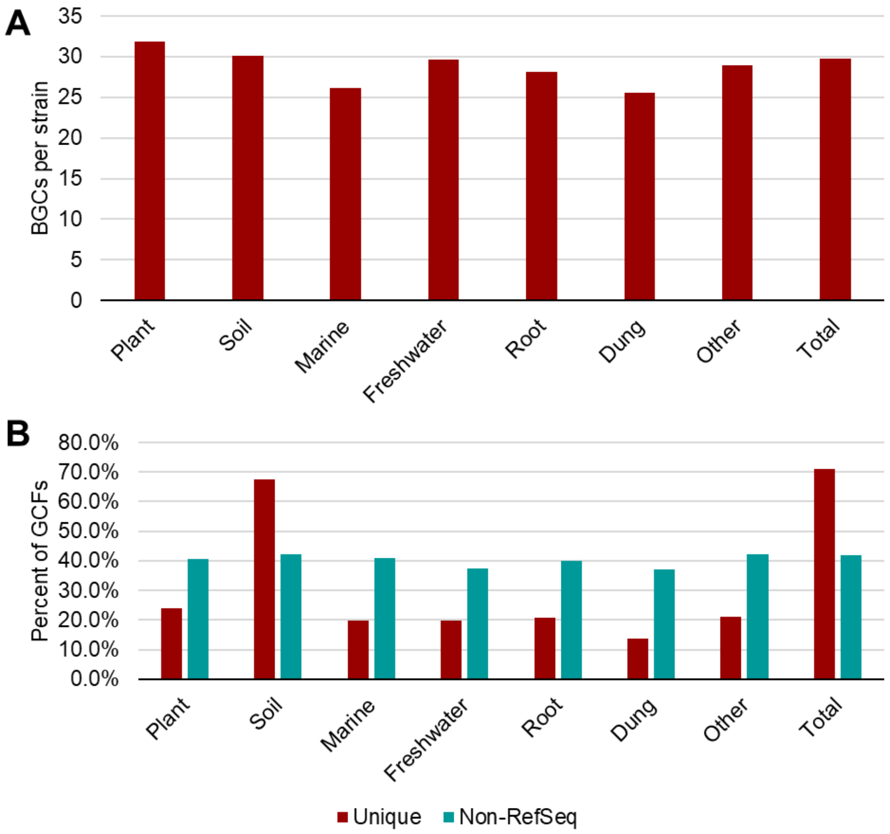

**Supplementary Fig. 15.** Analysis of biosynthetic potential of 957 NPDC Actinobacteria by environment. The 957 sequenced NPDC Actinobacteria with environmental metadata were categorized into seven major categories (65 'Plant', 651 'Soil', 32 'Marine', 28 'Freshwater', 63 'Root', 24 'Dung', and 94 'Other'). (A) The average number of BGCs was calculated for each category as well as for all 957 strains ('Total'). While plant-associated Actinobacteria had the highest number of BGCs, marine- and dung-associated Actinobacteria had the lowest. (B) For each environment, the percent of GCFs that were only identified in a single environment ('Unique') varies especially between 'Soil' (66.7%) and other environments, e.g., 'Dung' (13.8%). This trend does not correlate with the frequency of non-RefSeq GCFs, which are found in an environment-independent manner. Note: many strains have more detailed environmental metadata than presented here, but for simplicity, related environments have been grouped together into these general categories.

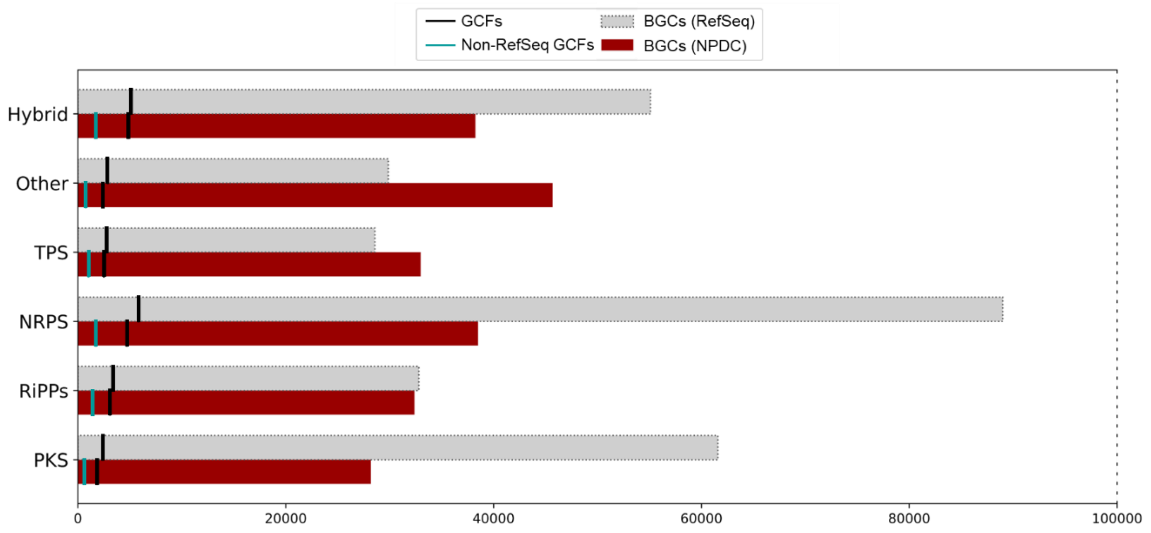

**Supplementary Fig. 16.** BGC and GCF biosynthetic classes in RefSeq and NPDC Actinobacteria. PKS = polyketide synthase; RiPPs = ribosomally synthesized and post-translationally modified peptides; NRPS = non-ribosomal peptide synthetase; TPS = terpene synthase; Hybrid = PKS + NRPS.

A

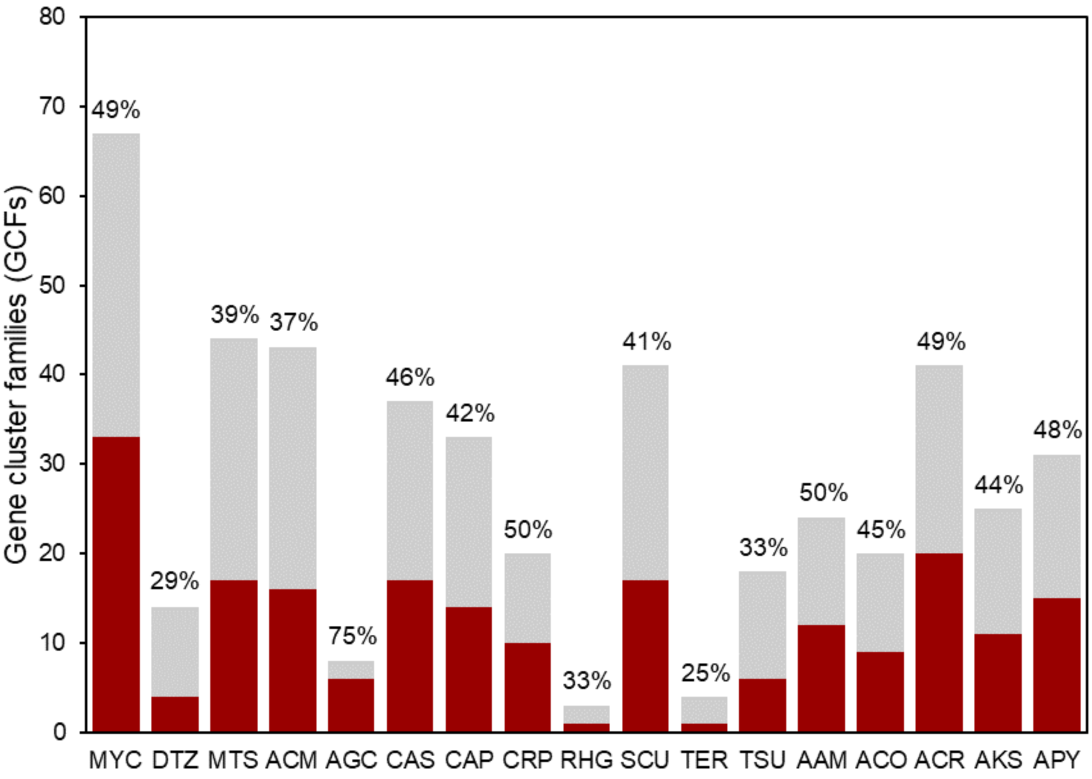

Gene cluster families (GCFs)
80
70
60
50
40
30
20
10
0
49%
29%
39%
37%
75%
46%
42%
50%
33%
41%
25%
33%
50%
45%
49%
44%
48%
MYC DTZ MTS ACM AGC CAS CAP CRP RHG SCU TER TSU AAM ACO ACR AKS APY

B

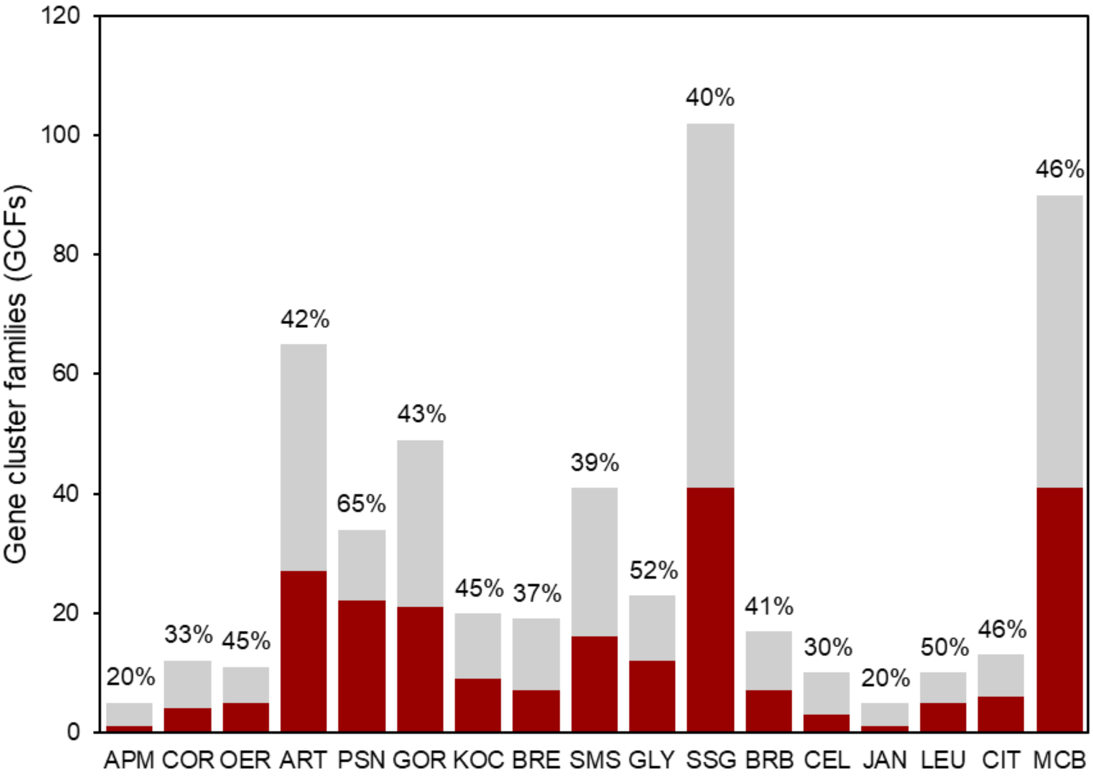

Gene cluster families (GCFs)
120
100
80
60
40
20
0
20%
33%
45%
42%
65%
43%
45%
37%
39%
52%
40%
41%
30%
20%
50%
46%
46%
APM COR OER ART PSN GOR KOC BRE SMS GLY SSG BRB CEL JAN LEU CIT MCB

C

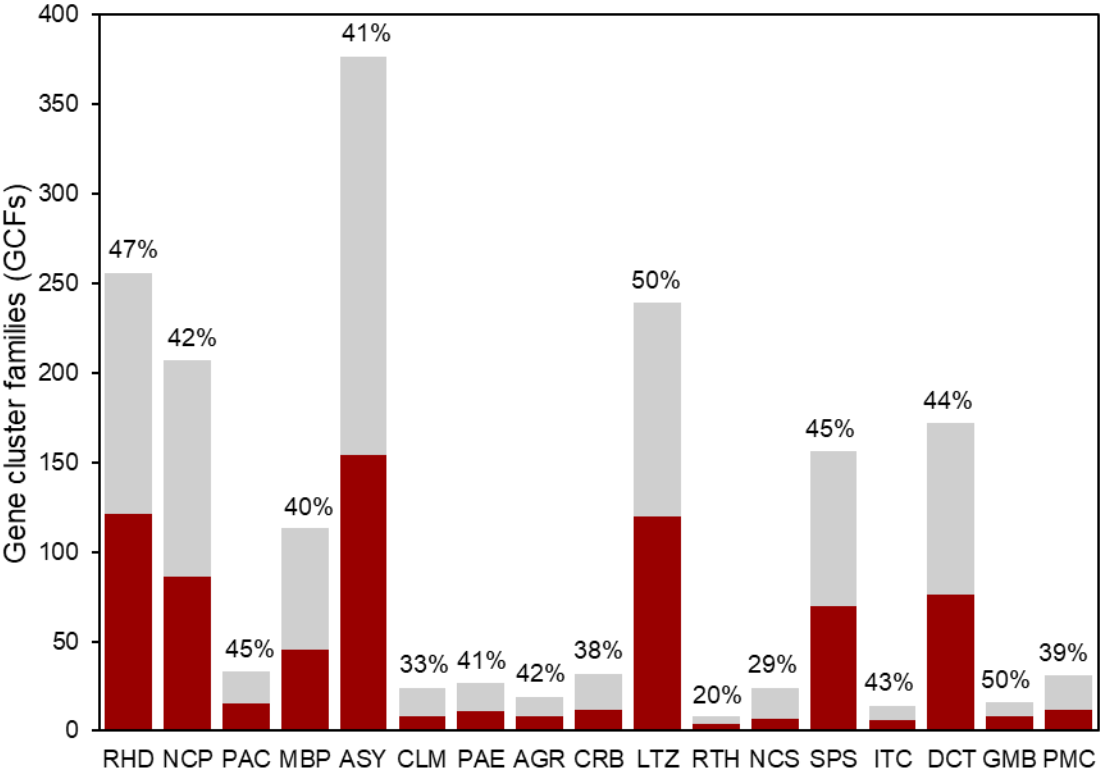

Gene cluster families (GCFs)
400
350
300
250
200
150
100
50
0
47%
42%
45%
40%
41%
33%
41%
42%
38%
50%
20%
29%
45%
43%
44%
50%
39%
RHD NCP PAC MBP ASY CLM PAE AGR CRB LTZ RTH NCS SPS ITC DCT GMB PMC

D

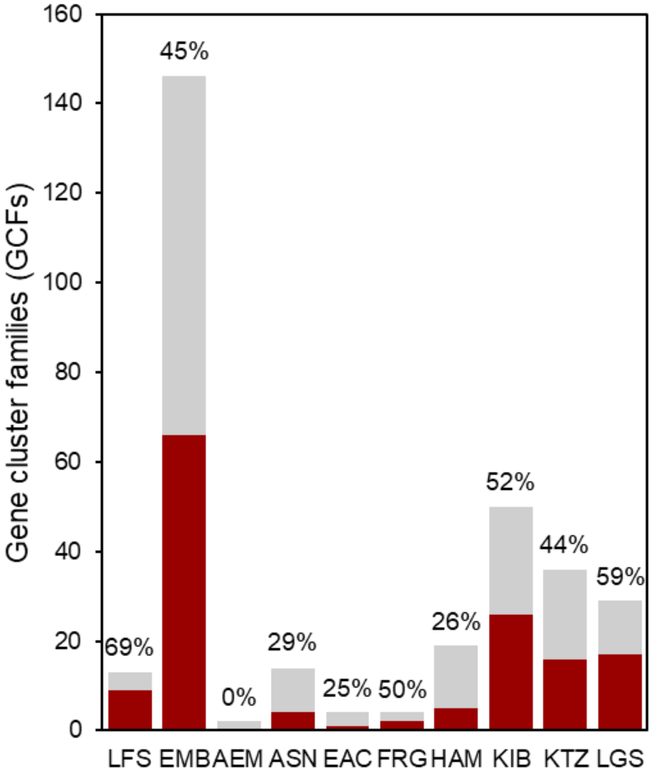

Gene cluster families (GCFs)
160
140
120
100
80
60
40
20
0
69%
45%
0%
29%
25%
50%
26%
52%
44%
59%
LFS EMB AEM ASN EAC FRG HAM KIB KTZ LGS

**E**

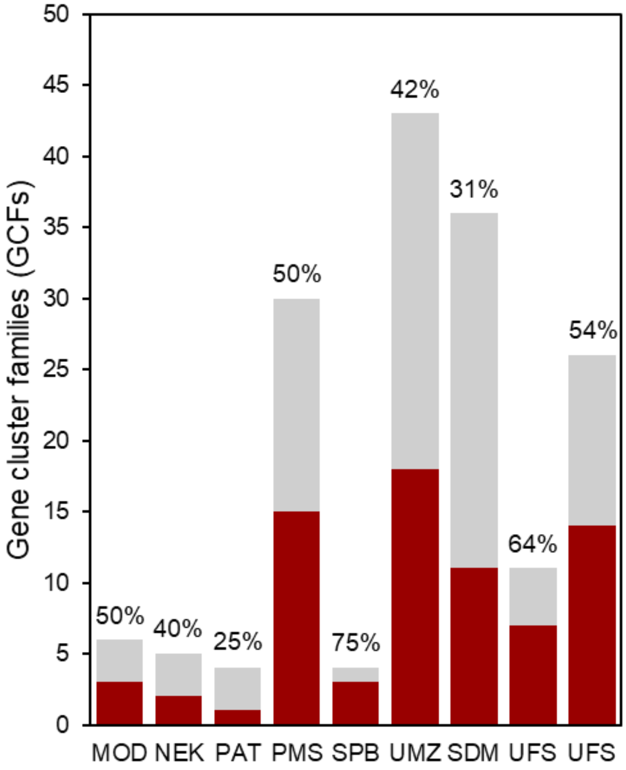

**Supplementary Fig. 17.** Distribution of non-RefSeq NPDC GCFs (red) and GCFs observed in both NPDC and RefSeq genomes (grey) across NPDC genera. Also see Fig. 4B. Genera are grouped based on the total number of GCFs. See Supplementary Table 10 for abbreviations.

A

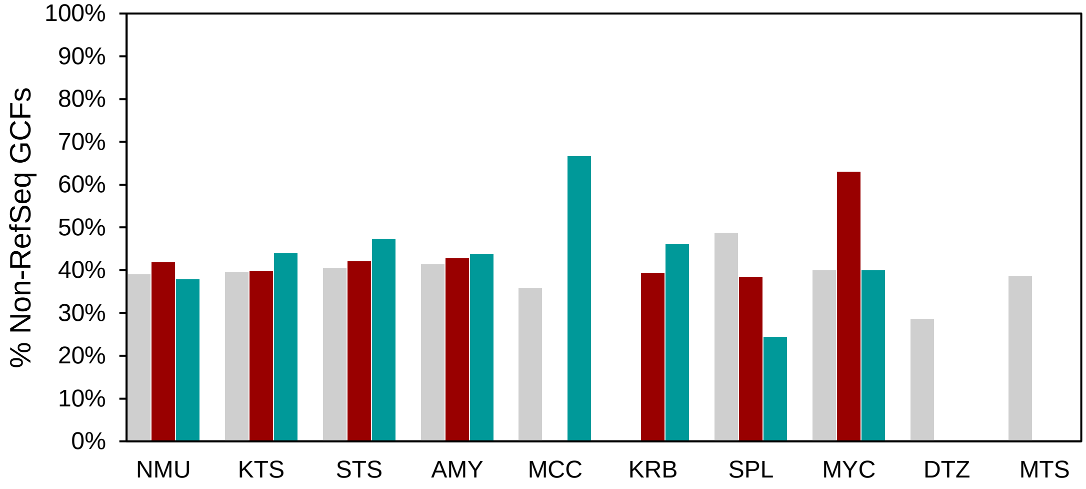

100%
90%
80%
70%
60%
50%
40%
30%
20%
10%
0%
% Non-RefSeq GCFs
NMU
KTS
STS
AMY
MCC
KRB
SPL
MYC
DTZ
MTS

B

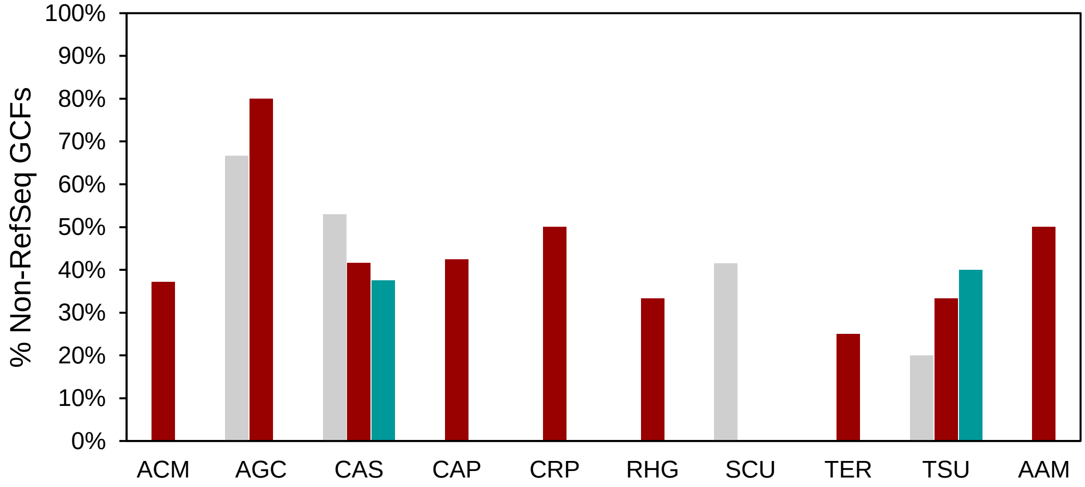

100%
90%
80%
70%
60%
50%
40%
30%
20%
10%
0%
% Non-RefSeq GCFs
ACM
AGC
CAS
CAP
CRP
RHG
SCU
TER
TSU
AAM

C

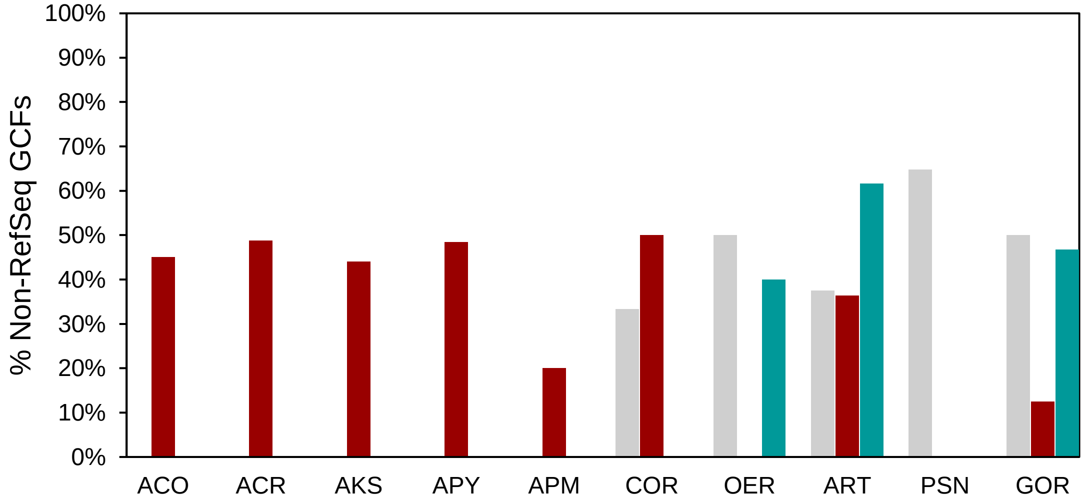

100%
90%
80%
70%
60%
50%
40%
30%
20%
10%
0%
% Non-RefSeq GCFs
ACO
ACR
AKS
APY
APM
COR
OER
ART
PSN
GOR

D

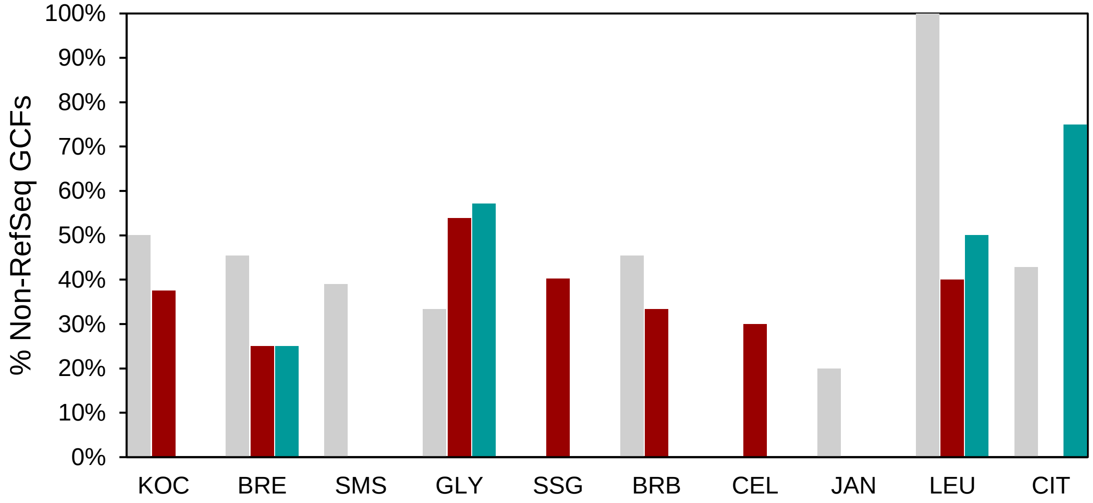
100%
90%
80%
70%
60%
50%
40%
30%
20%
10%
0%
% Non-RefSeq GCFs
KOC
BRE
SMS
GLY
SSG
BRB
CEL
JAN
LEU
CIT

E

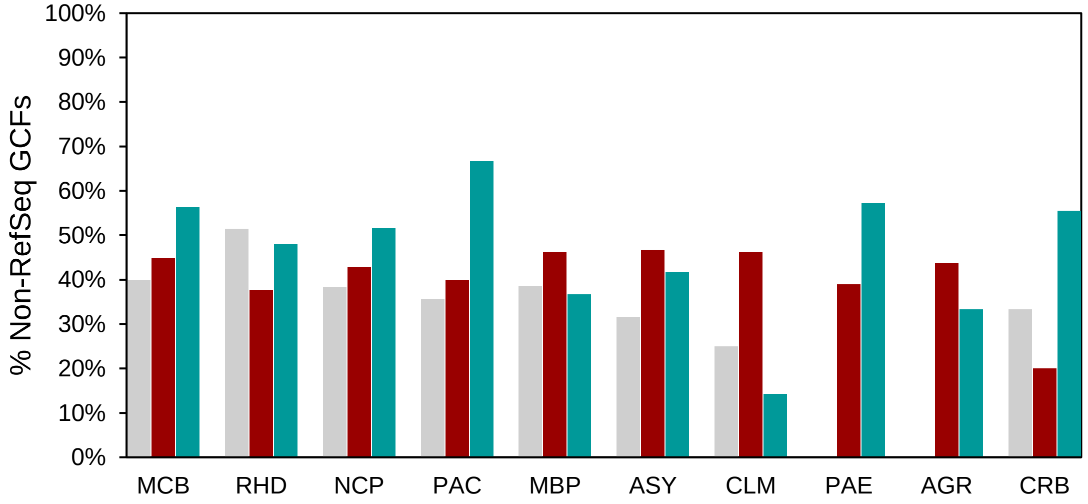
100%
90%
80%
70%
60%
50%
40%
30%
20%
10%
0%
% Non-RefSeq GCFs
MCB
RHD
NCP
PAC
MBP
ASY
CLM
PAE
AGR
CRB

F

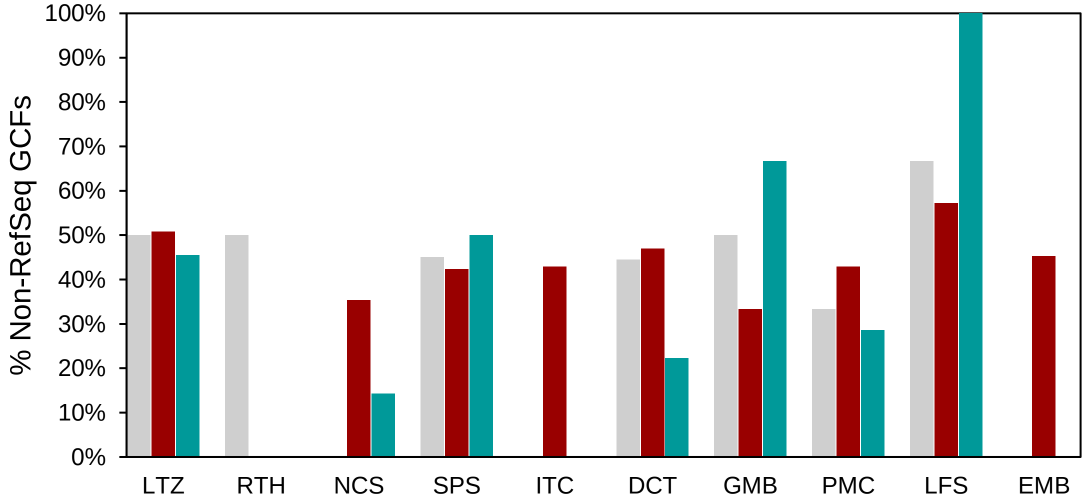
100%
90%
80%
70%
60%
50%
40%
30%
20%
10%
0%
% Non-RefSeq GCFs
LTZ
RTH
NCS
SPS
ITC
DCT
GMB
PMC
LFS
EMB

G

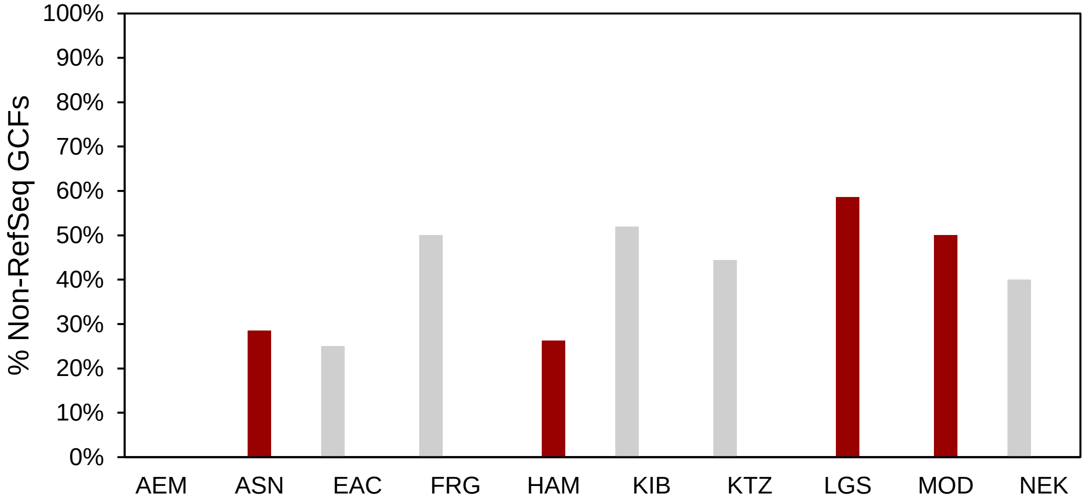

H

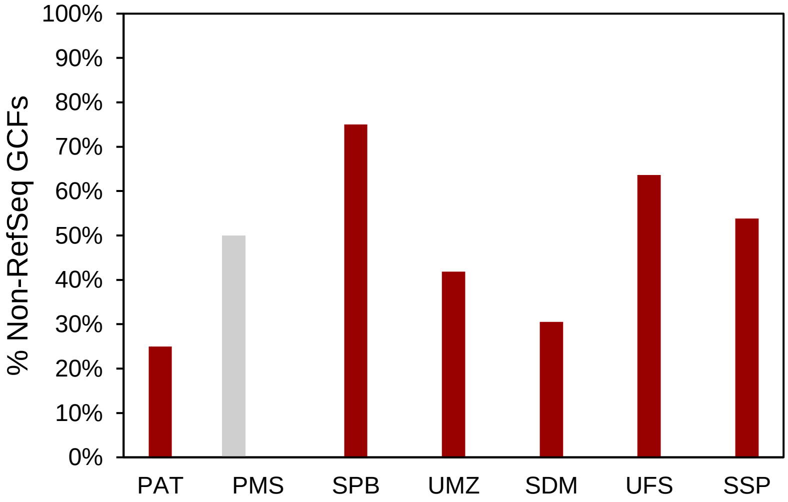

**Supplementary Fig. 18.** Percentages of non-RefSeq NPDC GCFs in RefSeq species (grey), new species (red), and shared between them (teal) across NPDC genera. Also see Fig. 4C. See Supplementary Table 10 for abbreviations.

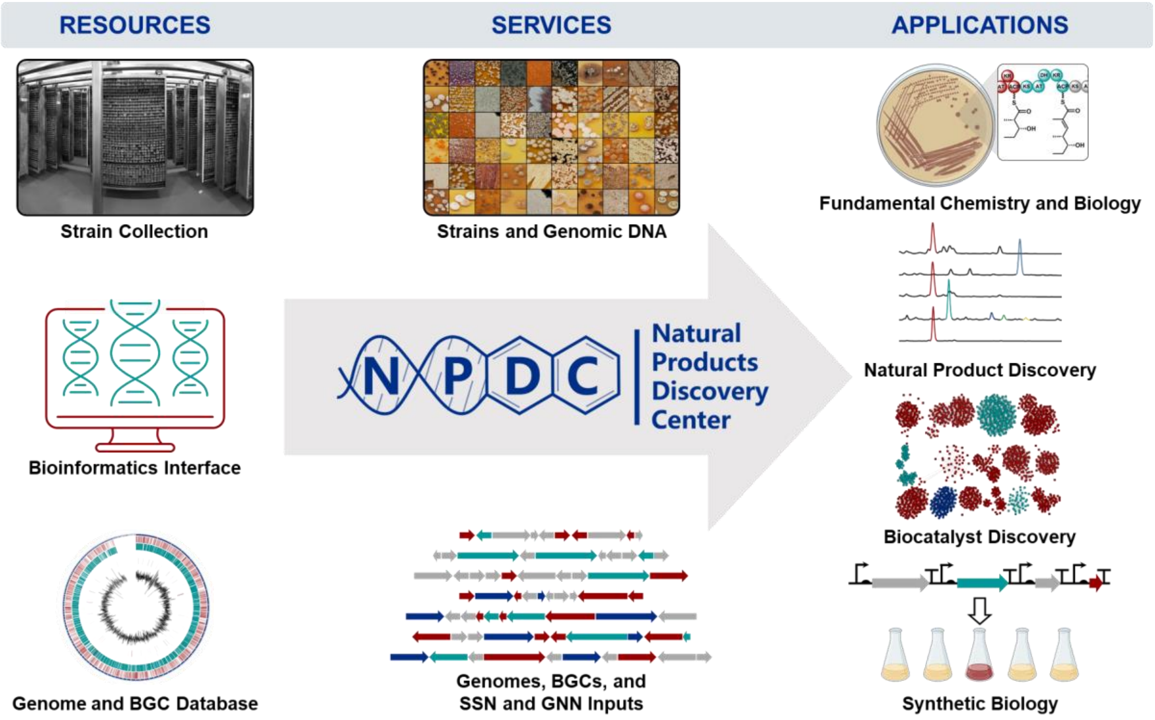

**Supplementary Fig. 19.** The NPDC Portal enables NP training, research, and associated applications. The NPDC provides a strain collection containing more than 122,000 bacterial strains, a database with more than 8000 genomes and 220,000 BGCs, and an online portal with BGC-centric bioinformatics tools for exploring the collection. Strains can be requested, and genomes, BGCs, and SSN/GNN input files can be downloaded. The NPDC is expected to have a broad, multi-disciplinary impact.

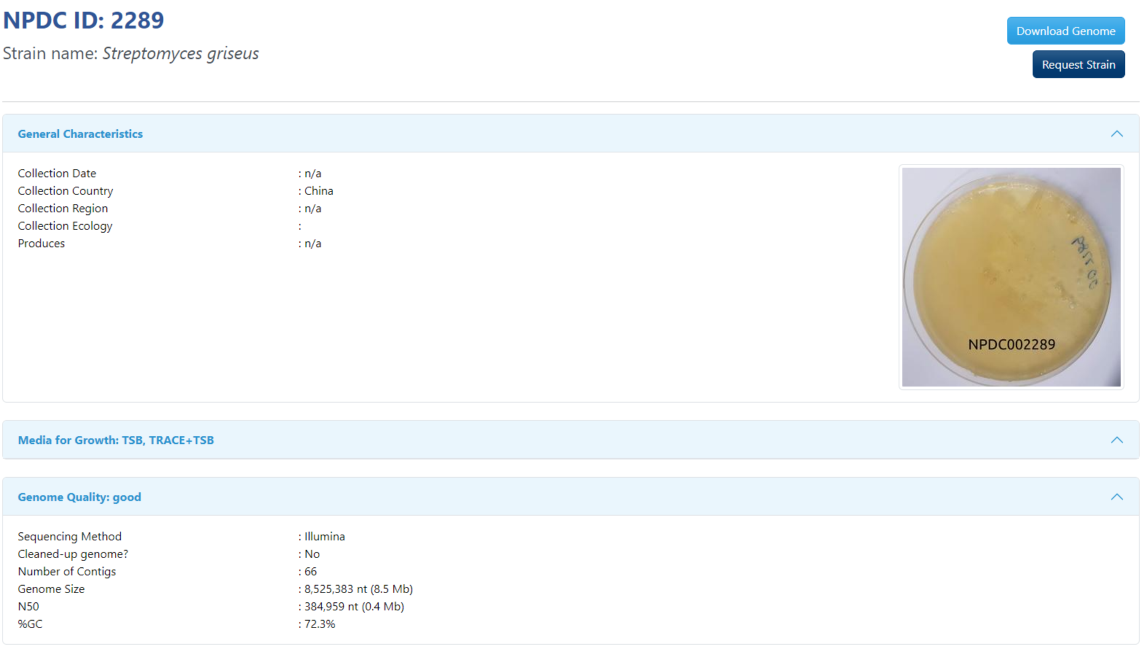

**Supplementary Fig. 20.** NPDC Portal strain page. For each strain, any associated metadata can be found on its individual webpage. If the strain has been sequenced, genome assembly statistics, BGCs, and related genomes are provided.

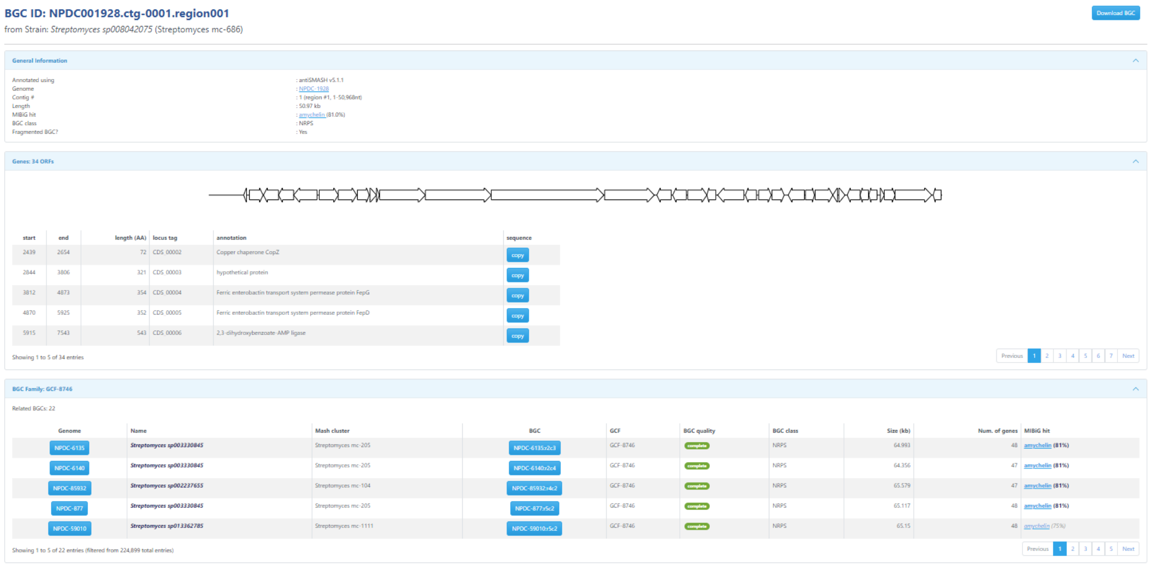

**Supplementary Fig. 21.** NPDC Portal BGC page. Each BGC has a dedicated page from which the antiSMASH-generated .gbk file can be downloaded. Further, the genes within the BGC can be viewed directly in the browser, and links to related BGCs at the bottom of each page are provided.

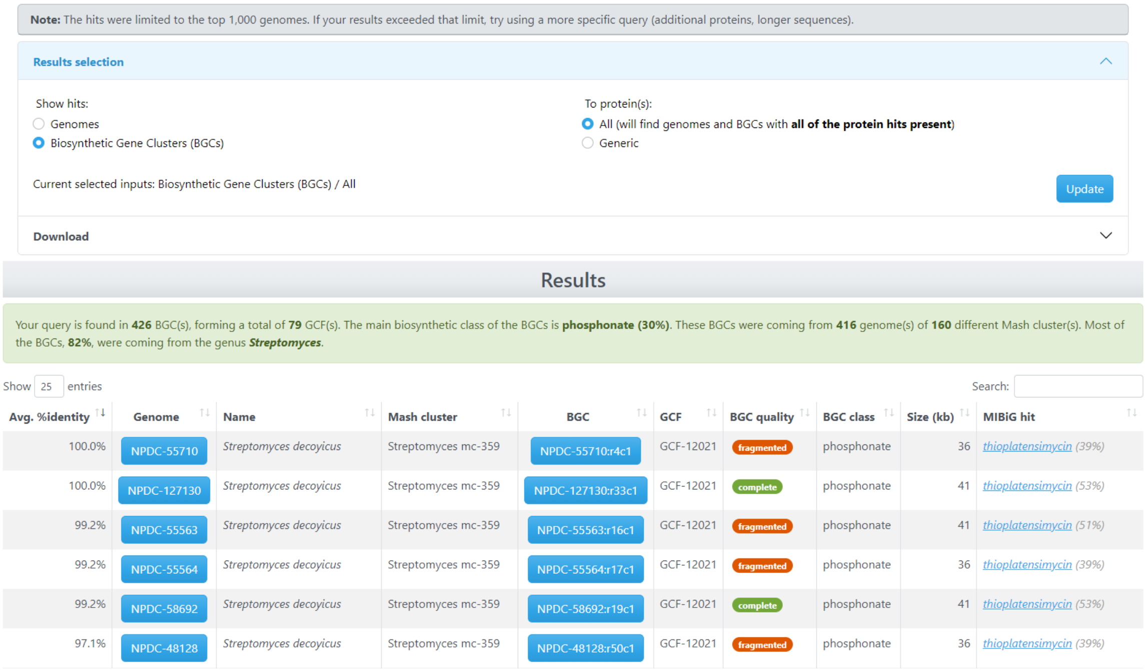

**Note:** The hits were limited to the top 1,000 genomes. If your results exceeded that limit, try using a more specific query (additional proteins, longer sequences).

**Results selection**

Show hits:
- Genomes
- Biosynthetic Gene Clusters (BGCs)

To protein(s):
- All (will find genomes and BGCs with **all of the protein hits present**)
- Generic

Current selected inputs: Biosynthetic Gene Clusters (BGCs) / All

Update

**Download**

Results

Your query is found in **426** BGC(s), forming a total of **79** GCF(s). The main biosynthetic class of the BGCs is **phosphonate (30%)**. These BGCs were coming from **416** genome(s) of **160** different Mash cluster(s). Most of the BGCs, **82%**, were coming from the genus ***Streptomyces***.

Show 25 entries

Search:

| Avg. %identity | Genome | Name | Mash cluster | BGC | GCF | BGC quality | BGC class | Size (kb) | MIBiG hit |
|---|---|---|---|---|---|---|---|---|---|
| 100.0% | NPDC-55710 | *Streptomyces decoyicus* | Streptomyces mc-359 | NPDC-55710:r4c1 | GCF-12021 | fragmented | phosphonate | 36 | *thioplatensimycin (39%)* |
| 100.0% | NPDC-127130 | *Streptomyces decoyicus* | Streptomyces mc-359 | NPDC-127130:r33c1 | GCF-12021 | complete | phosphonate | 41 | *thioplatensimycin (53%)* |
| 99.2% | NPDC-55563 | *Streptomyces decoyicus* | Streptomyces mc-359 | NPDC-55563:r16c1 | GCF-12021 | fragmented | phosphonate | 41 | *thioplatensimycin (51%)* |
| 99.2% | NPDC-55564 | *Streptomyces decoyicus* | Streptomyces mc-359 | NPDC-55564:r17c1 | GCF-12021 | fragmented | phosphonate | 36 | *thioplatensimycin (39%)* |
| 99.2% | NPDC-58692 | *Streptomyces decoyicus* | Streptomyces mc-359 | NPDC-58692:r19c1 | GCF-12021 | complete | phosphonate | 41 | *thioplatensimycin (53%)* |
| 97.1% | NPDC-48128 | *Streptomyces decoyicus* | Streptomyces mc-359 | NPDC-48128:r50c1 | GCF-12021 | fragmented | phosphonate | 36 | *thioplatensimycin (39%)* |

**Supplementary Fig. 22.** NPDC Portal BLASTP page. Users can utilize the DIAMOND-BLASTP tool to query up to five simultaneous protein sequences in either the full NPDC genome database or limit their searches to the NPDC BGC database. The results are saved for each user, and results can be sorted based on identity, taxonomy, GCF, BGC quality, BGC class, BGC size, or relatedness to MIBiG hits. The results page directly links each hit with genomes and BGCs. Under the ‘Downloads’ section, users can download a standard BLAST tabular result, a multiFASTA file containing all protein hits, and a multiFASTA file containing all protein sequences from BGCs encoding hits.

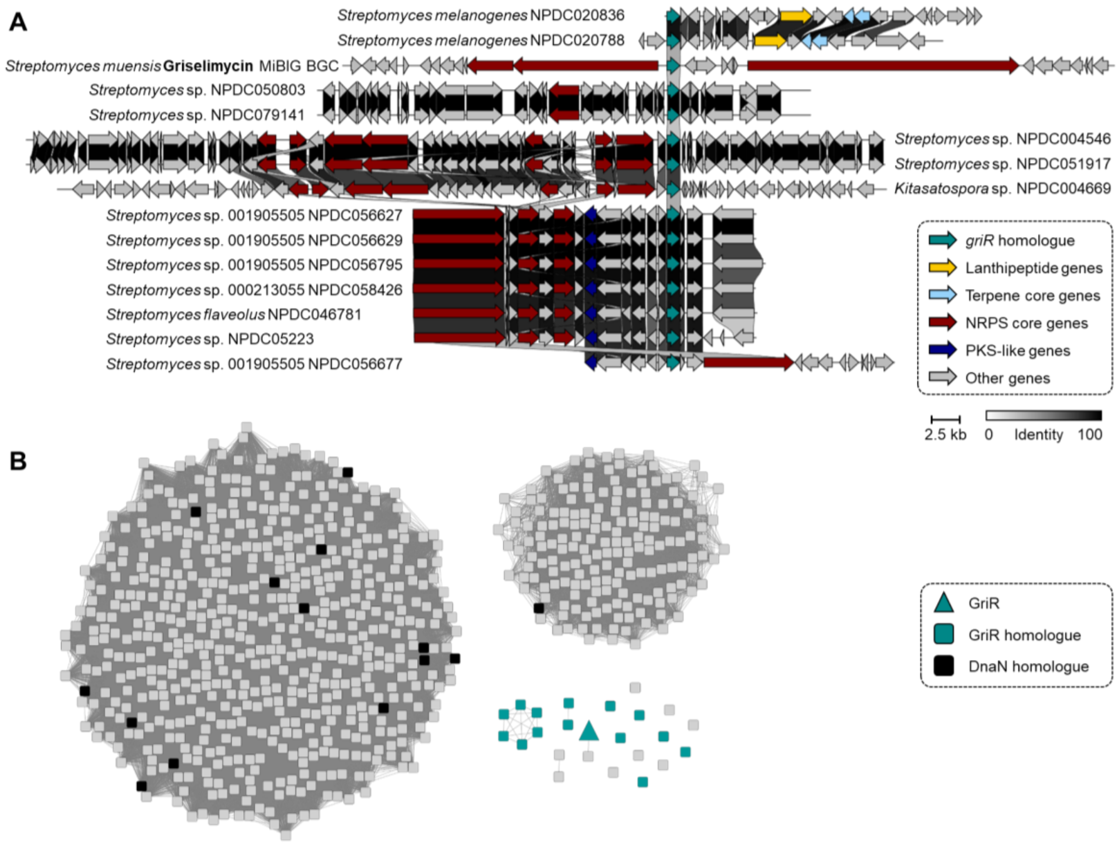

**Supplementary Fig. 23.** Genome mining for potential DnaN-targeting natural products. (A) *griR*-homologue containing BGCs identified in the NPDC with strain identifiers and taxonomy. (B) SSN of DnaN and GriR homologues (e-value $e^{-190}$) highlights the differences in protein sequence for distinguishing between primary metabolism proteins from potential resistance mediating copies. DnaN (black) and BGC-associated GriR (teal) homologues from strains highlighted in (A) are color-coded. Grey nodes represent DnaN sequences identified with the DIAMOND-BlastP search using the NPDC portal. Grey nodes not clustering with the two most well represented clusters potentially encode resistance-associated proteins but could not be associated with BGCs most often due to fragmentation of the genome in their vicinity.

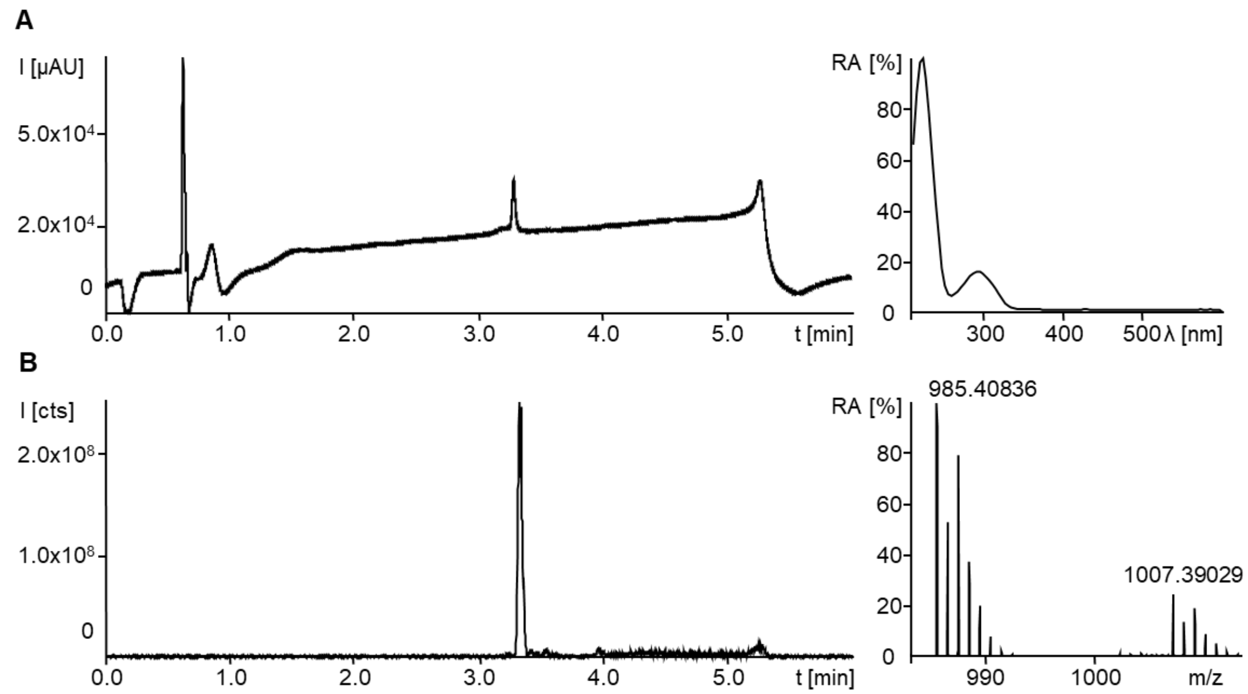

**Supplementary Fig. 24.** Chromatogram traces and spectra of bonnevillamide A purified from NPDC056627. (A) HPLC-UV/Vis chromatogram from 210-600 nm with corresponding UV/Vis spectrum at peak maximum. UV/Vis$_{max}$ are observed at 222 and 294 nm. (B) HPLC-MS base peak chromatogram from 300-1500 m/z with corresponding MS spectrum summed up over the peak. The two most abundant ions are the $[M+H]^+$ at 985.40836 ($C_{45}H_{67}O_{14}N_6Cl_2$ calc. 985.40868, Δ 0.33 ppm) and $[M+Na]^+$ at 1007.39029.

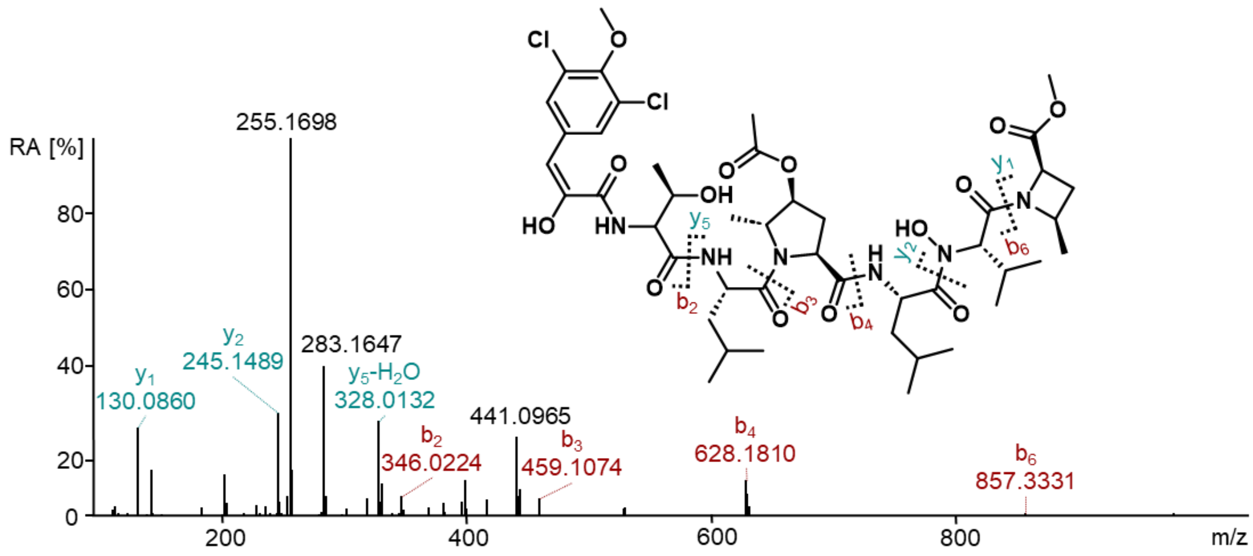

**Supplementary Fig. 25.** MS/MS spectra of bonnevillamide A purified from NPDC056627 with observed fragments of the y and b-series. The detected fragments match the ones described in literature. [2]

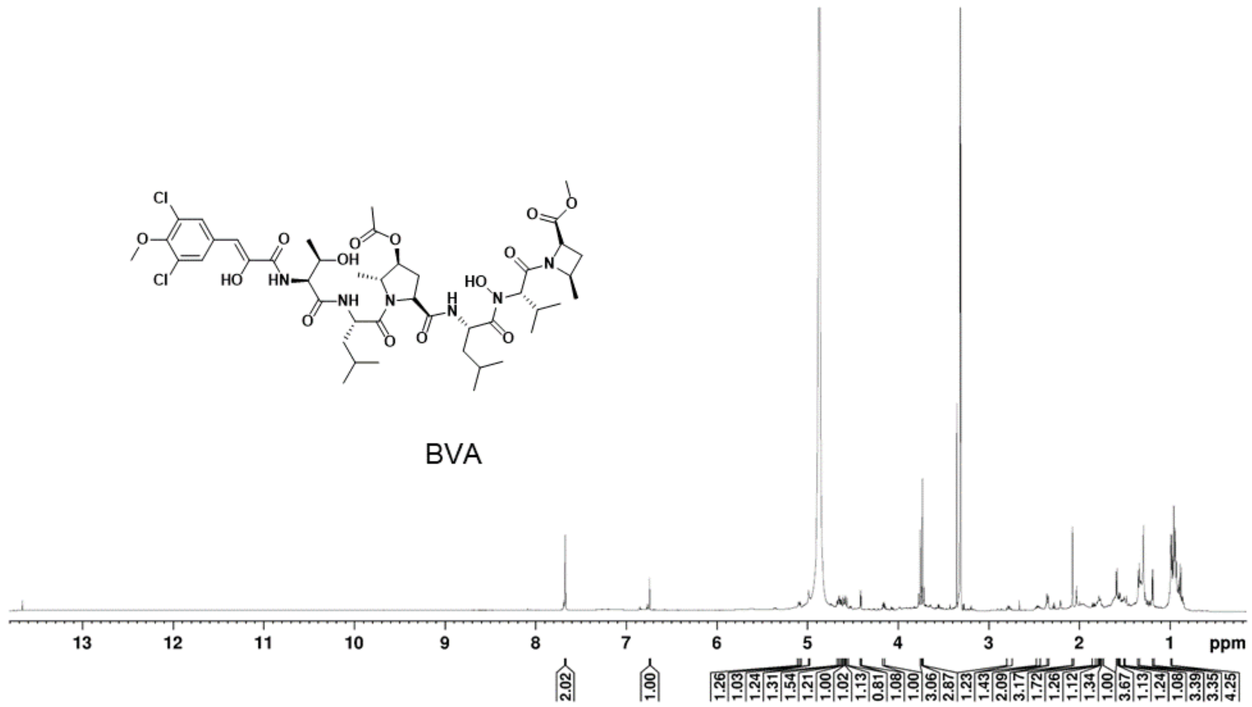

**Supplementary Fig. 26.** $^{1}$H spectrum of bonnevillamide A in MeOD-$d_4$ at 600 MHz.

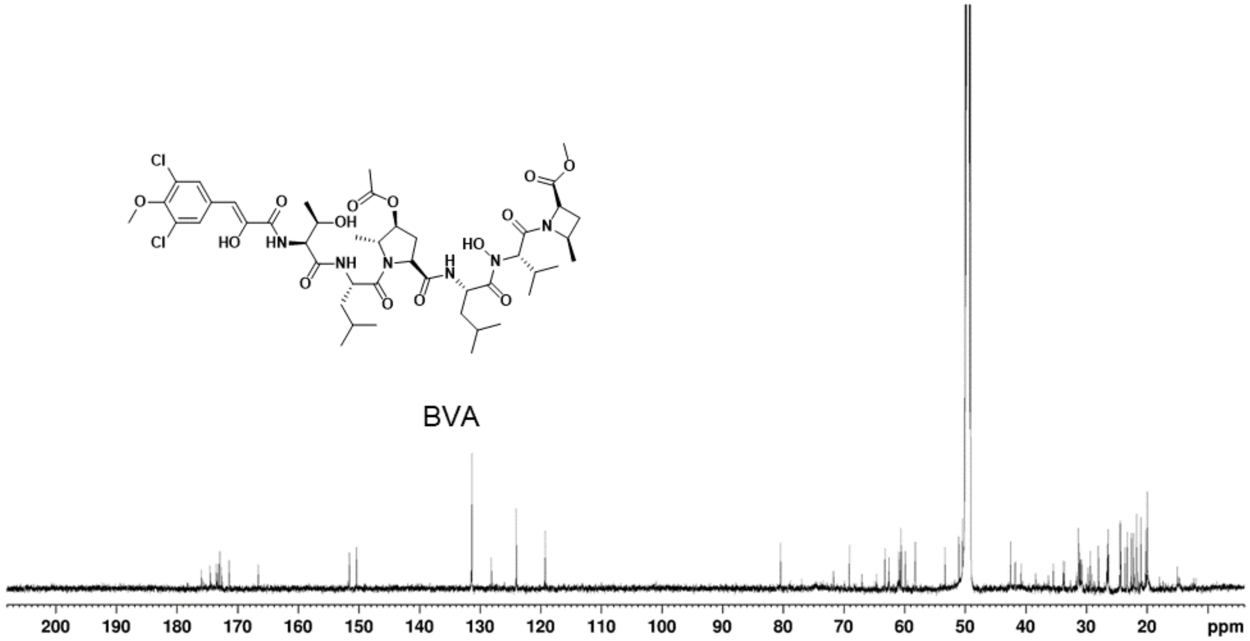

**Supplementary Fig. 27.** $^{13}C$ spectrum of bonnevillamide A in MeOD-$d_4$ at 151 MHz.

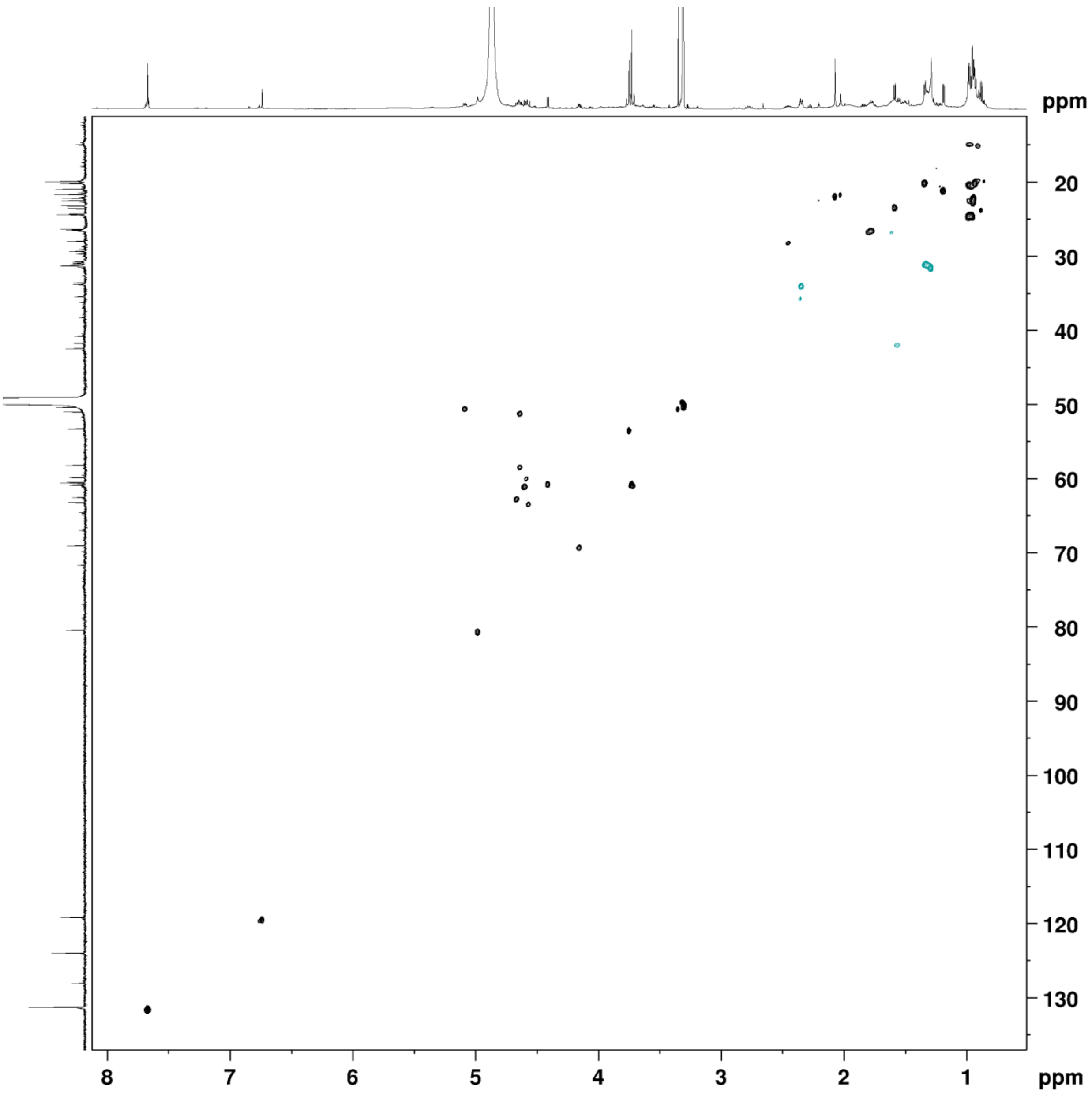

**Supplementary Fig. 28.** HSQC spectrum of bonnevillamide A in MeOD-$d_4$ at 600/151 MHz.

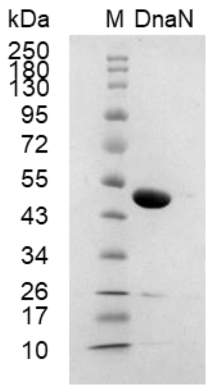

**Supplementary Fig. 29.** SDS-PAGE analysis of purified DnaN from *Micrococcus luteus* NPDC049463.

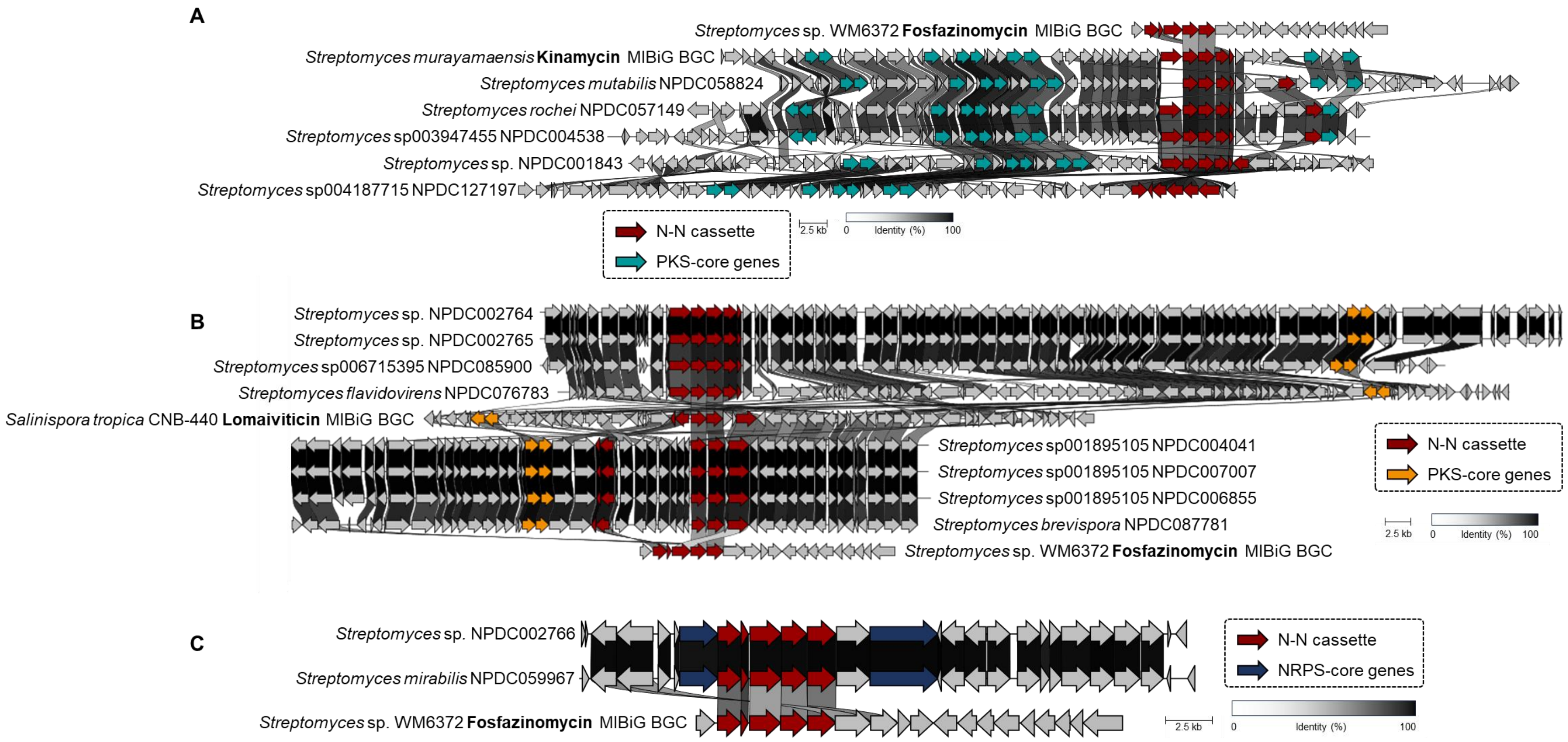

**Supplementary Fig. 30.** Clinker analysis of NPDC BGCs containing genes responsible for *N-N* bond formation. (A) BGCs related to the kinamycin MIBiG BGC are aligned. (B) BGCs related to the lomaiviticin MIBiG BGC are aligned. (C) BGCs in an unexplored GCF expected to encode for *N-N*-containing NPs are aligned.

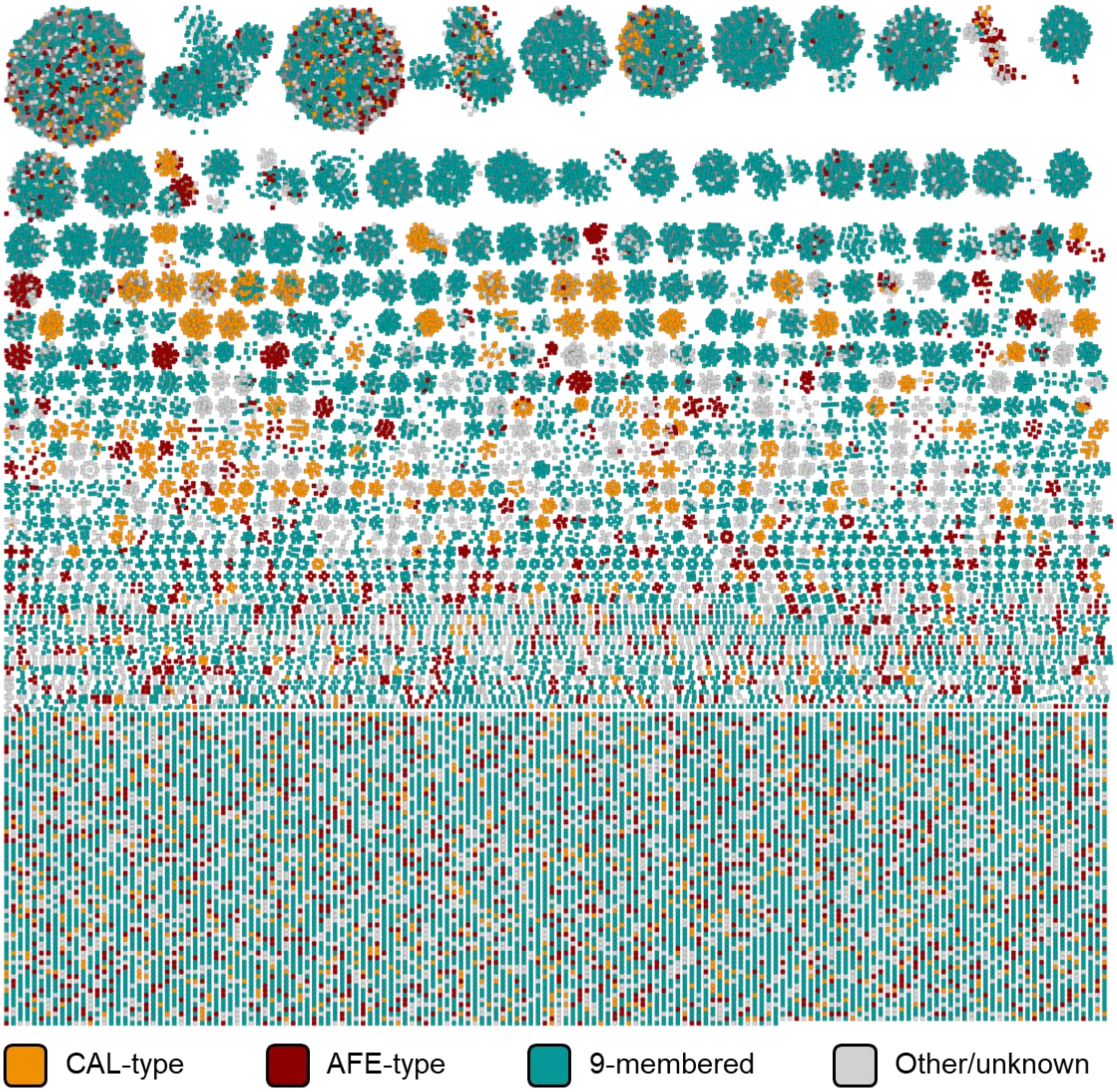

**Supplementary Fig. 31.** NPDC enediyne BGC-associated SSN colored by predicted enediyne type. Proteins encoded by NPDC enediyne BGCs mined from the NPDC BGC database were clustered using the EFI SSN tool (e-value e$^{-100}$) and color-coded by predicted enediyne subclass.[17] The protein sequences were obtained via the downloadable BGC multiFASTA file after querying the NPDC BGC database using TnmE4 and TnmE from tiancimycin A biosynthesis. CAL-type = calicheamicin-type enediynes; AFE-type = anthraquinone-fused enediynes; 9-membered = 9-membered enediynes. In this case, all proteins can be associated with BGCs expected to lead to NPs with the enediyne moiety, but the different enediyne-associated scaffolds require different—and difficult to predict—sets of biosynthetic enzymes.

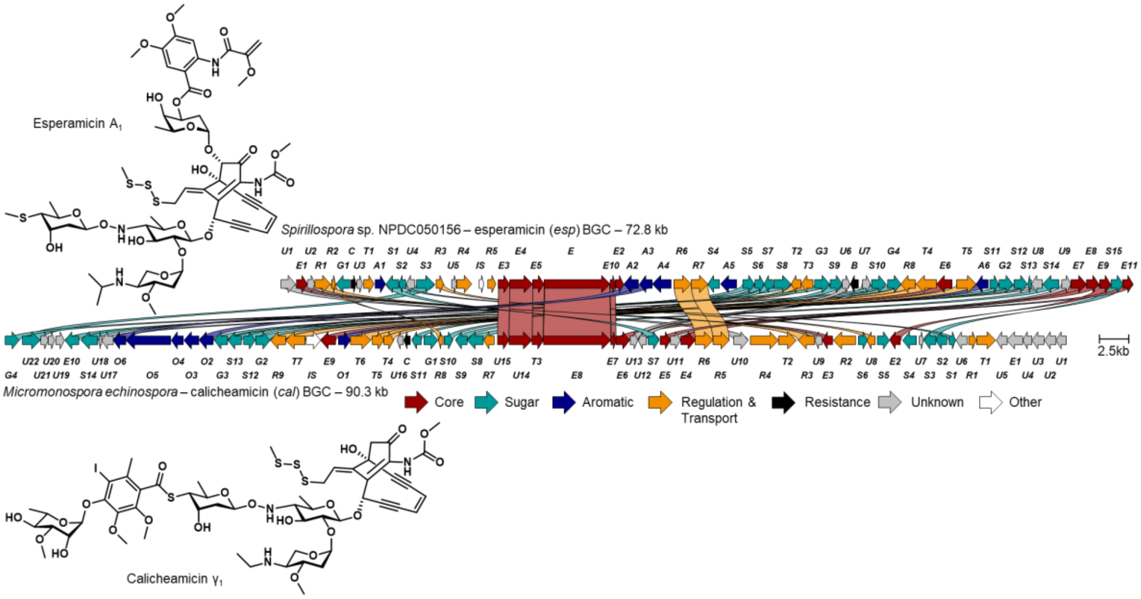

**Supplementary Fig. 32.** Comparison of the esperamicin and calicheamicin BGCs. There are substantial similarities both structurally and biosynthetically between the two enediyne NPs, as is highlighted through this color-coded Clinker analysis.[18]

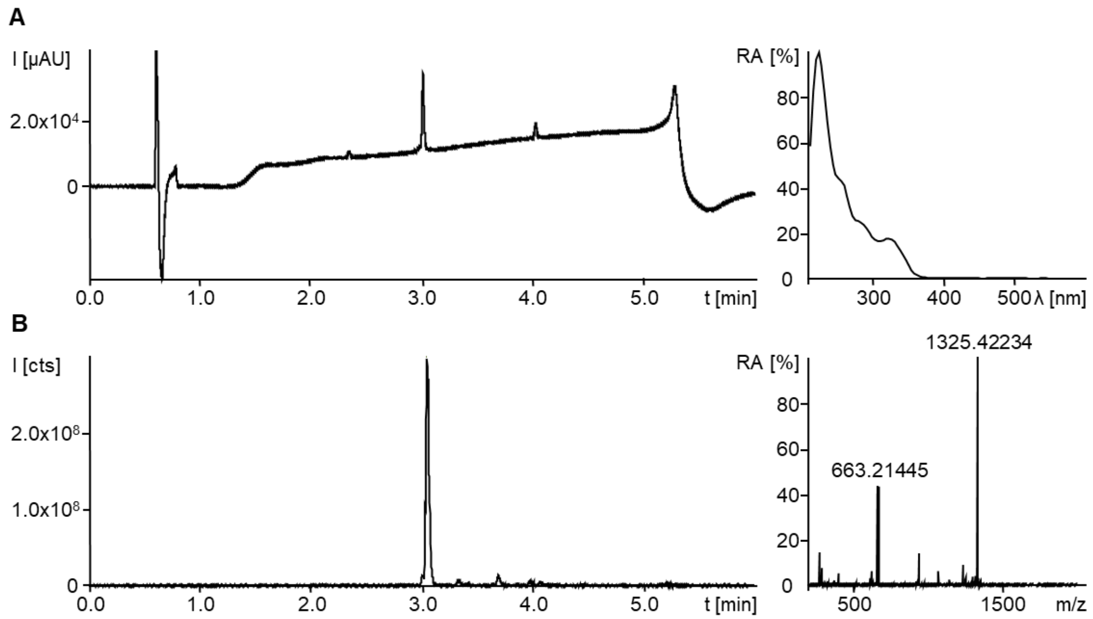

**Supplementary Fig. 33.** Chromatogram traces and spectra of esperamicin $A_1$ purified from NPDC048032. (A) HPLC-UV/Vis chromatogram from 210-600 nm with corresponding UV/Vis spectrum at peak maximum. $UV/Vis_{max}$ are observed at 222, 258, 278 and 322 nm. (B) HPLC-MS base peak chromatogram from 200-2000 m/z with corresponding MS spectrum summed up over the peak. The two most abundant ions are the $[M+H]^+$ at 1325.42234 ($C_{59}H_{81}O_{22}N_4S_4$ calc. 1325.42198, Δ 0.36 ppm) and $[M+2H]^{2+}$ at 663.21445.

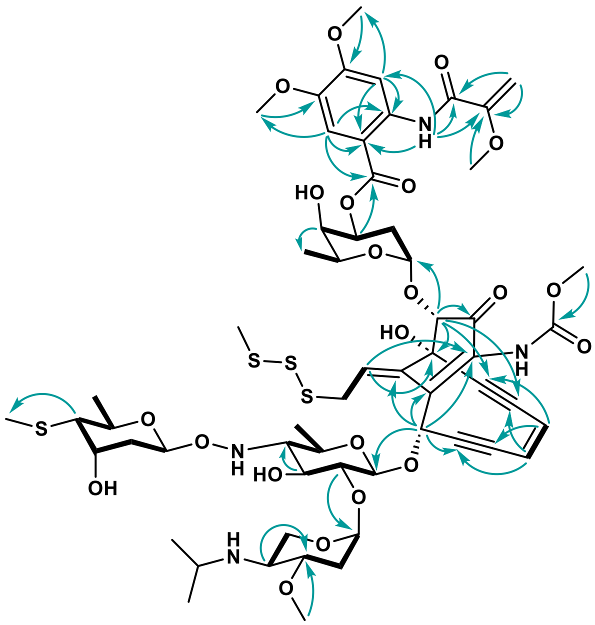

**Supplementary Fig. 34.** COSY-correlations and most relevant HMBC correlations in esperamicin $A_1$. COSY correlations are depicted as bolded bonds, and HMBC correlations are depicted as teal arrows.

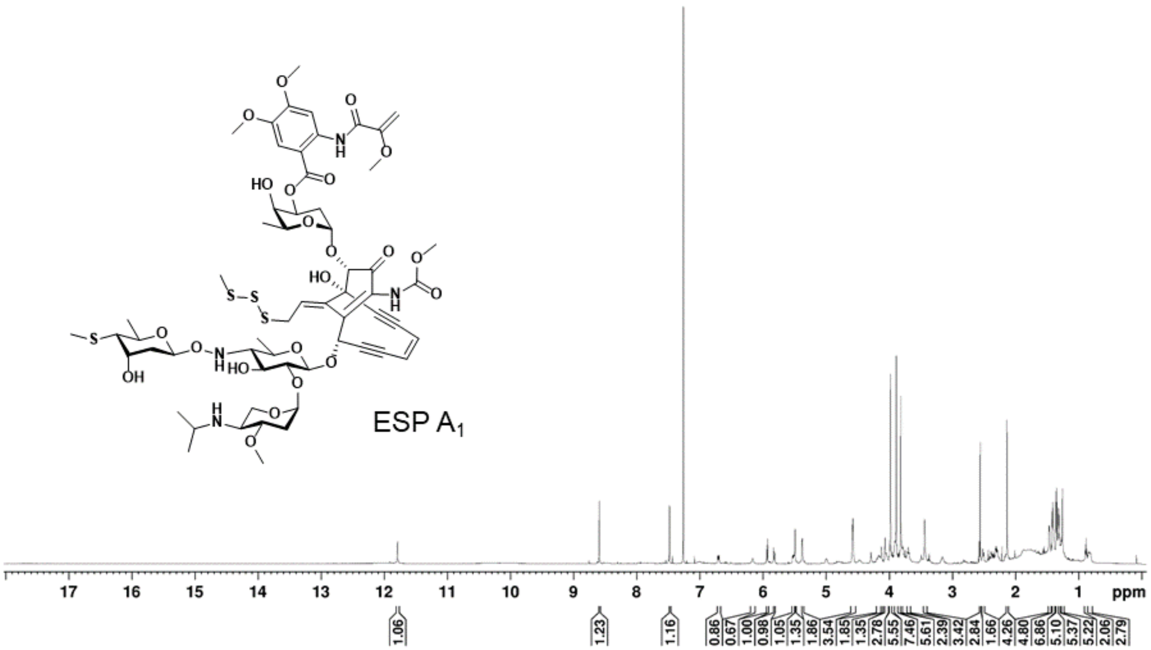

**Supplementary Fig. 35.** $^{1}H$ spectrum of esperamicin $A_1$ in $CDCl_3$ at 600 MHz.

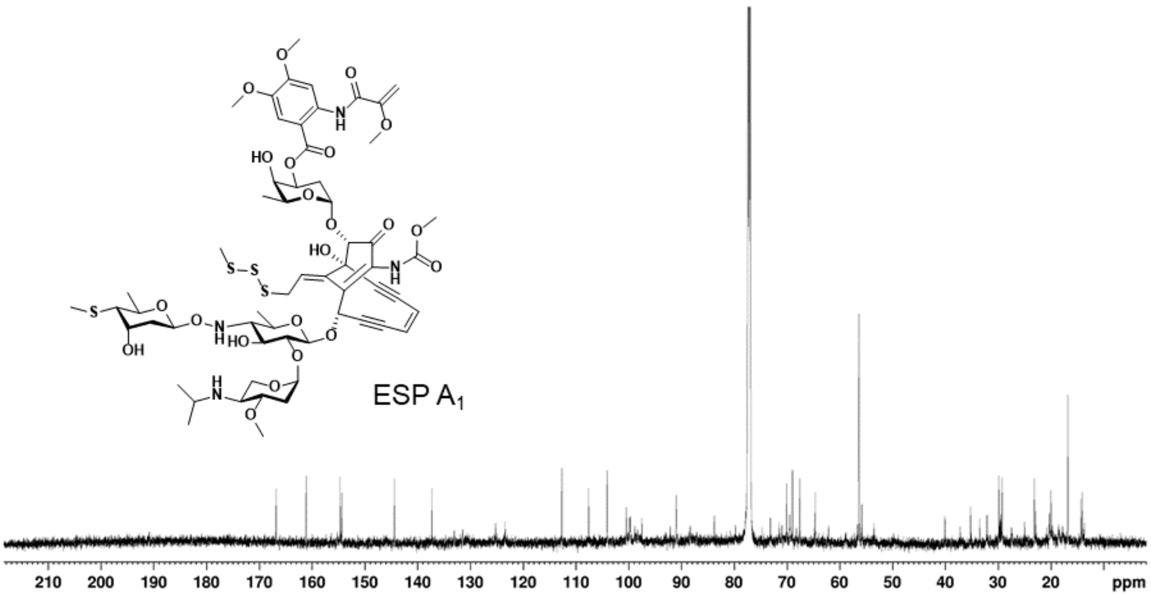

**Supplementary Fig. 36.** $^{13}C$ spectrum of esperamicin $A_1$ in $CDCl_3$ at 151 MHz.

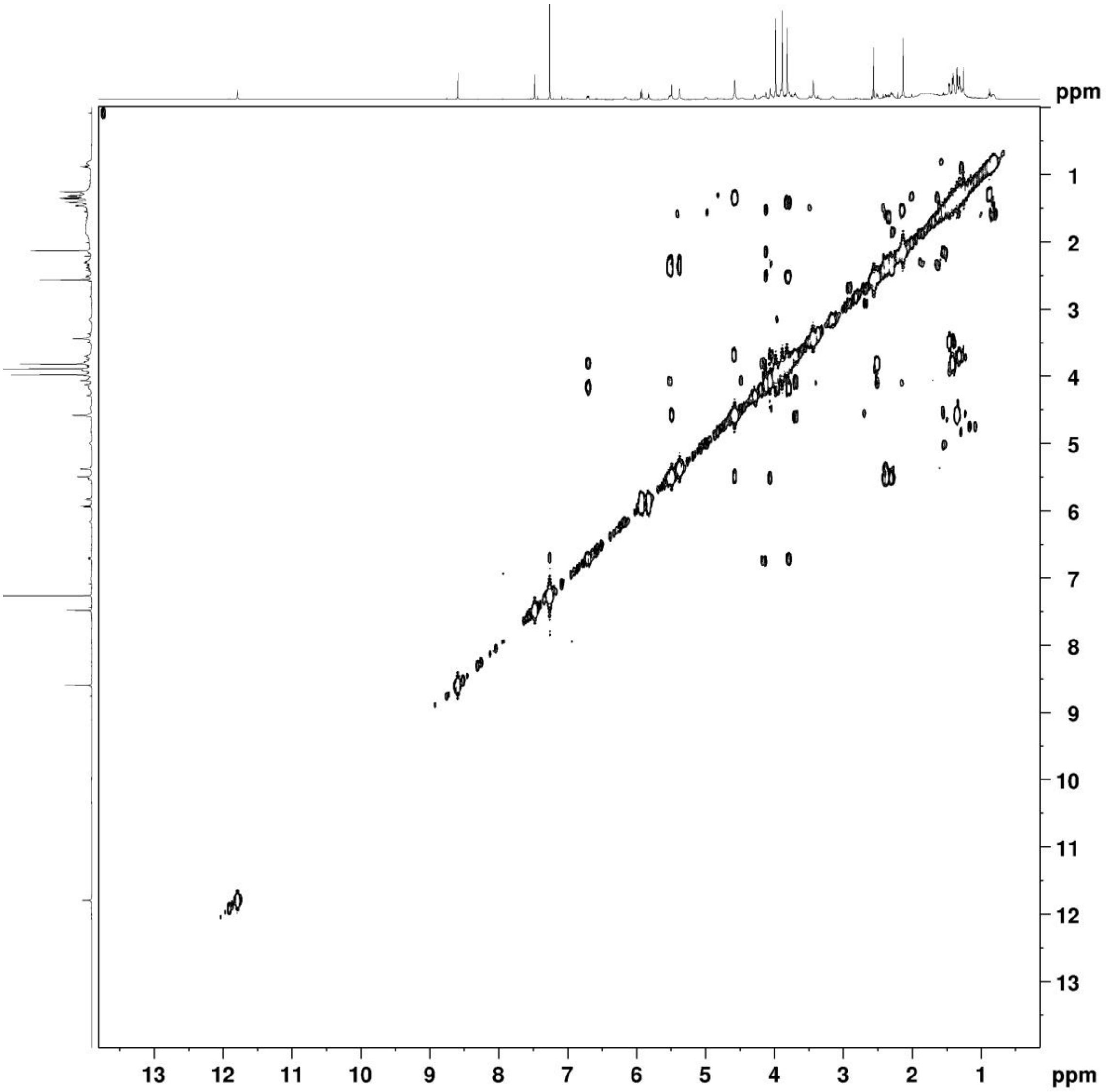

**Supplementary Fig. 37.** COSY spectrum of esperamicin A1 in $CDCl_3$ at 600 MHz.

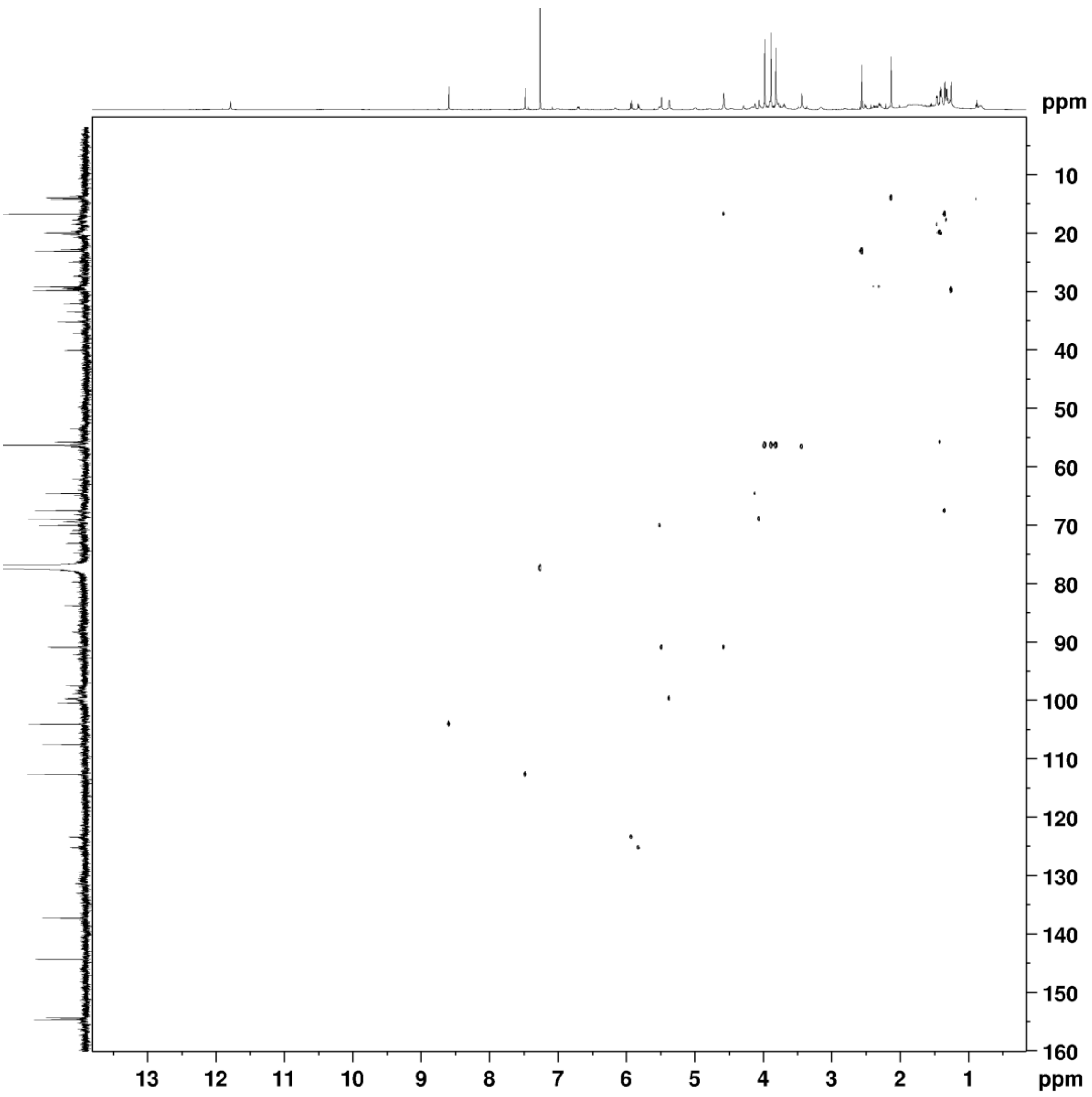

**Supplementary Fig. 38.** HSQC spectrum of esperamicin $A_1$ in $CDCl_3$ at 600/151 MHz.

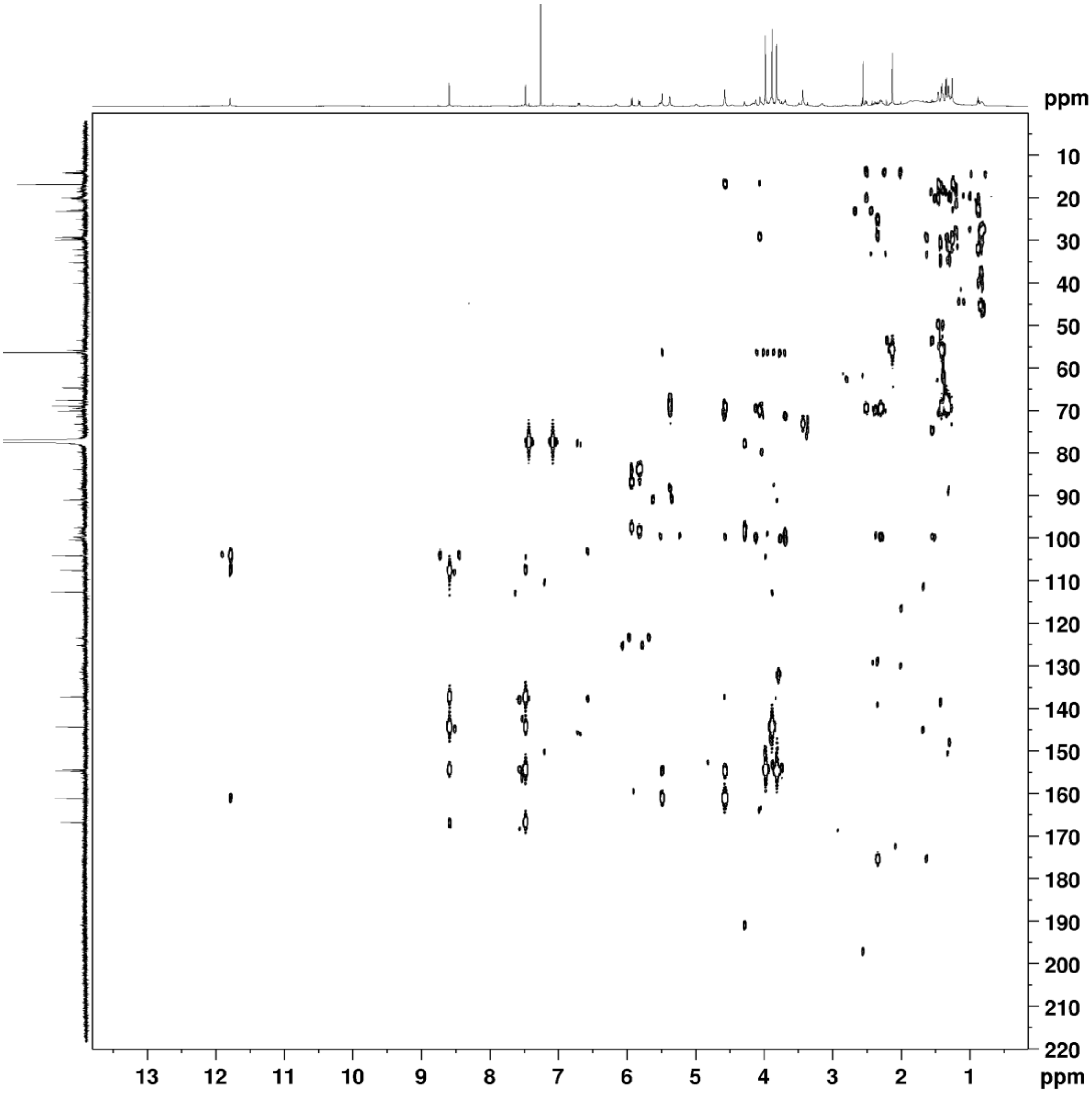

**Supplementary Fig. 39.** HMBC spectrum of esperamicin $A_1$ in $CDCl_3$ at 600/151 MHz.
